# Structural determinants of IGHV1-69 public antibodies conferring resilience to SARS-CoV-2 antigenic escape

**DOI:** 10.64898/2025.12.19.695307

**Authors:** Chuanying Niu, Xiaohan Huang, Qihong Yan, Banghui Liu, Xijie Gao, Yidong Song, Jingjing Wang, Longyu Wang, Zimu Li, Huiran Zheng, Ping He, Xiaodong Huang, Hang Yuan, Binqian Zou, Yuedong Yang, Fandi Wu, Yicheng Yao, Gul Habib, Xinwen Chen, Ling Chen, Jun He, Jianhua Yao, Jincun Zhao, Xiaoli Xiong

## Abstract

R1-32-like public antibodies, characterized by shared IGHV1-69/IGLV1-40 usage, are elicited in more than 50% of individuals with COVID-19 and have been implicated in driving recurrent mutations at L452_SARS2_ and F490_SARS2_ within their convergent epitope in the SARS-CoV-2 spike receptor-binding domain. These mutations effectively mediate escape from non-affinity-matured R1-32-like antibodies with germline-like sequences. Here, we characterize four affinity-matured R1-32-like antibodies, C092, C807, BD56-104, and BD56-597, that tolerate L452_SARS2_ and F490_SARS2_ mutations. We show that this tolerance arises from residues introduced by somatic hypermutation at convergent positions across multiple CDR loops and surrounding regions, thereby creating additional contacts that reinforce epitope binding. An unusual N354_SARS2_ glycosylation site, which emerged in BA.2.86 and became fixed in its descendants, is linked to escape from affinity-matured R1-32-like antibodies, implying ongoing selection by this public antibody class. Using an AI model trained on integrated structural, neutralization, and binding data, we further identified ZL525, an ultrapotent R1-32-like antibody with pan-SARS-CoV-2 variant activity, including against the highly evasive KP.3 variant carrying the N354_SARS2_ glycosylation, and broad sarbecovirus cross-reactivity extending to SARS-CoV-1. Together, these findings show how affinity maturation enables public antibodies to adapt to viral antigenic drift, reveal their role in shaping SARS-CoV-2 antigenic evolution, and demonstrate the potential of AI-empowered strategies for discovering broadly neutralizing antibodies.

## Introduction

The ongoing evolution of SARS-CoV-2, driven by immune pressure established from natural infections and vaccinations, has led to extensive amino acid substitutions in the spike (S) protein, the major viral surface glycoprotein and principal target of neutralizing antibodies ^1–3^. Several key residues in the receptor-binding domain (RBD), such as K417_SARS2_, L452_SARS2_, E484_SARS2_, F490_SARS2_, and N501_SARS2_, have become recurrent mutation hotspots across variants, enhancing immune evasion and viral fitness ^1,4–8^.

Based on their epitope locations, specifically whether they compete with ACE2 binding or require the RBD to adopt an ‘up’ conformation, SARS-CoV-2 RBD-targeting antibodies have been broadly classified into four epitope classes ^9^. In addition, many RBD-targeting antibodies isolated early in the pandemic exhibit germline-like sequences with low levels of somatic hypermutation ^10–12^. Some of these antibodies can be further grouped into distinct public (population) antibody classes, defined by convergent epitope recognition and shared genetic origins ^13–15^. Such population-level responses are thought to impose strong selective pressure on the virus and provide valuable insights into SARS-CoV-2 antigenic drift ^16–18^.

For example, many IGHV3-53/3-66-encoded public antibodies, which target the ACE2-binding site and are classified as epitope class 1, were effectively abrogated by the K417_SARS2_ mutation ^9,19,20^. Many IGHV1-2-encoded public antibodies, which recognize RBD in both ‘up’ and ‘down’ conformations and are classified as epitope class 2, are strongly evaded by E484_SARS2_ substitutions ^21,22^. Notably, we identified a public antibody class defined by the use of the IGLV6-57 light chain gene, which pairs with heavy chains of diverse genetic origins ^14^. These antibodies target a class 4 epitope relatively conserved among sarbecoviruses, encompassing RBD residues S371-S373-S375_SARS2_. Strikingly, we show that the featured Omicron S371L/F-S373P-S375F_SARS2_ mutations within this epitope mediate the escape of many IGLV6-57-class public antibodies ^14^. Other public antibody classes, including IGHV1-58/IGKV3-20 and IGHV2-5/IGLV2-14, display some breadth but remain variably susceptible to escape by mutations at F486_SARS2_ and positions 444-445_SARS2_, respectively ^23–26^.

We have also previously identified a class of R1-32-like public antibodies elicited in more than 50% of COVID-19 convalescents early in the pandemic. These antibodies are characterized by shared usage of the IGHV1-69/IGLV1-40 gene pairing and rely on the unique features of the IGHV1-69 gene, whose germline-encoded hydrophobic HCDR2 residues bind the hydrophobic RBD residues L452_SARS2_ and F490_SARS2_. This specific interaction renders them vulnerable to mutations at L452 and F490. ^16,27^. These residues have since emerged as mutation hot-spots for viral escape ^1,16,27^. While the Delta variant harbored the L452R_SARS2_ mutation, subsequent Omicron subvariants acquired L452R/Q/M_SARS2_ or F490S_SARS2_ mutations across multiple lineages (e.g., BA.4/5, BF.7, BQ.1.1, XBB, EG.5), reflecting sustained immune pressure at these positions ^28–30^.

Intriguingly, we found that several members of the R1-32-like public antibody class, including C092, C807, BD56-104, and BD56-597, retain robust binding to S-proteins of viral variants carrying mutations at positions 452_SARS2_ and 490_SARS2_. The mechanism by which these antibodies maintain cross-variant reactivity despite substitutions at key contact residues has remained unclear. In this study, we elucidate the structural and biochemical basis for their mutation tolerance. We show that somatic hypermutation introduced residues, following convergent patterns, confer resilience to antigenic epitope changes, including those located at L452_SARS2_ and F490_SARS2_, and resulting in broadened reactivity not only to SARS-CoV-2 variants but also to other SARS-related coronaviruses. Using our structural-functional relationship data, we trained an AI-based antibody discovery model ^31^, which led to the identification of ZL525, an elite R1-32-like antibody with ultrapotent, pan-variant neutralizing activity, capable of neutralizing the latest variants carrying a highly unusual epitope glycan mutation and extending cross-reactivity to SARS-CoV-1. Our findings illuminate the adaptability of public antibodies in response to the ongoing antigenic drift of SARS-CoV-2 and highlight the power of AI-driven approaches to identify broadly-neutralizing antibodies.

## Results

### Identification of R1-32-like public antibodies that tolerate spike mutations

We have previously defined IGHV1-69 encoded R1-32-like public antibodies as those that either exhibit a minimum of 80% amino acid similarity to R1-32 or FC08 in the HCDR3 region or contain a GYSGYG/D motif in HCDR3 ^16,27^. In our previous study, we identified and characterized 10 R1-32-like antibodies isolated from ancestral (or wildtype, WT) SARS-CoV-2 convalescents or vaccinees ^27,32–35^. Notably, among these, C092 and C807 showed no obvious reduction in binding to SARS-CoV-2 RBD variants with L452 or F490 mutations, including Delta (L452R_SARS2_), Kappa (L452R_SARS2_), and Lambda (L452Q_SARS2_+F490S_SARS2_) RBDs ^27^. We further identified BD56-104 and BD56-597 as mutation-resistant R1-32-like public antibodies that effectively neutralize Omicron subvariants BA.2.12.1 (L452Q_SARS2_), BA.2.13 (L452M_SARS2_), and BA.4/5 (L452R_SARS2_), through the analysis of published sequence and neutralization data ^28^ (**Fig. S1a**). Notably, both BD56-104 and BD56-597 were isolated from convalescents with Omicron BA.1 breakthrough infections ^28^. In contrast to R1-32, which has minimal somatic hypermutation (SHM), the 4 mutation-resistant antibodies, C092, C807, BD56-104, and BD56-597, carry 6-15 SHMs in their heavy chains and 3-4 SHMs in their light chains, indicative of substantial affinity maturation (**Fig.1a and b**).

To further characterize these R1-32-like public antibodies, we performed antigen binding assays for C092, C807, BD56-104, and BD56-597 against a more comprehensive and updated panel of SARS-CoV-2 RBDs, which we classified into 4 categories: 1) RBDs with single amino acid substitutions at key residues (K417_SARS2_, E484_SARS2_, T478_SARS2_, L452_SARS2_, F490_SARS2_); 2) RBDs from early VOC/VOI variants (Alpha to Lambda); 3) RBDs from early Omicron variants (BA.1 to BA.4/5); and 4) RBDs from later-emerging Omicron subvariants, including BQ, XBB, and EG variants. Within each category, mutants containing L452_SARS2_ and F490_SARS2_ mutations were further grouped together **(Fig. 1c and Fig. S1b-e)**.

**Fig. 1.**
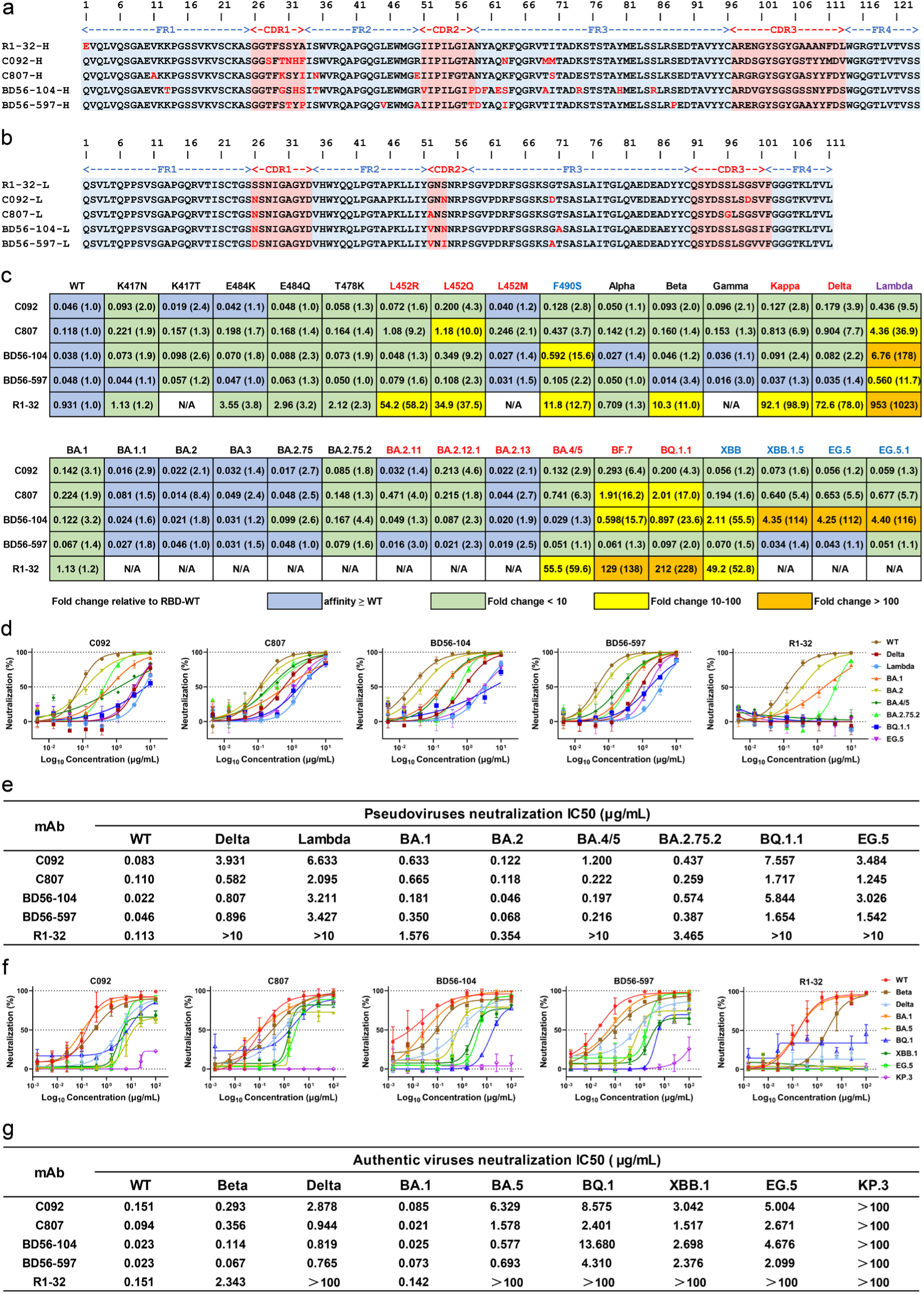
Sequence alignments, binding affinities, and neutralization activities of R1-32-like antibodies. **a**-**b**, Heavy chain (**a**) and Light chain (**b**) sequence alignments of germline-like R1-32 with selected affinity-matured R1-32-like antibodies (C092, C807, BD56-104, BD56-597). The sequences of R1-32 are used as the references, with somatic hypermutation-derived amino acids marked in red. The HCDRs and LCDRs residues are highlighted in salmon. **c**, Binding affinities of R1-32-like antibodies to a panel of SARS-CoV-2 RBDs. Binding assays were performed using biolayer interferometry (BLI). Dissociation constant *K*_D_s (nM) are shown, with fold changes relative to wild-type RBD indicated in parentheses. RBDs carrying L452, F490, or both substitutions are colored in red, blue, and purple, respectively. Binding affinities higher than the wild-type RBD are shown in blue, whereas reduced affinities (<10-fold, 10-100-fold, and >100-fold relative to wild-type) are shown in green, yellow, and orange, respectively. For R1-32, the *K*_D_s of Alpha, Beta, BA.1, K417N, E484K/Q, T478K, and L452Q were obtained from our previously published work ^27^. N/A means untested. RBD binding curves are shown in Fig. 4f **and Fig. S1b-e,** and *K*_D_ data used for comparisons are shown in **Tables S1 and S4**. **d**, Neutralization activities of R1-32-like antibodies towards SARS-CoV-2 wild-type, Delta, Lambda, Omicron BA.1, BA.2, BA.4/5, BA.2.75.2, and BQ.1.1 pseudoviruses. Data are presented as mean ± SD (n = 2). IC_50_ values are summarized in **e**. **f**, Neutralization activities of R1-32-like antibodies towards WT, Beta, Delta, Omicron BA.1, BA.5, BQ.1, XBB.1, EG.5, and KP.3 authentic viruses. Data are presented as mean ± SD (n = 2). IC_50_ values are summarized in **g.**

The results showed that C092, C807, BD56-104, and BD56-597 retained high-affinity binding (*K*_D_ < 1.2 nM) to SARS-CoV-2 RBDs carrying L452_SARS2_ and F490_SARS2_ single mutations. In contrast, the germline-like R1-32 exhibited substantially reduced binding to these mutant RBDs (*K*_D_ = 11.8-54.2 nM) **(Fig. 1c and Fig. S1b-e)**. Substitutions at K417_SARS2_, E484_SARS2_, and T478_SARS2_, commonly associated with the escape of epitope class 1 and class 2 antibodies ^9,20,21^, had minimal impact on the binding of C092, C807, BD56-104, and BD56-597 **(Fig. 1c and Fig. S1b-e)**, consistent with these residues being located at the periphery or outside of the epitope recognized by the R1-32 class of public antibodies.

C092, C807, BD56-104, and BD56-597 maintained very high affinities (*K*_D_ < 0.3 nM) towards RBDs of variants carrying no mutations at 452_SARS2_ or 490_SARS2_, including Alpha (B.1.1.7), Beta (B.1.351), Gamma (P.1), and multiple Omicron sublineages (BA.1, BA.1.1, BA.2, BA.3, BA.2.75, and BA.2.75.2). Consistent with their mutation tolerance, C092, C807, BD56-104, and BD56-597 also maintained high affinities (*K*_D_ < 2.1 nM) towards RBDs harboring L452_SARS2_ substitutions, including Kappa (B.1.617.1), Delta (B.1.617.2), Omicron BA.2.11, BA.2.12.1, BA.2.13, BA.4/5, BF.7, and BQ.1.1. For comparison, germline-like R1-32 exhibited substantially reduced binding to these mutants (*K*_D_ > 55 nM).

Notably, C092, C807, BD56-104, and BD56-597 maintained high affinities (*K*_D_ < 4.5 nM) towards RBDs of the later emerged XBB lineages (XBB, XBB.1.5), and EG.5 lineages (EG.5, EG.5.1), despite the presence of F490S_SARS2_ mutations in these variants. All four antibodies exhibited the weakest binding to the Lambda (C.37) RBD carrying both the L452_SARS2_ and F490_SARS2_ mutations, although their *K*_D_ values are still below 6.8 nM **(Fig. 1c and Fig. S1b-e)**. For comparison, the germline-like R1-32 binds Lambda RBD with a *K*_D_ of ∼ 1000 nM, representing a reduction of ∼1000-fold.

These RBD binding results confirm that BD56-104 and BD56-597, like C092 and C807, are tolerant to diverse variants carrying L452_SARS2_ and F490_SARS2_. Among them, we found that C092 and BD56-597 consistently exhibited more robust binding across RBD variants compared to C807 and BD56-104. C807 showed moderately but noticeably reduced binding to RBDs harboring L452_SARS2_ mutations, such as those in Kappa, Delta, Lambda, Omicron BA.4/5, BF.7, and BQ.1.1, characterized by accelerated dissociation kinetics (**Fig. S1c**). BD56-104 also showed moderate but noticeable sensitivity to F490_SARS2_ substitutions, displaying accelerated dissociation from RBDs of Lambda, XBB lineages (XBB, XBB.1.5), and EG.5 lineages (EG.5, EG.5.1) that carry this mutation (**Fig. S1d**).

### Mutation-tolerant antibodies retain potent neutralization against diverse SARS-CoV-2 variants

Neutralization activities of C092, C807, BD56-104, and BD56-597 were evaluated using VSV-based pseudovirus neutralization assays ^36^ against eight variants of SARS-CoV-2: the ancestral SARS-CoV-2 strain (WT), Delta (B.1.617.2), Lambda (C.37), Omicron BA.1 (B.1.1.529), BA.2, BA.4/5, BA. 2.75.2, BQ.1.1, and EG.5. The results showed that all four antibodies neutralized all tested variants, with IC_50_ values ranging from 0.022 to 7.557 µg/mL **(Fig. 1d and e)**. However, compared with the WT, obvious reductions in neutralization activity were observed for Delta (L452R_SARS2_), Lambda (L452Q_SARS2_+F490S_SARS2_), BA.4/5 (L452R_SARS2_), BQ.1.1 (L452R_SARS2_), and EG.5 (F490S_SARS2_) variants containing L452_SARS2_, F490_SARS2,_ or both mutations. By contrast, germline-like R1-32 lost neutralization (IC_50_ >10 µg/mL) against Delta, Lambda, BA.4/5, BQ.1.1, and EG.5 pseudoviruses.

Consistent with pseudovirus neutralization assays, C092, C807, BD56-104, and BD56-597 demonstrated potent neutralization (most IC_50_ <10 µg/mL) against authentic SARS-CoV-2 variants, including Delta (L452R_SARS2_), BA.5 (L452R_SARS2_), BQ.1(L452R_SARS2_), XBB.1(F490S_SARS2_), and EG.5 (F490S_SARS2_), while R1-32 showed no detectable neutralization (IC_50_ >100 µg/mL) against these variants **(Fig. 1f and g)**. However, all five antibodies lost neutralizing activities against the recent circulating variant KP.3. Complementing the neutralization data, C092, C807, BD56-104, and BD56-597 exhibited strong, non-dissociating binding to the S-trimers of variants neutralized in the neutralization assays **(Fig. S2)**. Binding affinity assays revealed moderately weakened binding with Lambda (L452Q_SARS2_+F490S_SARS2_), BA.4/5 (L452R_SARS2_), and BQ.1.1 (L452R_SARS2_) S-trimers relative to WT, consistent with the observed reduction in neutralization potency against these variants. Altogether, our neutralization and binding data demonstrate that C092, C807, BD56-104, and BD56-597 exhibit markedly enhanced breadth and mutation tolerance compared to the germline-like R1-32, maintaining neutralization activity against SARS-CoV-2 variants carrying L452_SARS2_, F490_SARS2,_ or even both escape mutations, with minimal to modest reductions in potency.

### Binding of R1-32-like antibodies disassembles S-trimer without blocking ACE2 binding

To elucidate how C092, C807, BD56-104, and BD56-597 tolerate mutations to cross-neutralize SARS-CoV-2 variants, we prepared cryo-electron microscopy (cryo-EM) samples using stoichiometric mixtures of Fab, ACE2, and BA.4/5 S-trimer (Fab:ACE2:S-protomer = 1:1:1). Cryo-EM structures of C092, C807, BD56-104, and BD56-597 Fabs bound to BA.4/5 S-trimer in complex with ACE2 were determined (**Fig. 2a and Figs. S5-9**).

**Fig. 2.**
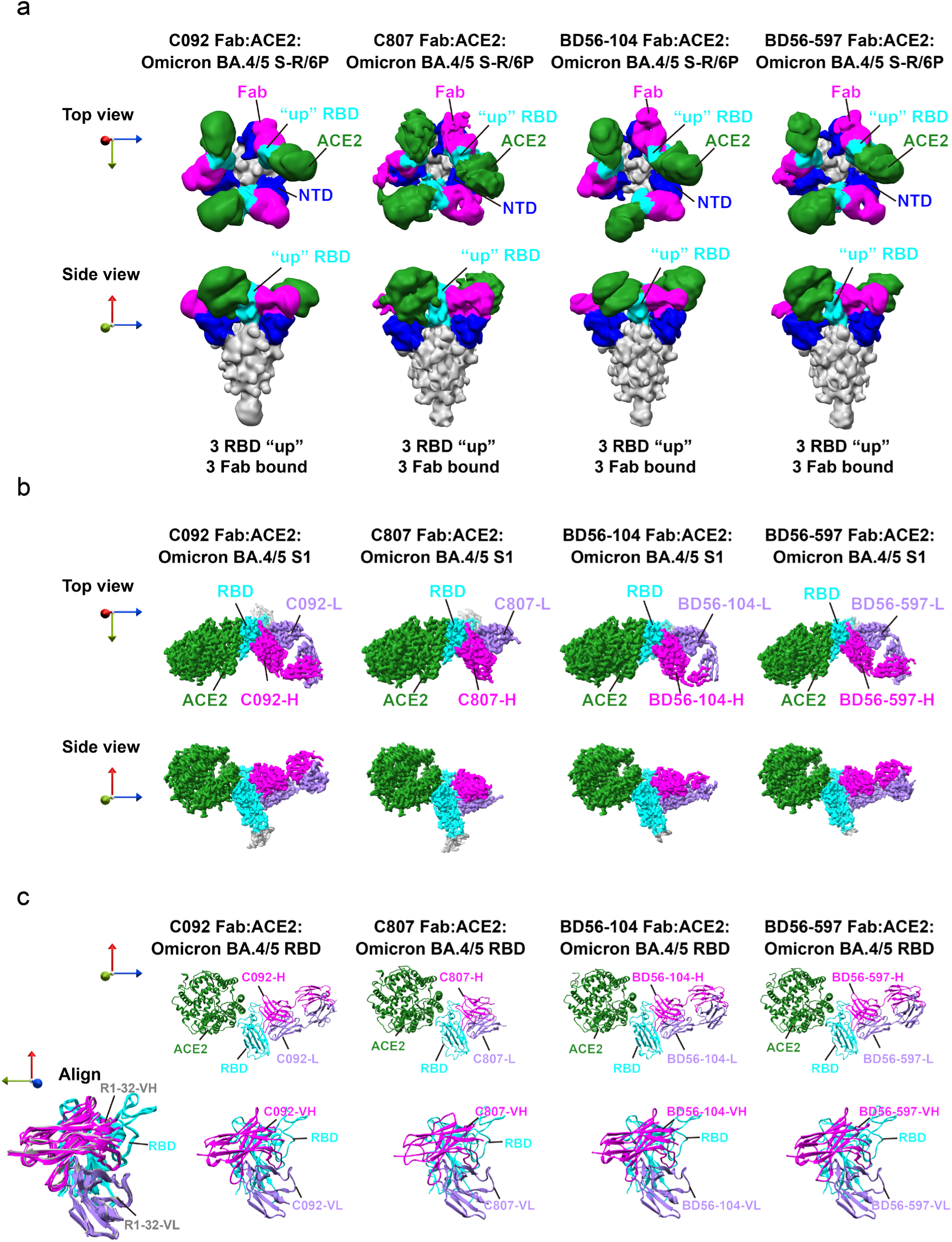
Structures of mutation-tolerant antibodies in complex with SARS-CoV-2 BA.4/5 S-trimer and ACE2. **a**, Cryo-EM densities (low-pass filtered to 12 Å) of BA.4/5 S-R/6P S-trimer bound to Fab and ACE2 at 3:3:3 molar ratio. C092, C807, BD56-104, and BD56-597 Fabs, NTD, RBD, and ACE2 are colored in magenta, blue, cyan, and dark green, respectively; the rest of the S-trimer is colored in grey. The 3 RBDs in “up” positions are indicated, each simultaneously engaging both ACE2 and antibody. **b**, Cryo-EM densities of BA.4/5 S1 bound to Fab and ACE2. C092, C807, BD56-104, and BD56-597 Fab-HC, Fab-LC, NTD, RBD, and ACE2 are colored in magenta, purple, blue, cyan, and dark green, respectively; the rest of the S1 is colored in grey. **c**, Structures of the BA.4/5 RBD:Fab:ACE2 complex (**upper panels**) derived from the density shown in panel **b**. Individual binding modes of C092, C807, BD56-104, and BD56-597 to the RBD are shown in the lower panels. Superposition of C092, C807, BD56-104, BD56-597, and R1-32 bound to RBD is shown in the left panels, with C092, C807, BD56-104, and BD56-597 Fabs rendered in color and the R1-32 Fab shown in grey.

For the C092 Fab:ACE2:S-trimer sample, each S-trimer was found to adopt either 2 or 3 RBD “up” conformations, allowing simultaneous binding of either 2 or 3 Fab and ACE2 molecules. All other structures showed S-trimers with all three RBDs in the “up” position, with each RBD concurrently engaging both ACE2 and Fab **(Figs. S5-8)**. Consistent with previous conclusions ^27^, these S-trimer structures confirm that R1-32-like public antibodies do not interfere with ACE2 binding, in agreement with results from the antibody-ACE2 competition assays **(Fig. S3)**. In addition to ligand bound S-trimer structures, we also observed isolated S1 domains bound to both Fab and ACE2, suggesting that a portion of S-trimers disassembles upon simultaneous antibody and ACE2 binding. **(Figs. S5-8)**. Based on these disassembled particles, Structures of C092, C807, BD56-104, and BD56-597 bound to ACE2 bound S1 domain were determined to high-resolutions (2.58 - 2.83 Å) **(Fig. 2b and Figs. S5-9)**. We previously concluded that R1-32 neutralizes SARS-CoV-2 by inducing disassembly of S-trimers, thereby rendering them incapable of mediating membrane fusion ^27^. Based on the imaging data, we infer that C092, C807, BD56-104, and BD56-597 similarly retain the ability to induce disassembly of S-trimers from variant viruses.

### R1-32-like antibodies impair spike fusogenic conformational change

To further confirm the neutralization mechanism, we conducted ligand-induced conformational change assays. The results showed that, like the WT, BA.4/5 S-trimers can be triggered by ACE2-Fc or YB9-258 ^37^, an epitope class 1 antibody that can mimic ACE2 function, to undergo the fusogenic conformational change, generating the proteinase K-resistant S2 fusion core **(Fig. S4a and c)**. ACE2-Fc retains the ability to trigger the EG.5.1 S-trimer, whereas YB9-258 loses this ability, likely due to epitope mutations in EG.5.1 that disrupt YB9-258 binding **(Fig. S4e)**. In line with our previous findings, S-trimers (WT, BA.4/5, and EG.5.1) pre-incubated with R1-32 and the four antibodies (C092, C807, BD56-104, and BD56-597) were unable to undergo structural transition into a post-fusion conformation **(Fig. S4a, c, and e).**

Further, assay results also showed that WT, BA.4/5, and EG.5 S-trimers pre-incubated with the four R1-32-like antibodies (C092, C807, BD56-104, or BD56-597) were unable to undergo the fusogenic conformational change upon subsequent addition of ACE2. **(Fig. S4b, d, and f)**. In contrast, although R1-32 efficiently impairs the fusogenic conformational change of the WT S-trimer, it only partially inhibits this transition in the BA.4/5 and EG.5 S-trimers **(Fig. S4b, d, and f)**, likely due to the presence of L452_SARS2_ or F490_SARS2_ mutations in these variants. This reduced inhibition correlates with R1-32’s diminished neutralization activity against the corresponding variants.

Together with our cryo-EM data showing disassembled S-trimers, the results from ligand-induced conformational change assays confirm that the four identified R1-32-like antibodies retain the ability to inhibit ACE2-induced fusogenic conformational changes of S-trimers from variant viruses, likely by destabilizing or disassembling S-trimers, as previously proposed for R1-32 against the WT virus ^27^. The results are further consistent with neutralization assays demonstrating superior resilience of C092, C807, BD56-104, and BD56-597 against L452_SARS2_- and F490_SARS2_-bearing variants compared to R1-32.

### Broadly neutralizing public antibodies maintain binding to the mutated convergent epitope

The newly determined C092, C807, BD56-104, and BD56-597 complex structures were aligned to our previously reported R1-32 complex structure (PDB: 7YDI), using the RBD as the reference **(Fig. 2c)**. Based on this alignment, the root mean square deviations (RMSDs) of the Fab variable regions were calculated as follows: C092 (1.5 Å), C807 (1.8 Å), BD56-104 (1.7 Å), and BD56-597 (1.5 Å). The low RMSD values indicate that these antibodies adopt binding modes and recognize epitopes nearly identical to those of R1-32 **(Fig. 2c, left panel)**.

Antibody interface analysis reveals that all four R1-32-like antibodies engage the RBD through both their heavy and light chains, with a predominant contribution from the heavy chains: C092 binds an epitope area of 1,350 Å^2^ (HC: 903 Å^2^, LC: 447 Å^2^); C807 binds an epitope area of 1,193 Å^2^ (HC: 820 Å^2^, LC: 373 Å^2^); BD56-104 binds an epitope area of 1,148 Å^2^ (HC: 765 Å^2^, LC: 383 Å^2^); and finally BD56-597 binds an epitope area of 1,290 Å^2^ (HC: 937 Å^2^, LC: 353 Å^2^). For comparison, the germline-like R1-32 binds an epitope area of 1,214 Å^2^ (HC: 813 Å^2^, LC: 401 Å^2^).

All four R1-32-like antibodies utilize their characteristic IGHV1-69 HCDR2, featuring hydrophobic residues at positions 52_H_, 54_H_, and 55_H_ (subscripts denote heavy (H) or light (L) chain), to engage the hydrophobic RBD residues F490_SARS2_ and L492_SARS2_ (**Fig. 3a-d, left panels**). Notably, within the epitopes of R1-32-like antibodies in the BA.4/5 RBD, the epitope substitution L452R_SARS2_ is present, with the aliphatic portion of R452_SARS2_ sidechain contacted by the hydrophobic HCDR2 residue 55_H_. HCDR3 residues 102_H_, 103_H_, 104_H_, and 107_H_ form polar interactions with RBD residues E465_SARS2_, R466_SARS2_, and I468_SARS2_ (**Fig. 3a-d, right panels**). Additionally, HCDR1 (**Fig. 3a-d, left panels**) and LCDR1 (**Fig. 3a-d, middle panels**) contribute to RBD binding in all four antibodies. Notably, in C092, unlike the other three antibodies, both the light chain framework region 3 (LFR3) and LCDR3 participate in RBD binding (**Fig. 3a, middle panel**). Also notably, in BD56-104 and BD56-597, the heavy chain backbone region 3 (HFR3) participates in RBD binding (**Fig. 3c and d, left panels**).

**Fig. 3.**
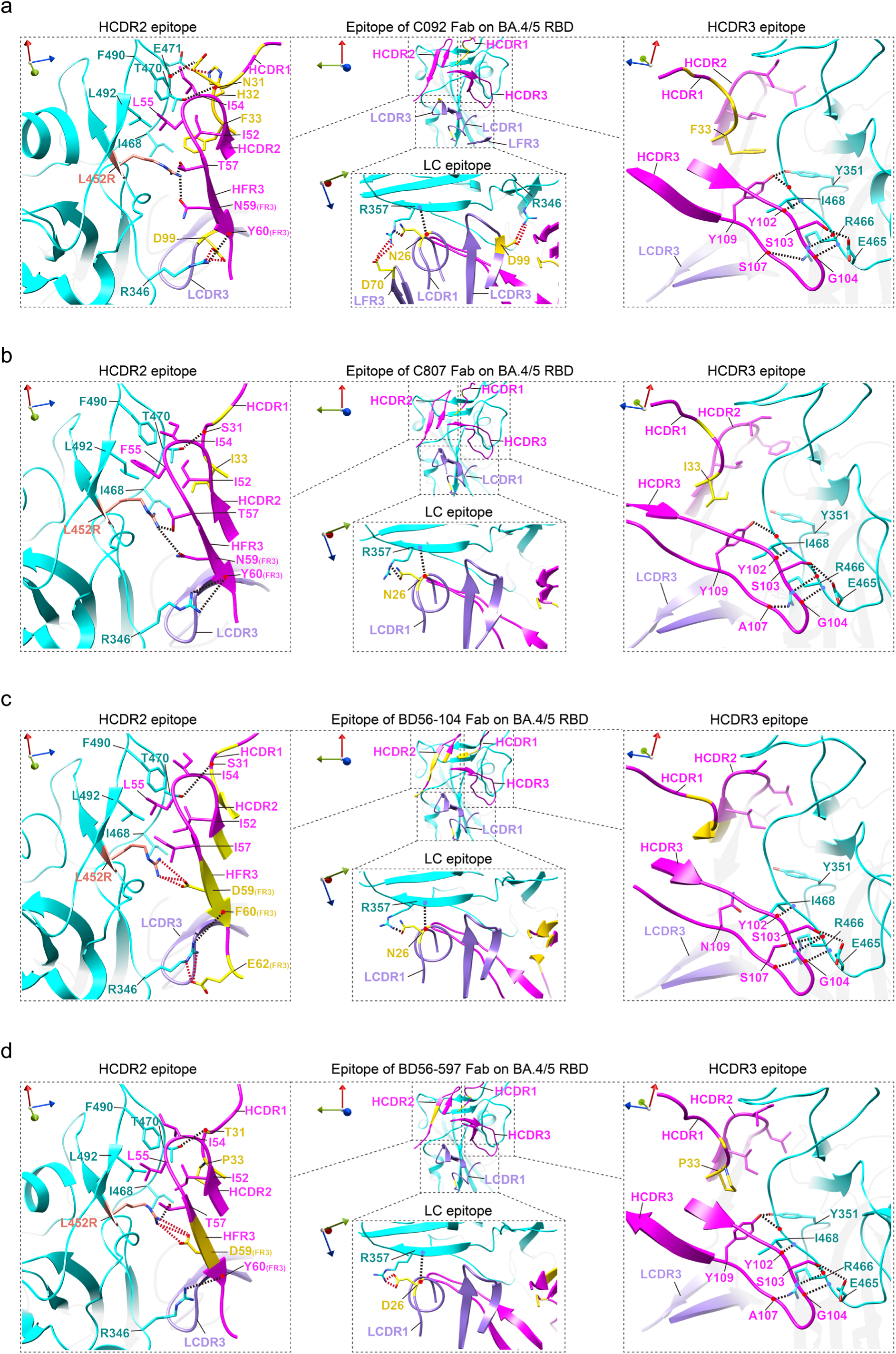
Epitopes of mutation-tolerant antibodies on BA.4/5 RBD. **a-d**, HCDR2, HCDR3, and LC epitopes of C092 (**a**), C807 (**b**), BD56-104 (**c**), BD56-597 (**d**) are shown from rotated views. RBD, Fab-HC, and Fab-LC residues are colored in cyan, magenta, and purple, respectively. Antibody residues introduced by somatic hypermutation are colored in yellow, while residue L452R mutated in BA.4/5 RBD is colored in salmon. The backbone carbonyl oxygens and amide nitrogens are indicated by red and blue dots, respectively. Hydrogen bonds and salt bridges are shown as black and red dash lines, respectively.

We previously found that somatic hypermutation (SHM)-introduced residues play a key role in enhancing antigen binding and conferring mutation tolerance ^22,37^. Compared with the germline antibody R1-32 (**Fig. S9e**), all four antibodies harbor SHM-introduced residues in HCDR1 and LCDR1 (**Fig. 3a-d**). In C092, SHM-introduced residues are also present in LFR3 and LCDR3 (**Fig. 3a**). In BD56-104 and BD56-597, SHM-introduced residues are also found in heavy chain framework region 3 (HFR3) (**Fig. 3c and d**). Many of these residues form additional contacts with the RBD, likely contributing to enhanced antigen binding (see below).

### Convergent somatic hypermutations confer enhanced binding and mutation tolerance

C092, C807, BD56-104, and BD56-597 likely resist epitope mutations by having undergone affinity maturation through somatic hypermutation. To investigate the affinity maturation pathway underlying their mutation tolerance, we curated ninety-eight R1-32-like antibodies from sequences reported by Cao *et al.*^28^ and analyzed the correlation between SHM patterns and neutralization activity (IC_50_ values). We examined whether specific mutations are enriched in the R1-32-like antibodies isolated from individuals with different exposure histories **(Fig. S10)**, as well as antibodies escaped by or tolerant to BQ.1.1 (L452R_SARS2_) and XBB (F490S_SARS2_) **(Fig. 4a)**. We found several SHMs are highly enriched in R1-32-like antibodies that tolerant to BQ.1.1 and XBB, such as A33P/S_H_, S35T_H_, and N59D_H_ on the heavy chain and S26N_L_, G52A/V_L_, and S54N/T_L_ on the light chain **(Fig. 4a)**. In addition, these SHMs were enriched in R1-32-like antibodies isolated from patients with breakthrough infections (**Fig. S10**). Several of these SHM-enriched sites are also directly involved in epitope binding in C092, C807, BD56-104, and BD56-597 (**Fig. 3**).

**Fig. 4.**
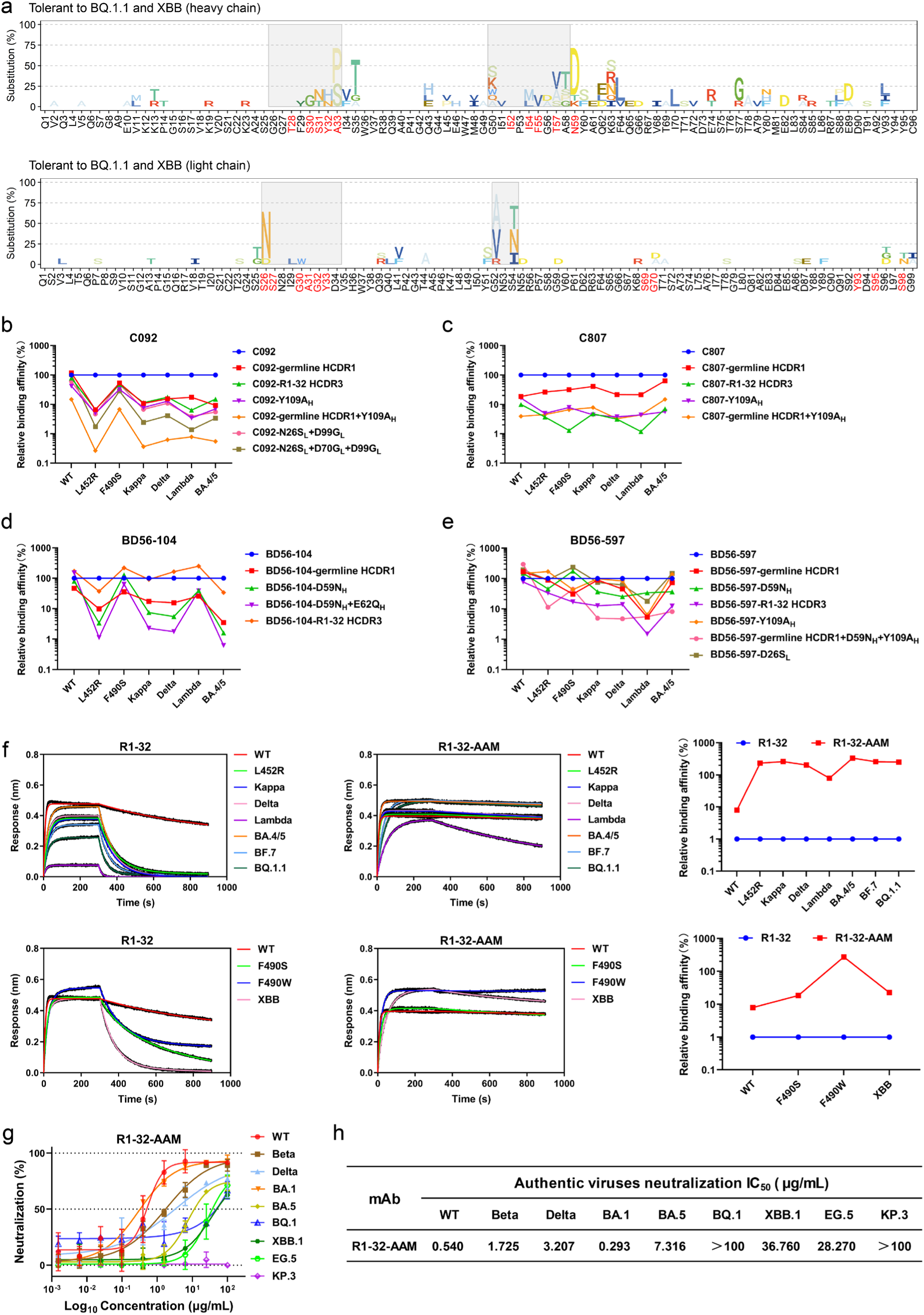
Affinity maturation confers mutation tolerance to R1-32-like antibodies. **a,** Frequency of somatic hypermutations (SHMs) in R1-32-like antibodies tolerant to BQ.1.1 and XBB. The residues involved in epitope binding are colored in red. **b-e**, Relative binding affinities of antibody mutants derived from C092 (**b**), C807 (**c**), BD56-104 (**d**), and BD56-597 (**e**) to the RBDs of SARS-CoV-2 wild-type, L452R, F490S, Kappa, Delta, Lambda, and BA.4/5. The constructed C092 mutants include C092-germline HCDR1 (HCDR1 reverted to germline sequence), C092-R1-32 HCDR3 (HCDR3 replaced with R1-32 HCDR3), C092-Y109A_H_, C092-germline HCDR1+Y109A_H_, C092-N26S_L_+D99G_L_, and C092-N26S_L_+D70G_L_+99G_L_. The C807 mutants include C807-germline HCDR1, C807-R1-32 HCDR3, C807-Y109A_H_, and C807-germline HCDR1+Y109A_H_. The BD56-104 mutants include BD56-104 germline HCDR1, BD56-104-D59N_H_, BD56-104+D59N_H_+E62Q_H_, and BD56-104-R1-32 HCDR3. The BD56-597 mutants include BD56-597-germline HCDR1, BD56-597-D59N_H_, BD56-597-R1-32 HCDR3, BD56-597-Y109A_H_, BD56-597-germline HCDR1-D59N_H_+Y109A_H_, BD56-597-D26S_L_. Affinity changes of mutants are normalized to *K*_D_s of the corresponding unmodified antibody (set as 100%). Binding curves are shown in **Fig. S11a-d,** and *K*_D_ values used for comparisons are shown in **Table S3**. **f**, Binding curves of R1-32 and R1-32-AAM to the RBDs of wild-type, L452R, Kappa, Delta, Lambda, BA.4/5, BF.7, BQ.1.1, F490S, F490W, and XBB. Affinity changes of R1-32-AAM are normalized to *K*_D_s of R1-32 (set as 1%). *K*_D_ values used for comparisons are shown in **Table S4**. **g**, Neutralization activities of R1-32-AAM towards SARS-CoV-2 wild-type, Beta, Delta, Omicron BA.1, BA.5, BQ.1, XBB.1, EG.5, and KP.3 authentic viruses. Data are presented as mean ± SD (n = 2). IC_50_ values are summarized in **h**.

In the C092:RBD complex, SHM introduced HCDR1 residues N31_H_ and H32_H_ form a hydrogen bond and a salt bridge with RBD residues T470_SARS2_ and E471_SARS2_, respectively (**Fig. 3a, left panel**). SHM introduced HCDR1 residue F33_H_ contacts RBD residues I468_SARS2_ and Y351_SARS2_, as well as HCDR2 residue I52_H_, thereby reinforcing the hydrophobic interactions involving I52_H_, I54_H_, and L55_H_ of HCDR2 and RBD residues F490_SARS2_ and L492_SARS2_ (**Fig. 3a, left and right panels**). Additionally, SHM-introduced residues in the light chain further enhance antigen binding: N26_L_ of LCDR1 and D70_L_ of LFR3 form hydrogen bonds and salt bridges with R357_SARS2_, while D99_L_ of LCDR3 forms salt bridges with R346_SARS2_ **(Fig. 3a, middle panel)**. To confirm the function of the SHM-introduced residues, we reverted them to their germline sequences and evaluated antigen binding by the revertants. Reverting C092 HCDR1 to germline reduced binding to L452R, Kappa (L452R_SARS2_), Delta (L452R_SARS2_), Lambda (L452Q_SARS2_+F490S_SARS2_), and BA.4/5 (L452R_SARS2_) RBDs by ∼ 10-fold (**Fig. 4b and Fig. S11a**). Likewise, reverting the light chain SHM-introduced residues (N26S_L_, D70G_L_, and D99G_L_) to germline reduced binding to the same set of RBDs by ∼10-100-fold.

In the C807:RBD complex, SHM-introduced HCDR1 residue I33_H_ contacts RBD residues I468_SARS2_ and Y351_SARS2_, as well as I52_H_ of HCDR2, via hydrophobic interactions **(Fig. 3b, left and right panels)**. Therefore, I33_H_ of C807, like F33_H_ in C092, enhances the hydrophobic interactions mediated by HCDR2. Reverting HCDR1 of C807 into the germline sequence reduced binding of RBDs with 452_SARS2_ and 490_SARS2_ mutations **(Fig. 4c and Fig. S11b)**.

In the BD56-104:RBD complex, SHM-introduced HFR3 residue D59_H_ forms salt bridges with the mutated RBD residue R452_SARS2_, while SHM-introduced HFR3 residue E62_H_ forms salt bridges with R346_SARS2_ **(Fig. 3c, left panel)**. Germline reversion of HFR3 residues (D59N_H_ and E62Q_H_) reduced binding to RBDs carrying 452_SARS2_ and 490_SARS2_ mutations. In particular, the combined BD56-104 HFR3 revertant (D59N_H_/E62Q_H_) showed an ∼100-fold reduction in binding to the BA.4/5 RBD harboring the L452R_SARS2_ mutation **(Fig. 4d and Fig. S11c)**.

In the BD56-597:RBD complex, similar to C092 and C807, SHM-introduced HCDR1 residue P33_H_ provides extra hydrophobic interactions to enhance antigen binding **(Fig. 3d, left and right panels)**. Further, similar to BD56-104, SHM-introduced HFR3 residue D59_H_ forms salt bridges with the mutated RBD residue R452_SARS2_ **(Fig. 3d, left panel)**. Notably, SHM-introduced LCDR1 residue D26_L_ interacts with R357_SARS2_ via salt bridges **(Fig. 3d, middle panel)**. Germline reversion of HCDR1, HFR3 (D59N_H_), and LCDR1 (D26S_L_) reduced binding to the RBDs carrying 452_SARS2_ and 490_SARS2_ mutations. Notably, towards the Lambda RBD, carrying both L452Q_SARS2_+F490S_SARS2_ mutations, a 10-fold reduction in binding was observed **(Fig. 4e and Fig. S11d)**.

### HCDR3 synergizes with somatic mutations to confer mutation tolerance

In addition to the extra interactions mediated by SHM-introduced residues, HCDR3 also contributes to enhanced antigen binding in C092, C807, and BD56-597. Antigen-binding assays showed that replacing the HCDR3s of C092, C807, or BD56-597 with that of R1-32 (97_H_-ARENGYSGYGAA**<u>A</u>**NFDL-113_H_, with A109_H_ underlined in bold) markedly reduced binding to the tested RBDs, with C807 being the most affected (**Fig. 4b-e and Fig. S11**). In contrast, HCDR3 substitution in BD56-104 had a comparatively minor impact on binding. Structural data revealed that the HCDR3s of C092, C807, and BD56-597 all contain a shared Y109_H_, which forms hydrogen bonds and hydrophobic interactions with RBD residues Y351_SARS2_ and I468_SARS2_ (**Fig. 3a, b, and d, right panels**). Substituting Y109_H_ with the alanine present in R1-32 also notably impaired antigen binding in these three antibodies. Moreover, combining HCDR1 germline reversion with the HCDR3 Y109A_H_ substitution in C092, C807, and BD56-597 produced an additive reduction in binding (**Fig. 4b-e and Fig. S11**), indicating that distinct antigen-contact regions act synergistically to achieve high-affinity binding.

### Artificial affinity maturation of R1-32 validates SHM-introduced residues in enhancing mutation tolerance

Our structural and antigen-binding data confirmed seven residues enriched with mutations and associated with enhanced antigen binding, A33F_H_, N59D_H_, Q62E_H_, and A109Y_H_ in the heavy chain, and S26D_L_, G70D_L_, and G99D_L_ in the light chain. We introduced these residues into R1-32 to generate an antibody called termed “R1-32-AAM” (R1-32-artificial-affinity-matured). Notably, these SHM-introduced residues not only mediate additional RBD interactions in C092, C807, BD56-104, and BD56-597, but most of them are also enriched in R1-32-like antibodies that retain tolerance to BQ.1.1 and XBB (**Fig. 4a**).

Binding assays showed that R1-32-AAM exhibits markedly enhanced antigen-binding activity compared with R1-32. Relative to R1-32, R1-32-AAM bound to RBDs bearing L452 mutation, including L452R_SARS2_, Kappa (L452R_SARS2_), Delta (L452R_SARS2_), Lambda (L452Q_SARS2_+F490S_SARS2_), BA.4/5 (L452R_SARS2_), BF.7 (L452R_SARS2_), and BQ.1.1 (L452R_SARS2_) with faster association, slower dissociation, and resulting in ∼100-fold enhanced affinity (**Fig. 4f**). It also displayed enhanced binding to RBDs carrying F490S_SARS2_ or F490W_SARS2_, and XBB variant (F490S_SARS2_), with ∼10-100-fold higher affinity than R1-32. Consistent with these binding data, R1-32-AAM neutralized authentic Delta (L452R_SARS2_), BA.5 (L452R_SARS2_), XBB.1 (F490S_SARS2_), and EG.5 (F490S_SARS2_) viruses (**Figs. 1f and g and 4g and h**), confirming that SHM-introduced residues enriched among R1-32-like antibodies confer tolerance to L452_SARS2_ and F490_SARS2_ substitutions. Together with structural differences to R1-32 (**Fig. S9e**), these findings indicate that the breadth of C092, C807, BD56-104, and BD56-597 is largely attributable to affinity-maturation-introduced residues, particularly within and around the CDRs, which mediate additional antigen interactions.

### Affinity maturation broadens the activity of R1-32-like antibodies toward SARSr-CoV RBDs

We further explored potential cross-reactivities of C092, C807, BD56-104, and BD56-597 toward SARSr-CoV RBDs using antigen-binding assays **(Fig. 5a and Fig. S12)**, testing representative type-1 to type-3 RBDs as previously defined by the extent of deletions within the receptor-binding motif (RBM) ^38^. All four antibodies bound the “type-1” RBDs of Pangolin-GD-2019, Bat-RaTG13, Pangolin-GX-2017, Laos-20-52, Laos-20-236, and Bat-RsSHC014 with affinities ranging from 0.016 to 48 nM. Notably, C092, C807, and BD56-597 cross-reacted to WIV1 RBD, whereas BD56-104 did not. Only C807 and BD56-597 bound the SARS-CoV-1 RBD, albeit with much compromised affinities (*K*_D_ < 341 nM) **(Fig. 5a)**. Remarkably, all four antibodies also bound the “type-2” BtKY72 RBD, despite its RBM sequence differing notably from the “type-1” SARS-CoV-2 RBD. Moreover, they retained binding to the “type-3” RBDs of GX2013, HeB2013, and RmYN02, which contain substantial loop deletions within the RBM region **(Fig. 5a)**. Among them, C092 and C807 exhibited tighter binding affinities than BD56-104 and BD56-597, demonstrating broader cross-reactivity towards SARSr-CoV RBDs. In contrast, R1-32 showed much weaker binding to the RBDs of Bat-RaTG13, Pangolin-GX-2017, and Bat-RsSHC014 (4.76-297 nM) than C092, C807, BD56-104, BD56-597, and failed to bind the RBDs of Bat-WIV1, SARS-CoV-1, BtKY72, GX2013, HeB2013, and RmYN02 **(Fig. 5a)**, indicating that affinity maturation confers broader SARSr-CoVs binding to R1-32-like antibodies.

**Fig. 5.**
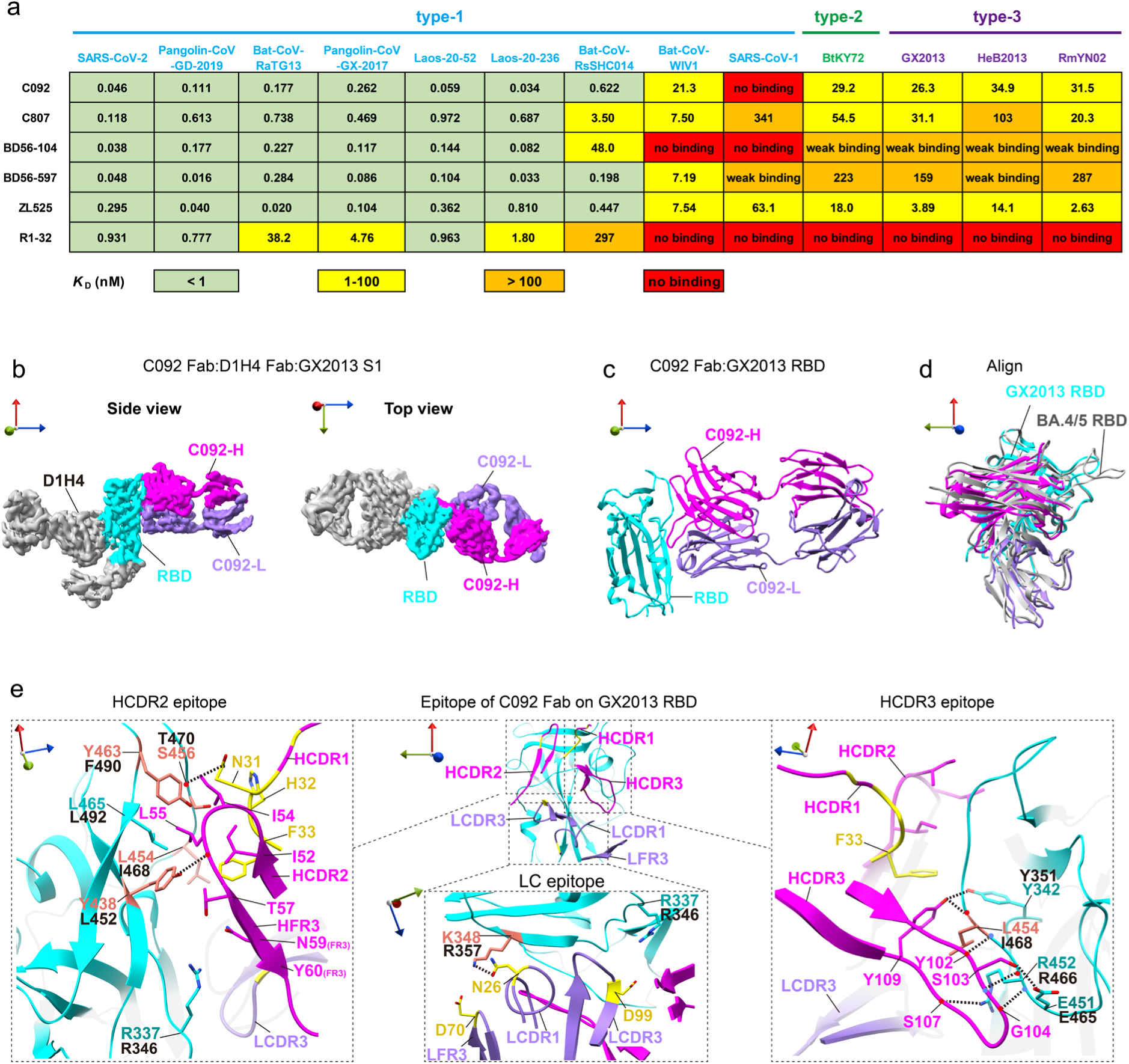
Affinity maturation confers R1-32-like antibodies with broader SARSr-CoVs binding. **a**, Binding affinities of R1-32-like antibodies to the RBDs of SARS-related coronaviruses (SARSr-CoVs). *K*_D_s (nM) are shown, with values <1, 1-100, >100 nM, and no binding colored in green, yellow, orange, and red, respectively. Binding curves and detailed kinetic parameters are provided in **Fig. S12 and Table S5**. **b**, Cryo-EM densities of the GX2013 S1:C092 Fab:D1H4 Fab complex. C092 Fab-HC, Fab-LC, and RBD are colored in magenta, purple, and cyan, respectively; the rest of the S1 and D1H4 Fab are colored grey. **c**, Structure of the GX2013 RBD:C092 Fab complex derived from the density in panel **b**. **d**, Comparison of C092 binding to GX2013 RBD and BA.4/5 RBD; the BA.4/5 RBD:C092 Fab complex is shown in grey. **e**, C092 HCDR2, HCDR3, and LC epitopes are shown from rotated views. Residues introduced by somatic hypermutation are colored in yellow, and epitope residues differing from SARS-CoV-2 wild-type RBD are colored in salmon. GX2013 RBD residues are labeled with the corresponding SARS-CoV-2 residues indicated in black. The backbone carbonyl oxygens and amide nitrogens are indicated by red and blue dots, respectively, and hydrogen bonds are shown as black dash lines.

We have established that C092, C807, BD56-104, and BD56-597 can well tolerate and maintain almost uncompromised neutralization towards variants carrying 1 mutation at RBD residues 452_SARS2_ or 346_SARS2,_ including Delta (L452R_SARS2_), BA.5 (L452R_SARS2_), and BA.2.75.2 (R346T_SARS2_). Neutralization was somewhat reduced towards variants with 2 mutations within the RBD epitope, including Lambda (L452Q_SARS2_+F490S_SARS2_), BQ.1.1 (R346T_SARS2_+L452R_SARS2_), XBB.1 (R346T_SARS2_+F490S_SARS2_), and EG.5 (R346T_SARS2_+F490S_SARS2_) **(Fig. 1c-g)**. We noted that all four R1-32-like antibodies exhibited markedly impaired binding to the SARS-CoV-1 RBD. Within the R1-32-like antibody epitope, SARS-CoV-1 and SARS-CoV-2 differ at L452_SARS2_, F490_SARS2_, R346_SARS2_, R357_SARS2_, and T470_SARS2_. To assess the mutation tolerance of the four R1-32-like antibodies (**Fig. S13)**, we introduced R346T_SARS2_, R357T_SARS2_, and T470N_SARS2_ mutations into the Lambda (L452Q_SARS2_+F490S_SARS2_) RBD, towards which C092, C807, BD56-104, and BD56-597 already exhibited 9.5-178-fold reduction in binding affinity (**Fig. 1c)**. Introduction each single mutation into the Lambda RBD further reduced binding affinity by 4.6-59-fold (**Fig. S13b)**. We previously found that antibody neutralization is more impacted by reduced antigen association than by accelerated antigen dissociation, as the latter can be compensated by avidity ^22^. Notably, these mutated RBDs impacted both antibody association and dissociation, with association rates (*k*_on_) reduced by 2.6-12-fold (**Fig. S13c)**, thereby 3 epitope mutations likely impact neutralization. For BD56-104, Lambda+R346T_SARS2_+T470N_SARS2_, completely abolished binding. For C092, C807, and BD56-597, we found that only when all three mutations (R346T_SARS2_, R357T_SARS2_, and T470N_SARS2_) were introduced simultaneously into the Lambda RBD did the antibodies completely lose binding **(Fig. S13b and c)**. These data suggest that C092, C807, BD56-104, and BD56-597 can still maintain antigen binding, withstanding 3-4 mutations in the epitope.

To further investigate cross-reactivity, the C092 Fab and D1H4 Fab ^22^ were incubated with the GX2013 S-trimer. Cryo-EM imaging of the sample revealed S-trimer disassembly. A 3.45 Å resolution structure of the C092 Fab:D1H4 Fab:GX2013 S1 complex was determined from disassembled S-trimer particles **(Fig. 5b and c and Fig. S14)**. The complex was aligned to the C092 Fab:BA.4/5 RBD complex using the RBDs as references to compare the difference of C092 binding to GX2013 and BA.4/5 RBDs. The result shows that C092 binds to the same epitope area on both the GX2013 and BA.4/5 RBDs **(Fig. 5d)**. C092 binds an epitope area of 1,003 Å^2^ on GX2013 RBD, with 675 Å^2^ buried by HCDRs, and 328 Å^2^ buried by LCDRs. This binding interface is smaller than the C092:BA.4/5 RBD interface (1,350 Å^2^), consistent with weaker binding of C092 to GX2013 RBD than BA.4/5.

In the C092:GX2013 RBD complex, the HCDR2 loop contains hydrophobic residues I52_H_, I54_H_, and L55_H_ and interacts with residues Y438_GX_ (L452_SARS2_), Y463_GX_ (F490_SARS2_), and L465_GX_ (L492_SARS2_) on GX2013 RBD via hydrophobic interaction. These hydrophobic interactions are further strengthened by the SHM-introduced HCDR1 residue F33_H_ via extra hydrophobic contacts **(Fig. 5e, left panels)**. The HCDR3 contacts GX2013 RBD residues - E451_GX_ (E465_SARS2_), R452_GX_ (R466_SARS2_), L454_GX_ (I468_SARS2_) and Y342_GX_ (Y351_SARS2_), with Y102_H_, S103_H_, G104_H_, S107_H_, and Y109_H_ via extensive hydrogen bonds **(Fig. 5e, right panel)**. Further, SHM-introduced HCDR1 N31_H_ and LCDR1 N26_L_ form hydrogen bonds with S456_GX_ (T470_SARS2_) and K348_GX_ (R357_SARS2_), respectively, to strengthen antigen binding. **(Fig. 5e, left and middle panel)**. Therefore, we conclude that affinity maturation is a key driver for broadening the reactivity of R1-32-like antibodies toward SARSr-CoV RBDs, with SHM-introduced residues mediating additional antibody-antigen interactions.

### AI-guided discovery of an ultrapotent broadly neutralizing antibody tolerant to glycan-mediated immune escape

Structure-guided artificial affinity maturation of R1-32 markedly enhanced its tolerance to viral escape mutations, whereas natural affinity maturation broadened the activity of this antibody class against sarbecoviruses with considerably diverged RBD epitopes, highlighting their potential for further expansion of breadth against emerging SARS-CoV-2 variants. Although C092, C807, BD56-104, and BD56-597 retained neutralizing activity against variants generally with 2 epitope mutations, including Lambda (L452Q_SARS2_+F490S_SARS2_), BQ.1.1 (R346T_SARS2_+L452R_SARS2_), XBB.1 (R346T_SARS2_+F490S_SARS2_), and EG.5 (R346T_SARS2_+F490S_SARS2_), the emerging KP.3 variant completely escaped their neutralization (**Fig. 1f and g**).

To discover antibodies with broader reactivity, we employed an AI-driven approach to discover R1-32-like antibodies with ultrapotent activities. We developed a deep learning model to predict antibody-antigen binding affinity, followed by subsequent stratified virtual screening to prioritize candidates (**Fig. 6a**). We pre-trained separate RoFormer-based ^39^ language models for antibody heavy and light chains and antigen, using a masked amino acid prediction task. Antigen representations were generated using the evolutionary-scale model ESM2 ^40^. Embeddings from both antibody and antigen models were integrated through a multilayer perceptron, followed by fine-tuning based on a consolidated antigen binding affinity dataset.

**Fig. 6.**
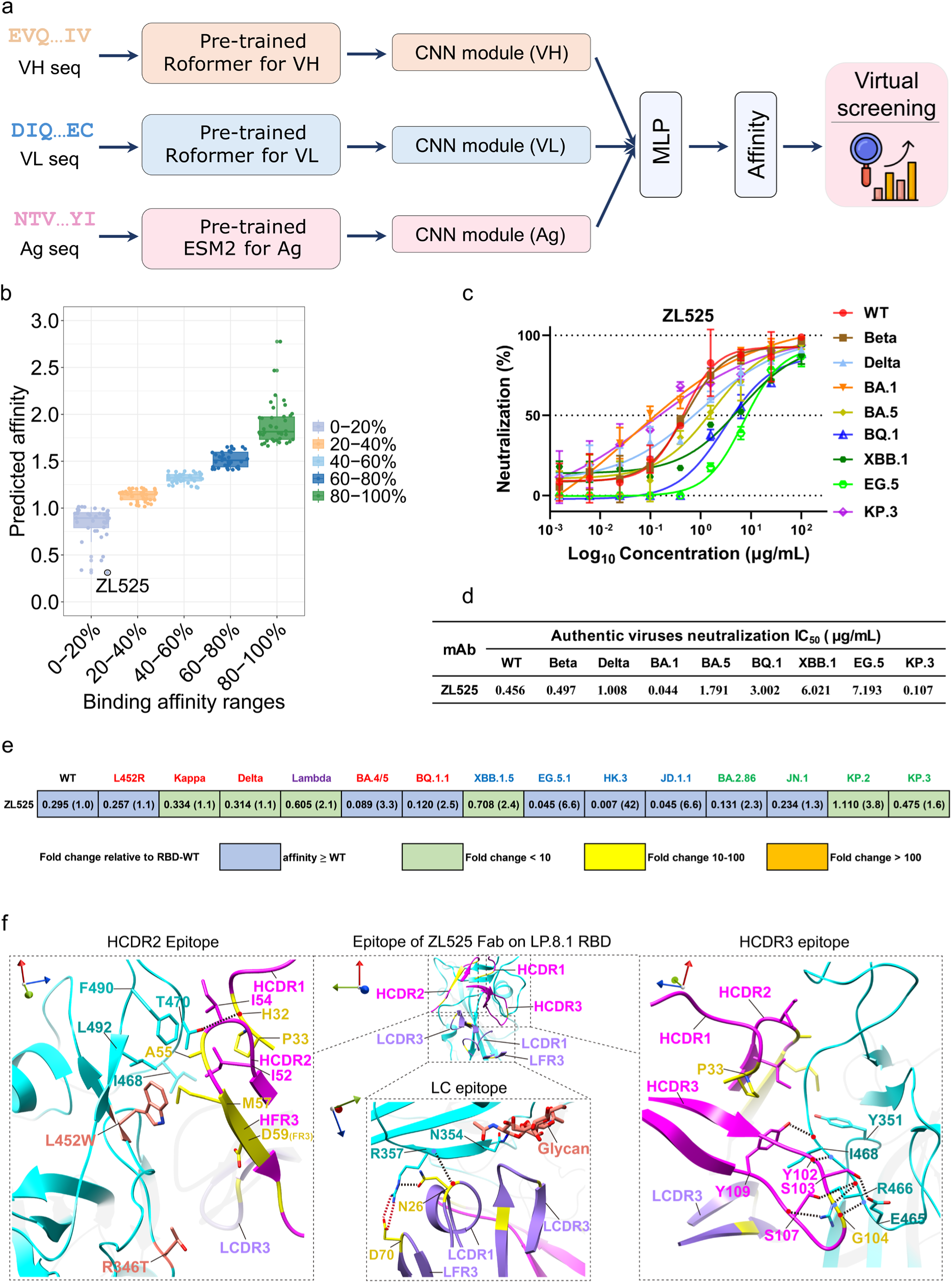
AI-guided discovery and structural characterization of the glycan-tolerant broadly neutralizing antibody ZL525. **a**, Workflow of the virtual screening pipeline based on pre-trained antibody language models. **b**, Distribution of predicted affinities of the 190 R1-32-like mAbs. **c**, Neutralization activities of the ZL525 towards SARS-CoV-2 wild-type, Beta, Delta, Omicron BA.1, BA.5, BQ.1, XBB.1, EG.5, and KP.3 authentic viruses. Data are shown as mean ± SD (n = 2). IC_50_ values are summarized in **d**. **e,** Binding affinities of ZL525 to RBDs carrying L452_SARS2_ or F490_SARS2_ substitutions and newly emerging variants. Dissociation constant *K*_D_s (nM) are shown, with fold changes relative to wild-type RBD indicated in parentheses. Binding affinities higher than the wild-type RBD are shown in blue, whereas reduced affinities (<10-fold, 10-100-fold, and >100-fold relative to wild-type) are shown in green, yellow, and orange, respectively. RBDs carrying L452, F490, or both substitutions are colored in red, blue, and purple, respectively. RBDs carrying L452 substitution together with N354 glycosylation are colored in green. Binding assays were performed by BLI, and detailed kinetic parameters are provided in **Table S8. f**, ZL525 HCDR2, HCDR3, and LC epitopes are shown from rotated views. Antibody residues introduced by somatic hypermutation are colored in yellow, and residues L452W_SARS2_ and R346T_SARS2_ mutated in the LP.8.1 RBD are colored in salmon. The backbone carbonyl oxygens and amide nitrogens are indicated by red and blue dots, respectively. Hydrogen bonds and salt bridges are shown as black and red dash lines, respectively.

The constructed affinity prediction model was used to screen 190 curated R1-32-like mAbs. A lead candidate, ZL525, which derived from an individual with sequential BA.5 and XBB breakthrough infections and exhibited higher predicted affinity than C092, C807, BD56-104, and BD56-597 (**Fig. 6b**). Authentic virus neutralization assays showed that ZL525 effectively neutralized all tested SARS-CoV-2 variants, including Beta, Delta, BA.1, BA.5, BQ.1, XBB.1, EG.5, and KP.3, with IC_50_ values ranging from 0.044 to 7.193 µg/mL **(Fig. 6c and d)**. Of note, ZL525 exhibited an excellent IC_50_ value of 0.107 µg/mL against KP.3, in contrast to C092, C807, BD56-104, and BD56-597, which completely lost neutralizing activity against the emerging KP.3 variant (IC_50_ > 100 µg/mL) **(Fig. 1f and g)**, underscoring ZL525 as an ultrapotent broadly neutralizing antibody. Consistent with the neutralization assays, ZL525 exhibited high affinities (below 0.71 nM) for the RBDs carrying L452_SARS2_, F490_SARS2_, or both mutations, including Delta, Kappa, Lambda, BA.4/5, BQ.1.1, XBB.1.5, and EG.5.1**(Fig. 6e and Fig. S16c)**. Furthermore, ZL525 maintained strong binding affinities (below 1.11 nM) for newly emergent variants, including HK.3, JD.1.1, BA.2.86, JN.1, KP.2, and KP.3, confirming its potential as a broadly therapeutic candidate. In addition, ZL525 demonstrated enhanced cross-reactivity towards sarbecoviruses, by comparison with C092, C807, BD56-104, and BD56-597, binding to all tested sarbecovirus RBDs, including SARS-CoV-1(**Fig. 5a**).

To understand ZL525 mutation tolerance, the LP.8.1 S-trimer was incubated with ZL525 Fab and ZL58 Fab (of a IGHV3-53 antibody). S-trimers in a 3-RBD “up” conformation with each RBD bound to a ZL525 Fab and a ZL58 Fab were imaged. By focused refinement, a 2.93Å ZL525 Fab: ZL58 Fab:LP.8.1 RBD complex structure was determined. **(Fig. 6f and Fig. S15)**. As expected, ZL525 recognizes the convergent epitope bound by other R1-32-like antibodies. Of note, its HCDR2 contains two characteristic SHM-introduced residues A55_H_ and M57_H_ to contact the RBD hydrophobic patch W452_SARS2_, F490_SARS2_, and L492 _SARS2_ on RBD **(Fig. 6f, left panel)**. Structural superposition of the BA.4/5 RBD:C092 and LP.8.1 RBD:ZL525 complexes reveals a conformational perturbation in the HCDR2 loop of ZL525. This conformational perturbation appears to allow HCDR2 to better contact the mutated L452W_SARS2_ residue **(Fig. S16a).** The above hydrophobic interactions are further enhanced by SHM-introduced HCDR1 residue P33_H_. Similar to C092, SHM-introduced residues LCDR1 residue N26_L_ and LFR3 residue D70_L_ form hydrogen bonds and salt bridges with R357, respectively, to further enhance interaction **(Fig. 6f, middle panel)**. Compared with C092, C807, BD56-104, and BD56-597, ZL525 exhibits a distinctive structural feature: SHM introduced residues within HCDR2 (A55_H_ and M57_H_) directly contact epitope. Although BD56-104 and BD56-597 contain SHM-introduced residues in HCDR2 (I51V_H_ and A58P_H_ in BD56-104, A58T_H_ in BD56-597), these residues are not involved in epitope contact with their side-chains orienting away from the epitope.

Notably, our structure reveals that the newly introduced N354_SARS2_ glycosylation ^41,42^ maps directly onto this public antibody epitope **(Fig. 6f, middle panel and Fig. S16b)**, suggesting that glycan shielding may obstruct antibody recognition. To test this, we engineered T356K_SARS2_ mutants of BA.2.86, JN.1, KP.2, and KP.3 RBDs to eliminate the N354_SARS2_ glycosylation. All four antibodies exhibited restored high-affinity binding to these glycan-revertants, with markedly improved association rates (*k*_on_) **(Fig. S16d)**, confirming that the N354_SARS2_ glycosylation contributes to their escape. In contrast, the AI-identified antibody ZL525 exhibits greater tolerance to this glycan-mediated shielding, likely due to structural adaptations introduced by somatic hypermutation. Collectively, these findings establish N354_SARS2_ glycan shielding as a key antigenic change driving immune evasion in emerging KP.3 lineage variants, consistent with previous reports^43^, and highlight ZL525 as a broadly neutralizing antibody capable of overcoming this glycan-mediated escape mechanism.

## Discussion

Through a series of structural and biochemical analyses, we now understand that affinity-matured R1-32-like public antibodies, including C092 and C807 ^27^ as well as BD56-104 and BD56-597 ^28^, retain binding to their convergent epitope, tolerating the L452_SARS2_ and F490_SARS2_ mutations through additional interactions mediated by SHM-introduced residues. We further find that affinity-matured C092, C807, BD56-104, and BD56-597 maintain neutralizing activity against viruses carrying at least two epitope mutations, exemplified by the Lambda variant (L452Q_SARS2_ + F490S_SARS2_). Together, these findings show that affinity maturation raises the genetic barrier to viral escape, such that the virus must acquire multiple coordinated epitope substitutions to overcome neutralization. As a result, affinity-matured antibodies may sustain protective immunity over longer periods, underscoring their essential role in maintaining durable immune protection.

In addition, C092, C807, BD56-104, and BD56-597 exhibit broad cross-reactivity with a diverse panel of SARS-related coronavirus (SARSr-CoV) RBDs, including Pangolin-GD-2019 ^44^, Bat-RaTG13, Pangolin-GX-2017 ^45^, Laos-20-52, Laos-20-236 ^46^, Bat-RsSHC014, Bat-WIV1^47^, SARS-CoV-1^48,49^, BtKY72, GX2013, HeB2013, and RmYN02 ^38^. This substantial expansion in breadth indicates that affinity maturation not only enhances tolerance to SARS-CoV-2 epitope mutations but also extends the reactivity of R1-32-like antibodies to antigenically diverse sarbecoviruses. The likely presence of such affinity-matured public antibodies further suggests that a degree of pre-existing immunity to diverse sarbecoviruses may now be present in the human population after the SARS-CoV-2 pandemic. The finding that affinity maturation broadens antibody reactivity toward heterologous sarbecoviruses, with substantially diverged epitopes, provides a conceptual framework for new strategies to design and discover broadly neutralizing antibodies that do not necessarily rely on targeting highly conserved epitopes.

Furthermore, structural characterization of affinity matured R1-32-like public antibodies revealed specific mutations relative to germline R1-32-like antibodies in and around the CDR loops, including A33F_H_, N59D_H_, Q62E_H_, A109Y_H_, S26D_L_, G70D_L_, and G99D_L_, that enhance antigen binding, increase resilience to epitope mutations, and expand antibody breadth. Notably, many of these sites (33_H_, 59_H_, 62_H_, and 26_L_) coincide with SHM-enriched sites in R1-32-like antibodies that tolerate BQ.1.1 (L452R_SARS2_) and XBB (F490S_SARS2_), suggesting convergent affinity-maturation patterns that promote increased resilience to antigenic drift. We confirmed this concept by introducing mutations at these seven sites into R1-32 to generate an artificially affinity-matured variant, R1-32-AAM, which exhibits improved tolerance to the L452_SARS2_ and F490_SARS2_ mutations. These findings demonstrate how deciphering in vivo maturation patterns can guide in vitro maturation strategies to enhance antibody potency for therapeutic applications, as previously envisioned ^50^.

R1-32-like public antibodies utilize their characteristic IGHV1-69 germline-encoded hydrophobic HCDR2 loops to bind a convergent epitope on the SARS-CoV-2 RBD, defined by a hydrophobic patch formed by residues L452_SARS2_, F490_SARS2_, and L492_SARS2_ ^16,27^. Likely owing to selection pressure from R1-32-like and other IGHV1-69 antibodies, residues L452_SARS2_ and F490_SARS2_ have become antigenic mutation hotspots ^16,51,52^, impairing antigen binding for most early-isolated R1-32-like antibodies that retained germline-like sequences and lacked extensive affinity maturation ^16,27^. The identification of R1-32-like antibodies that are tolerant to epitope mutations implies that members of this class likely remain active in the population and continue to exert immune pressure on this site. The emergence of the novel N354_SARS2_ glycosylation may reflect a viral adaptation to this pressure, highlighting viral evolution potentially driven by this class of public antibodies.

Since the emergence of the BA.2.86 variant (**Fig. S16b**), the N354_SARS2_ glycosylation mutation has become fixed in circulating strains ^41,43^. We have shown that the synergistic effect of sequence variation and glycan-mediated structural occlusion enables viral escape from previously identified mutation-tolerant R1-32-like antibodies (C092, C807, BD56-104, and BD56-597). To identify antibodies with even stronger mutation-tolerance, we applied AI-driven antibody language modelling and discovered an ultrapotent antibody, ZL525, which retains strong neutralizing activity against the KP.3 variant carrying the N354_SARS2_ glycosylation. Structural analysis indicates that SHM-introduced smaller or more flexible hydrophobic residues (L/F55A_H_ and I/T57M_H_) in HCDR2 likely confer enhanced conformational adaptability, allowing ZL525 to engage the altered hydrophobic surface formed by F490_SARS2_, L492_SARS2_, and more importantly the substituted L452W_SARS2_ (**Fig. 6f**), while counteracting the shielding effect of the newly introduced N354_SARS2_ glycan. Moreover, ZL525 extends cross-reactivity to the SARS-CoV-1 S-protein, which is substantially diverged from SARS-CoV-2. Together, these findings indicate that affinity maturation can further evolve R1-32-like antibodies to acquire breadth that not only counters viral escape but also enables cross-reactivity to more divergent sarbecoviruses.

The successful discovery of the ultrapotent broadly neutralizing antibody ZL525 through AI-driven language modelling highlights the transformative potential of machine learning in accelerating antibody discovery. By decoding the functional patterns of somatic hypermutations (SHMs) that confer broad neutralization, our study demonstrates how AI can systematically infer critical sequence-structure-function relationships that might otherwise require extensive experimental screening ^53^. This approach is particularly powerful for targeting rapidly evolving pathogens, like SARS-CoV-2, where conventional antibody discovery methods struggle to keep pace with viral escape variants ^54^. Recent advances in deep learning-based protein modelling, such as AlphaFold and antibody-specific language models, have begun to bridge the gap between sequence analysis and functional prediction ^54^. Future integration of structural constraints with generative AI could further enhance the precision of antibody optimization ^55^, paving the way for rapid discovery of antibody therapeutics to counter emerging viral threats. Together, these findings illustrate the dynamic interplay between population-level antibody responses and viral antigenic evolution, and highlight how integrating mechanistic immunology with AI-driven discovery can accelerate the development of antibodies resilient to rapidly evolving pathogens.

## Methods

### COVID-19 patient and vaccine recipient enrollment

Between December 2022 and December 2023, we enrolled a cohort of 64 individuals with RT-PCR-confirmed SARS-CoV-2 infection. All participants had received three doses of an inactivated COVID-19 vaccine (Sinopharm or CoronaVac) before experiencing sequential breakthrough infections with the BA.5 and XBB subvariants, which corresponded to the two major epidemic waves during this period. All participants provided written informed consent before enrollment. The study was conducted under the approval and supervision of the Ethics Committee of the First Affiliated Hospital of Guangzhou Medical University (Ethics IDs: ES-2020-65).

### Cells and viruses

Expi293F cells (Thermo Fisher Scientific, Cat# A14527) were cultured in FreeStyle^TM^293 Expression Medium (Thermo Fisher Scientific, Cat# A1435101) in a shaking incubator at 37 °C with 8% CO_2_. Human embryonic kidney (HEK) 293T (ATCC, Cat# CRL-3216), HEK293T-ACE2, and Vero E6 cells (ATCC, Cat# CRL-1586) were cultured in 10% FBS-supplemented Dulbecco’s Modified Eagle’s Medium (DMEM) at 37°C, 5% CO_2_. Expi293F cells were used for recombinant protein expression. HEK293T cells were used to generate VSV-based pseudoviruses, while HEK293T-ACE2 and Vero E6 cells were used for neutralization assays. VSV-based pseudotyped viruses were generated as previously described ^36^. Authentic SARS-CoV-2 viruses, including WT, Beta, Delta, and Omicron (BA.1, BA.4/5, BQ.1, XBB.1, EG.5, and KP.3), were isolated from clinical specimens of COVID-19 patients and stored at the Guangzhou Customs District Technology Center BSL-3 laboratory. The SARS-CoV-2 Delta strain was kindly provided by the Guangdong Provincial Center for Disease Control and Prevention, China. All experiments involving authentic SARS-CoV-2 were conducted under biosafety level 3 (BSL-3) conditions at the Guangzhou Customs District Technology Center BSL-3 Laboratory.

### Expression and purification of human monoclonal antibodies

The heavy chain (IgH) and light chain (IgL) genes of antibodies were amplified and cloned into the expression vector pCMV3 using ClonExpress Ultra One Step Cloning Kit (Vazyme, C115). When the density of Expi293F cells reached 2.5 × 10^6^ cells/mL, the paired IgH and IgL plasmids were transiently co-transfected into the cells at a ratio of 1:1 using transfection reagent Polyethylenimine (PEI). The transfected cells were transferred to a shaking incubator at 33 °C with 8% CO_2_. Five days post-transfection, the IgG was purified from the cell supernatant using protein A affinity chromatography. Firstly, the supernatant was harvested by centrifugation and incubated with Protein A Resin (Genscript, China) for 2 h to enable antibody binding. After the resin was washed with PBS (Gibco), pH 7.4, the IgG bound to the resin was eluted using 0.1 M citric acid, pH 3.0, and neutralized immediately with an equal volume of 1 M Tris-HCl, pH 8.0, to maintain the pH within the neutral range. Subsequently, the fractions containing IgG were concentrated and buffer-exchanged into PBS (pH 7.4) using a 50 kDa MWCO Amicon Ultra filtration device (Merck Millipore). Purified antibodies were aliquoted, flash-frozen, and stored at −80 °C until needed.

### Fab fragment production

To produce Fab gene fragment, the VH and the human IgG1 constant region CH1 were amplified from the IgH gene. The plasmids of paired Fab-IgH and IgL were transiently co-transfected into the Expi293F cells at a ratio of 1:1 using the transfection reagent Polyethylenimine (PEI) when the cell density reached 2.5 × 10^6^ cells/mL. Five days post-transfection at 33 °C, the culture supernatant was collected by centrifugation and incubated with LambdaFabSelect (Cytiva) resin for 2 h to enable Fab binding. After the resin was washed with PBS, the Fab bound to the resin was eluted with 0.1 M glycine (pH 2.0) into 1/2th volume 1 M Tris-HCl, pH 8.0, to maintain the pH within the neutral range. Subsequently, the target elutions were concentrated by a 30 kDa MWCO Amicon Ultra filtration device (Merck Millipore). Finally, the Fab was further purified and buffer-exchanged into PBS using a Superdex 200 increase 10/300 GL column (Cytiva). Purified Fab was aliquoted, flash frozen, and stored at −80 °C until needed.

### Expression and purification of SARS-CoV-2 S trimer and RBD

SARS-CoV-2 Omicron BA.4/5 S trimer ectodomain (14-1211) with HexaPro ^56^ mutations (F817P, A892P, A899P, A942P, K986P, and V987P) and deletion of amino acids “PRRA” (681-684) at the fusion site was constructed into the expression vector pCDNA3.1 with an N-terminal µ-phosphatase signal peptide and a C-terminal TEV-cleavage site, a T4 foldon trimerization motif, and a hexahistidine tag ^57,58^. Various SARS-CoV-2 variants S trimer ectodomain (S-R) with “PRRA” (681-684) deletion at the fusion site were also constructed using the same method. The coding sequence of SARS-CoV-2 RBD (residues 332-527) with an N-terminal µ-phosphatase signal peptide and a C-terminal hexahistidine tag was cloned into the expression vector pCDNA3.1 ^57^. All expression plasmids of S-trimer and RBD were transiently transfected into Expi293F cells using transfection reagent Polyethylenimine ^59^. Five days post-transfection at 33 °C, the cultural supernatant was collected and supplemented with 25 mM phosphate, pH 8.0, 300 mM NaCl, 5 mM imidazole, and 0.5 mM PMSF recirculated onto a HiTrap TALON crude column (Cytiva). Subsequently, the column was washed with buffer A (25 mM phosphate, pH 8.0, 300 mM NaCl, 5 mM imidazole), and target protein was eluted with a 100 mL linear gradient to 100% buffer B (25 mM phosphate, pH 8.0, 300 mM NaCl, 500 mM imidazole). Fractions containing the target protein were concentrated with an MWCO Amicon Ultra filtration device (Merck Millipore) and buffer-exchanged into PBS. Additionally, all RBDs were further purified and buffer-exchanged into PBS using a Superdex 75 increase 10/300 GL column (Cytiva) to remove RBD dimers. All SARS-CoV-2 spike and RBD variants were purified as described above. Purified S-trimers and RBDs were aliquoted, flash frozen, and stored at −80 °C until needed.

### Binding kinetics and affinity assessment of antibodies by BLI

Binding kinetics and affinities of mAbs against SARS-CoV-2 spikes or RBDs and SARSr-CoVs RBDs were assessed by Biolayer Interferometry (BLI) on an Octet RED96 instrument (FortéBio, USA). The assays were performed at 25 °C, and all proteins were diluted to PBST buffer (PBS with 1mg/mL BSA and 0.02% v/v Tween-20). Biosensors were pre-equilibrated in the PBST buffer for 10 min before the experiments. Initially, antibodies (11 µg/mL) were immobilized onto Protein A biosensors to a level of ∼1.5 nm. After a 60 s baseline step in PBST buffer, biosensors loaded with mAb were exposed (300 s) to the spikes (from 200 nM to 3.125 nM in 2-fold serial dilutions) or various RBDs (200 nM) to measure association. Subsequently, the biosensors were dipped into PBST (600 s) to measure dissociation of spikes or RBDs from the biosensor surface. A blank reference was set for each reaction. Data was reference-subtracted and analyzed using the FortéBio data analysis software HT v12.0.2.59 (FortéBio) by fitting to single or double phase association and dissociation kinetics to determine *k*_on_, *k*_off_ and *K*_D_. Raw data and fit data were plotted in GraphPad Prism 8.0.

### Antibody competition assay by BLI

The competitive assays between antibodies or between antibodies and ACE2 were measured by Biolayer Interferometry (BLI) using an Octet RED96 instrument (FortéBio, USA). All experimental steps were performed at 25 °C. All proteins were diluted to PBST buffer (PBS with 1mg/mL BSA and 0.02% v/v Tween-20), and anti-His biosensors were pre-equilibrated in PBST buffer for 10 min before being immobilized with purified SARS-CoV-2 RBD-His protein (30 μg/mL). Biosensors loaded with RBD were saturated with the first ligand (antibodies or ACE2 with a concentration of 200nM), and then binding of the second ligand (antibodies or ACE2 with a concentration of 200nM) or buffer as control was measured for 300 s. Data was analyzed using the FortéBio data analysis software HT v12.0.2.59 (FortéBio) and raw data was plotted in GraphPad Prism 8.0.

### Pseudovirus neutralization assay

The neutralization potency of antibodies against SARS-CoV-2 wild-type and multiple variants (including Delta, Lambda, and Omicron sublineages BA.1, BA.2, BA.4/5, BA.2.75.2, BQ.1.1, and EG.5) was evaluated using a pseudovirus neutralization test following established protocols ^36^. The antibody samples were diluted to an initial concentration of 10 μg/mL and subjected to eight gradient dilutions (3× serial dilutions), while the pseudovirus was diluted to 1.3×10^4^ TCID_50_/mL. After incubating the antibody samples and pseudovirus at 37 °C for 1 hour, HEK293T-ACE2 cells were added. After 24 hours of culture, luciferase detection reagent was added and placed in the multifunctional micropore detector (PE Ensight) to read the luminous value. Results were analyzed, and IC_50_ values were calculated by GraphPad Prism 8.0.

### Authentic virus neutralization assay

All experiments involving authentic SARS-CoV-2 virus were conducted under biosafety level 3 (BSL-3) containment at the Guangzhou Customs District Technology Center. Antibodies were initially diluted to 100 μg/mL and then subjected to 4-fold serial dilutions. Each diluted mAb sample was combined with an equal volume of virus suspension containing 200 focus-forming units (FFU) of specified SARS-CoV-2 strains (WT, Beta, Delta, and Omicron subvariants BA.1, BA.4/5, BQ.1, XBB.1, EG.5, and KP.3). Mixtures were added to Vero E6 cells in 96-well cell culture plates for 1 h at 37 °C. After removal of the mixtures, fresh overlay medium consisting of MEM supplemented with 1.2% carboxymethylcellulose was added to each well. Following 24 hours of incubation, the overlay was discarded, and the cells were fixed with 4% paraformaldehyde (Biosharp, China, Cat# BL539A) and permeabilized using 0.2% Triton X-100 (Sigma, USA, Cat# T8787). Cells were incubated with a human anti-SARS-CoV-2 nucleocapsid protein monoclonal antibody (obtained by laboratory screening) at 37 °C for 1 h. After three washes with 0.15% PBST, cells were incubated with an HRP-labeled goat anti-human secondary antibody (Jackson ImmunoResearch Laboratories, Cat# 609-035-213) at 37 °C for 1 h. Following additional washes, foci were developed using TrueBlue Peroxidase Substrate (KPL, Gaithersburg, MD, Cat# 50-78-02), and counted with an ELISPOT reader (Cellular Technology Ltd. Cleveland, OH). The foci reduction neutralization test titer (FRNT_50_) was calculated by the SpearmanKarber method.

### Cryo-EM sample preparation and data collection

The BA.4/5 S-R/6P S-trimer was mixed with the corresponding Fab and ACE2 at a 3(S-protomer):3(Fab):3(ACE2) molar ratio for 1 minute to form the C092, C807, BD56-104, and BD56-597 Fab complexes. Similarly, the GX2013 S-trimer was mixed with C092 and D1H4, and the LP.8.1 S-trimer was mixed with ZL525 and ZL58, each at a 3:3:3 molar ratio for 1 minute to assemble the C092 Fab:D1H4 Fab:GX2013 S-trimer and ZL525 Fab:ZL58 Fab:LP.8.1 S-R/6P complexes, respectively. All mixtures contained 0.1% octyl glucoside, except for the C092 Fab:D1H4 Fab:GX2013 complex, which contained 0.05% octyl glucoside. Samples were loaded onto freshly glow-discharged (15 mA, 30 s) holey carbon grids (Quantifoil, Cu R1.2/R1.3) at 4 °C and 100% humidity. The grids were blotted for 2.5 s with a force of 4 and samples were frozen immediately in liquid ethane using a Vitrobot (Thermo Fisher).

For the C092 Fab:ACE2:Omicron BA.4/5 S-R/6P and C092 Fab:D1H4 Fab:GX2013 S-trimer complexes, cryo-EM data were collected using a 200 keV Talos Arctica electron microscope (ThermoFisher Scientific) equipped with a K3 direct electron detector (Gatan). Movies were recorded using the SerialEM version 3.8.7 software at a nominal magnification of 45,000× with a calibrated pixel size of 0.88 Å and a defocus range from −0.8 to −2.5 μm. Samples were exposed for 1.83 s with a dose rate of 25.4 e^−^/pixel/s fractionated over 27 frames, giving a total electron dose of 60 e^−^/Å^2^.

To achieve higher resolution, a separately prepared C092 Fab:ACE2:BA.4/5 S-R/6P complex was additionally imaged in a Titan Krios electron microscope (Thermo Fisher Scientific) operating at 300 keV equipped with a SelectrisX energy filter (slit width 10 eV) and a Falcon 4 direct electron detector. The C807, BD56-104, and BD56-597 complexes, as well as the ZL525 Fab:ZL58 Fab:LP.8.1 S-R/6P complex, were also imaged in the Titan Krios electron microscope under the same setup. Movie stacks were automatically recorded using EPU with the electron event representation (EER) mode at a nominal magnification of ×165,000 with a calibrated pixel size of 0.73 Å and nominal defocus values ranging between −0.6 to −2.0 μm. Each stack was recorded and exposed at a dose rate of 6.53 e−/pixel/s for 4.08 s resulting a total dose of ∼ 50 e^−^/Å^2^. All movie stacks were imported into cryoSPARC live (v3.3.2/v4.2.0) ^60^ for pre-processing, which includes patched motion correction, contrast transfer function (CTF) estimation, and bad image rejection.

### Cryo-EM data processing

Data processing was carried out using cryoSPARC v3.3.2/v4.2.0. After removing bad images, particles were picked by blob-picking on images and extracted for 2D Classification. Well-defined 2D classes were selected as templates for template-picking.

For the C092 Fab:ACE2:BA.4/5 S-R/6P dataset collected at 200 keV **(Fig. S5)**, template-picked particles were extracted and subjected to two rounds of 2D classification to remove low-quality particles. Particles of either disassembled spike (S1) or Fab bound S-trimers were separated by 2D classification into two sub-datasets. For the S1 sub-dataset, Ab-initio Reconstruction was performed to generate initial 3D models, before the selected models were used as references in Heterogeneous Refinement. A well-defined structure was constructed by Non-uniform Refinement. To improve local resolution, a soft mask was applied to the ACE2:RBD:Fab interface for Local Refinement, yielding a final map at 3.84 Å resolution. For the S-trimer sub-dataset, three rounds of Ab-initial Reconstruction were conducted to remove bad particles. All resulting models from the final round were used as references in Heterogeneous Refinement. Particles generating highly similar structures were combined and subject to Non-uniform Refinement. This process yielded two distinct S-trimer reconstructions, featuring either 2 or 3 RBD “up” conformations, allowing simultaneous binding of either 2 or 3 Fab and ACE2 molecules.

To improve resolution, a separate C092 Fab:ACE2:BA.4/5 S-R/6P complex dataset was collected on a Titan Krios operating at 300 keV **(Fig. S6)**. Template-picked particles were similarly split into S1 and S-trimer subsets via 2D classification. For the S1 sub-dataset, Ab-initio Reconstruction and Heterogeneous Refinement were performed to generate 3D reconstructions. To improve reconstruction, Topaz Training and Picking was performed before picked particles were subjected to 2D Classification, Ab-initio Reconstruction, and Heterogeneous Refinement again. The final particles were subjected to Non-uniform Refinement, followed by Local Refinement with a mask focused on the ACE2:RBD:Fab interface regions to yield a final map at 2.58 Å resolution. For the S-trimer sub-dataset collected at 300 keV, A good 3D reconstruction was not obtained due to the absence of side-view projections in 2D classes, even after Topaz picking.

For the C807/BD56-104/BD56-597 Fab:ACE2:BA.4/5 S-R/6P datasets **(Figs. S6 and S7)**, Template-picked particles were similarly divided into two sub-datasets. The S1 sub-datasets were processed in a workflow identical to the C092 Fab:ACE2:BA.4/5 S-R/6P S1 sub-dataset collected at 300 keV. The S-trimer sub-datasets were processed using similar workflows, each yielding a structure showing a S-trimer simultaneously bound by three Fabs and three ACE2s. For the C807 Fab bound S-trimer sub-dataset, Topaz training was performed to further pick S-trimer particles. The Topaz picked particles were subjected to 2D Classification, Ab-initio Reconstruction, Heterogeneous Refinement, and Non-uniform Refinement to reconstruct a final map. For the BD56-104 Fab bound S-trimer sub-dataset, Topaz training and picking was performed after Heterogeneous Refinement. For the BD56-597 S-trimer sub-dataset, reconstruction was obtained without performing Topaz picking.

For the C092 Fab:D1H4 Fab:GX2013 dataset collected at 200 keV **(Fig. S14)**, Fab bound S1 particles from disassembled S-trimers were template-picked before subjecting to 2D Classification, Ab-initio Reconstruction, and Homogeneous Refinement. Topaz training and picking were performed to further pick particles. The final reconstruction was obtained using Topaz picked particles using Non-uniform Refinement, with a resolution of 3.45 Å.

For the ZL525 Fab:ZL58 Fab:LP.8.1 S-R/6P dataset **(Fig. S15)**, only Fab bound S-trimers were observed by auto-picking and subsequent 2D classification. S-trimer particles were subject to Ab-initio Reconstruction, Homogeneous Refinement, and Non-uniform Refinement, imposing C3 symmetry to generate a map with a resolution of 2.75 Å. To improve the resolution of the RBD:Fab interfaces, 3D classification with a focused mask (ZL525 Fab:ZL58 Fab:LP.8.1 RBD) was applied to remove low-quality particles, followed by Local Refinement. Ultimately, the focused ZL525 Fab:ZL58 Fab:LP.8.1 RBD map was refined to a resolution of 2.93 Å.

All resolutions were estimated at the 0.143 criterion from phase-randomization-corrected Fourier shell correlation (FSC) curves calculated between two independently refined half-maps, multiplied by soft-edged solvent masks, in either RELION v4.0 or cryoSPARC ^60^. Additional data processing details are summarized in **Figs. S5-8** and **Table S9**.

### Cryo-EM model building and structure refinement

A previously determined structure of R1-32 Fab:ACE2:SARS-CoV-2 RBD complex (PDB: 7YDI) was fitted into the Cryo-EM map in UCSF Chimera v1.15 ^61^ and used as the starting model for model building of the four complexes (C092/C807/BD56-104/BD56-597 Fab:ACE2:Omicron BA.4/5 RBD. Manual model building was carried out in Coot ^62^ in order to adjust Ramachandran, rotamers, bond geometry restraints, etc. Structure refinement was performed automatically using real-space refinement in PHENIX ^63^. For C092/C807/BD56-104/BD56-597 Fab:ACE2:Omicron BA.4/5 S-timer complexes, the structures (PDB:7YE5/7YEG) were used as initial model. For C092 Fab:D1H4 Fab:GX2013 RBD and ZL525 Fab:ZL58 Fab LP.8.1 RBD complexes, the structures (PDB:8ZY6 and 8WXL) were used as initial model respectively. Model validation statistics were summarized in **Table S9**.

### Ligand-induced conformational change assay

SARS-CoV-2 S-R spike at 1 mg/mL (7.09 µM) was incubated with ACE2-Fc or antibodies at a 1:1.1 molar ratio for 1 h at room temperature. The samples were subsequently treated with 50 µg/mL proteinase K for 30 min at 4 °C. Non-reducing SDS-PAGE loading buffer (5×) was added to each sample immediately before boiling at 98 °C for 5 min to stop the reaction. Samples were separated by SDS–PAGE and transferred onto a PVDF membrane. After blocking with 5% (w/v) skimmed milk in TBST buffer, the membrane was incubated with a rabbit anti-SARS-CoV-2 S2 polyclonal antibody (1:2,500 dilution, Sino Biological, 40590-T62) and subsequently with a goat anti-rabbit IgG conjugated to horseradish peroxidase (1:1,000 dilution, Beyotime, A0208). The S proteins on the membrane were detected by chemical luminescence using Pierce ECL western blotting substrate (Thermo Fisher, 32106). To further assess the effect of non-ACE2 competing antibodies on S-R spike conformation, the S-R spike protein was pre-incubated with antibodies (1 h), followed by incubation with ACE2-Fc (1 h). Samples were analyzed by western blotting as described above.

### Pre-trained antibody language models, ESM2 model, and virtual screening

Similar to our previously described Ab2binder model ^31^, we pre-trained antibody-specific language models on 1.4 billion antibody sequences sourced from the OAS database. After removing duplicate sequences and those containing non-canonical amino acids, the curated dataset consisted of 1.2 billion heavy chain and 210 million light chain sequences. Using the RoFormer ^39^ architecture, separate models were pre-trained for heavy and light chains. A Unique Amino Acid (UAA) tokenizer was adopted to represent each residue as a discrete token. Self-supervised learning was performed via a Masked Amino Acid (MAA) objective, in which randomly obscured residues were predicted from contextual cues, enabling the model to infer structural and functional properties of antibody sequences. To represent antigen inputs, we utilized the evolutionary-scale language model ESM2 (150M parameters), pre-trained on UniRef50 ^64^, to capture deep phylogenetic and structural features from protein sequences. Antigen embeddings were derived from ESM2 to encode functional constraints across diverse protein families. For antibody–antigen binding affinity prediction, features from both antibody chains, extracted using our pre-trained antibody models, were combined with ESM2-derived antigen embeddings. These representations were integrated via a multi-layer perceptron (MLP) to estimate binding affinities. The model was fine-tuned on a consolidated affinity dataset assembled from multiple public sources ^1,12,28,32,33,65–68^, enhancing its capacity to discern subtle interactions between antibodies and antigenic variants. Finally, we applied the optimized model to screen a library of 190 R1-32-like monoclonal antibodies in silico, of which 102 antibodies were derived from published data and 88 antibodies were isolated from convalescent individuals. Detailed information on isolating the latter set of antibodies will be described in a separate study ^69^. Candidates were ranked by predicted affinity, and top-scoring antibodies were prioritized for experimental validation.

## Data availability

Cryo-EM density maps for the structures of R1-32-like antibodies in complex with S-trimer or S1 fragment have been deposited in the Electron Microscopy Data Bank (EMDB) with accession codes EMD-66890, EMD-66891, EMD-66892, EMD-66893, EMD-66894, EMD-66895, EMD-66896, EMD-66897, EMD-66898, EMD-66899, EMD-66900, and EMD-66901. Related atomic models have been deposited in the Protein Data Bank (PDB) under accession codes 9XHY, 9XHZ, 9XI0, 9XI1, 9XI2, 9XI3, 9XI4, 9XI5, 9XI6, 9XI7, 9XI8, and 9XI9, respectively.

## Acknowledgements

We thank the staff of the GIBH-CAS Cryo-EM Facility for their help with cryo-EM sample preparation and data collection. We thank the staff of the Guangzhou Laboratory Cryo-EM Facility for their help with cryo-EM sample data collection. This work was supported by the National Key R&D Program of China (2021YFA1300903 to X.X.); the National Natural Science Foundation of China (82341085 and 32570199 to X.X., 82495200 and 82495203 to J.Z., 32361163669, 32170189, and 32241021 to J.H., 92469301 to L.C., 82201932 to Q.Y.); the Emergency Key Program of Guangzhou Laboratory (EKPG21-06 to X.X.); Major Project of Guangzhou National Laboratory (SRPG22-002 to X.X., EKPG21-30-2 to J.Z.); the Science and Technology Planning Project of Guangdong Province, China (2023B1212060050 and 2023B1212120009 to X.X.); the Basic Research Project of Guangzhou Institutes of Biomedicine and Health, Chinese Academy of Sciences (GIBHBRP24-02 to X.X.); the 111 Project (D18010 to J.Z.); the Natural Science Fund of Guangdong Province (2025A1515011245 to B.L.); the Young Doctoral Starting Sail Project of the Guangzhou Municipal Science and Technology Bureau (2024A04J4195 to B.L.). X.X. acknowledges Start-up grants from the Chinese Academy of Sciences.

## Contributions

Q.Y., C.N., X.H., and B.L. conceived the study under the direction of X.X. and J.Z.; Q.Y. performed the initial antibody analysis and screening from the published data; C.N., B.L., X.H. and P.H. expressed and purified antibodies; B.L., C.N., and J.W. purified spikes and RBDs for cryo-EM and other experiments using constructs and protocols developed by X.X.; C.N., B.L., and L.W. performed BLI assays; X.H., H.Z., and Y.Y. performed pseudovirus and authentic virus neutralization assays; C.N. performed western blots to assay spike structural change; C.N., B.L., X.G., H.Y., and B.Z. preformed cryo-EM sample preparation and collected cryo-EM data under the supervision of J.H. and X.X.; X.G., C.N., and Z.L. processed cryo-EM data under the supervision of J.H. and X.X.; X.X., C.N., and B.L. analysed cryo-EM structures with assistance from X.G. and Q.Y.; Y.S., Q.Y., Y.Y., and F.W. performed AI-based antibody discovery under the supervision of J.Y.; C.N., Q.Y., B.L., X.G. and X.H. prepared the figures under the supervision of X.X. and J.H.; X.X., C.N., and Q.Y. wrote the paper with input from all co-authors; G.H. and X.H. reviewed and edited the manuscript. X.X., Z.J., J.Y., J.H., L.C., and X.C. supervised the research.

## Reference

1 Cao, Y. et al. Imprinted SARS-CoV-2 humoral immunity induces convergent Omicron RBD evolution. Nature 614, 521–529, doi:10.1038/s41586-022-05644-7 (2023).

2 Murin, C. D., Wilson, I. A. & Ward, A. B. Antibody responses to viral infections: a structural perspective across three different enveloped viruses. Nat Microbiol 4, 734–747, doi:10.1038/s41564-019-0392-y (2019).

3 Cao, Y. et al. Potent Neutralizing Antibodies against SARS-CoV-2 Identified by High-Throughput Single-Cell Sequencing of Convalescent Patients’ B Cells. Cell 182, 73–84.e16, doi:10.1016/j.cell.2020.05.025 (2020).

4 Liu, Y. et al. The N501Y spike substitution enhances SARS-CoV-2 infection and transmission. Nature 602, 294–299, doi:10.1038/s41586-021-04245-0 (2022).

5 Mannar, D. et al. Structural analysis of receptor binding domain mutations in SARS-CoV-2 variants of concern that modulate ACE2 and antibody binding. Cell Rep 37, 110156, doi:10.1016/j.celrep.2021.110156 (2021).

6 Wang, R. et al. Analysis of SARS-CoV-2 variant mutations reveals neutralization escape mechanisms and the ability to use ACE2 receptors from additional species. Immunity 54, 1611–1621.e1615, doi:10.1016/j.immuni.2021.06.003 (2021).

7 Yuan, M. et al. Structural and functional ramifications of antigenicdrift in recent SARS-CoV-2 variants. science (2021).

8 Mannar, D. et al. SARS-CoV-2 Omicron variant: Antibody evasion and cryo-EM structure of spike protein-ACE2 complex. Science 375, 760–764, doi:10.1126/science.abn7760 (2022).

9 Barnes, C. O. et al. SARS-CoV-2 neutralizing antibody structures inform therapeutic strategies. Nature 588, 682–687, doi:10.1038/s41586-020-2852-1 (2020).

10 Sokal, A. et al. Maturation and persistence of the anti-SARS-CoV-2 memory B cell response. Cell 184, 1201–1213.e1214, doi:10.1016/j.cell.2021.01.050 (2021).

11 Yuan, M. et al. Structural basis of a shared antibody response to SARS-CoV-2. Science 369, 1119–1123, doi:10.1126/science.abd2321 (2020).

12 Muecksch, F. et al. Affinity maturation of SARS-CoV-2 neutralizing antibodies confers potency, breadth, and resilience to viral escape mutations. Immunity 54, 1853–1868 e1857, doi:10.1016/j.immuni.2021.07.008 (2021).

13 Chen, E. C. et al. Convergent antibody responses to the SARS-CoV-2 spike protein in convalescent and vaccinated individuals. Cell Rep 36, 109604, doi:10.1016/j.celrep.2021.109604 (2021).

14 Yan, Q. et al. Antibodies utilizing VL6-57 light chains target a convergent cryptic epitope on SARS-CoV-2 spike protein and potentially drive the genesis of Omicron variants. Nat Commun 15, 7585, doi:10.1038/s41467-024-51770-3 (2024).

15 Zhang, Y. et al. Analysis of B Cell Receptor Repertoires Reveals Key Signatures of the Systemic B Cell Response after SARS-CoV-2 Infection. J Virol 96, e0160021, doi:10.1128/jvi.01600-21 (2022).

16 Yan, Q. et al. Shared IGHV1-69-encoded neutralizing antibodies contribute to the emergence of L452R substitution in SARS-CoV-2 variants. Emerg Microbes Infect 11, 2749–2761, doi:10.1080/22221751.2022.2140611 (2022).

17 Patel, A. et al. Molecular basis of SARS-CoV-2 Omicron variant evasion from shared neutralizing antibody response. Structure 31, 801–811.e805, doi:10.1016/j.str.2023.04.010 (2023).

18 Kaku, C. I. et al. Evolution of antibody immunity following Omicron BA.1 breakthrough infection. Nat Commun 14, 2751, doi:10.1038/s41467-023-38345-4 (2023).

19 Yan, Q. et al. Germline IGHV3-53-encoded RBD-targeting neutralizing antibodies are commonly present in the antibody repertoires of COVID-19 patients. Emerg Microbes Infect 10, 1097–1111, doi:10.1080/22221751.2021.1925594 (2021).

20 Zhang, Q. et al. Potent and protective IGHV3-53/3-66 public antibodies and their shared escape mutant on the spike of SARS-CoV-2. Nat Commun 12, 4210, doi:10.1038/s41467-021-24514-w (2021).

21 Rapp, M. et al. Modular basis for potent SARS-CoV-2 neutralization by a prevalent VH1-2-derived antibody class. Cell Rep 35, 108950, doi:10.1016/j.celrep.2021.108950 (2021).

22 Liu, B. et al. An unconventional VH1-2 antibody tolerates escape mutations and shows an antigenic hotspot on SARS-CoV-2 spike. Cell Rep 43, 114265, doi:10.1016/j.celrep.2024.114265 (2024).

23 Dong, J. et al. Genetic and structural basis for SARS-CoV-2 variant neutralization by a two-antibody cocktail. Nat Microbiol 6, 1233–1244, doi:10.1038/s41564-021-00972-2 (2021).

24 Yuan, M. et al. Molecular analysis of a public cross-neutralizing antibody response to SARS-CoV-2. Cell Rep 41, 111650, doi:10.1016/j.celrep.2022.111650 (2022).

25 Wang, Y. et al. A large-scale systematic survey reveals recurring molecular features of public antibody responses to SARS-CoV-2. Immunity 55, 1105–1117.e1104, doi:10.1016/j.immuni.2022.03.019 (2022).

26 Westendorf, K. et al. LY-CoV1404 (bebtelovimab) potently neutralizes SARS-CoV-2 variants. Cell Rep 39, 110812, doi:10.1016/j.celrep.2022.110812 (2022).

27 He, P. et al. SARS-CoV-2 Delta and Omicron variants evade population antibody response by mutations in a single spike epitope. Nat Microbiol 7, 1635–1649, doi:10.1038/s41564-022-01235-4 (2022).

28 Cao, Y. et al. BA.2.12.1, BA.4 and BA.5 escape antibodies elicited by Omicron infection. Nature 608, 593–602, doi:10.1038/s41586-022-04980-y (2022).

29 Wang, Q. et al. Alarming antibody evasion properties of rising SARS-CoV-2 BQ and XBB subvariants. Cell 186, 279–286.e278, doi:10.1016/j.cell.2022.12.018 (2023).

30 Tegally, H. et al. Emergence of SARS-CoV-2 Omicron lineages BA.4 and BA.5 in South Africa. Nat Med 28, 1785–1790, doi:10.1038/s41591-022-01911-2 (2022).

31 He, H. et al. De novo generation of SARS-CoV-2 antibody CDRH3 with a pre-trained generative large language model. Nat Commun 15, 6867, doi:10.1038/s41467-024-50903-y (2024).

32 Gaebler, C. et al. Evolution of antibody immunity to SARS-CoV-2. Nature 591, 639–644, doi:10.1038/s41586-021-03207-w (2021).

33 Wang, Z. et al. Naturally enhanced neutralizing breadth against SARS-CoV-2 one year after infection. Nature 595, 426–431, doi:10.1038/s41586-021-03696-9 (2021).

34 Wang, Z. et al. mRNA vaccine-elicited antibodies to SARS-CoV-2 and circulating variants. Nature 592, 616–622, doi:10.1038/s41586-021-03324-6 (2021).

35 Zhang, L. et al. A proof of concept for neutralizing antibody-guided vaccine design against SARS-CoV-2. Natl Sci Rev 8, nwab053, doi:10.1093/nsr/nwab053 (2021).

36 Nie, J. et al. Quantification of SARS-CoV-2 neutralizing antibody by a pseudotyped virus-based assay. Nat Protoc 15, 3699–3715, doi:10.1038/s41596-020-0394-5 (2020).

37 Yu, H. et al. Somatically hypermutated antibodies isolated from SARS-CoV-2 Delta infected patients cross-neutralize heterologous variants. Nat Commun 14, 1058, doi:10.1038/s41467-023-36761-0 (2023).

38 Wang, J. et al. SARS-related coronavirus S-protein structures reveal synergistic RBM interactions underpinning high-affinity human ACE2 binding. Sci Adv 11, eadr8772, doi:10.1126/sciadv.adr8772 (2025).

39 Su, J. et al. RoFormer: Enhanced transformer with Rotary Position Embedding. Neurocomputing 568, 127063, 10.1016/j.neucom.2023.127063 (2024).

40 Lin, Z. et al. Evolutionary-scale prediction of atomic-level protein structure with a language model. Science 379, 1123–1130, doi:10.1126/science.ade2574 (2023).

41 Li, L. et al. Spike structures, receptor binding, and immune escape of recently circulating SARS-CoV-2 Omicron BA.2.86, JN.1, EG.5, EG.5.1, and HV.1 sub-variants. Structure 32, 1055–1067.e1056, doi:10.1016/j.str.2024.06.012 (2024).

42 Yang, H. et al. Structural basis for the evolution and antibody evasion of SARS-CoV-2 BA.2.86 and JN.1 subvariants. Nat Commun 15, 7715, doi:10.1038/s41467-024-51973-8 (2024).

43 Liu, P. et al. Spike N354 glycosylation augments SARS-CoV-2 fitness for human adaptation through structural plasticity. Natl Sci Rev 11, nwae206, doi:10.1093/nsr/nwae206 (2024).

44 Wrobel, A. G. et al. Structure and binding properties of Pangolin-CoV spike glycoprotein inform the evolution of SARS-CoV-2. Nat Commun 12, 837, doi:10.1038/s41467-021-21006-9 (2021).

45 Zhang, S. et al. Bat and pangolin coronavirus spike glycoprotein structures provide insights into SARS-CoV-2 evolution. Nat Commun 12, 1607, doi:10.1038/s41467-021-21767-3 (2021).

46 Ou, X. et al. Host susceptibility and structural and immunological insight of S proteins of two SARS-CoV-2 closely related bat coronaviruses. Cell Discov 9, 78, doi:10.1038/s41421-023-00581-9 (2023).

47 Qiao, S. & Wang, X. Structural determinants of spike infectivity in bat SARS-like coronaviruses RsSHC014 and WIV1. J Virol 98, e0034224, doi:10.1128/jvi.00342-24 (2024).

48 Zhang, X. et al. Disulfide stabilization reveals conserved dynamic features between SARS-CoV-1 and SARS-CoV-2 spikes. Life Sci Alliance 6, doi:10.26508/lsa.202201796 (2023).

49 Yuan, Y. et al. Cryo-EM structures of MERS-CoV and SARS-CoV spike glycoproteins reveal the dynamic receptor binding domains. Nat Commun 8, 15092, doi:10.1038/ncomms15092 (2017).

50 Vajda, S., Porter, K. A. & Kozakov, D. Progress toward improved understanding of antibody maturation. Curr Opin Struct Biol 67, 226–231, doi:10.1016/j.sbi.2020.11.008 (2021).

51 Pushparaj, P. et al. Immunoglobulin germline gene polymorphisms influence the function of SARS-CoV-2 neutralizing antibodies. Immunity, doi:10.1016/j.immuni.2022.12.005 (2022).

52 Starr, T. N., Greaney, A. J., Dingens, A. S. & Bloom, J. D. Complete map of SARS-CoV-2 RBD mutations that escape the monoclonal antibody LY-CoV555 and its cocktail with LY-CoV016. Cell Rep Med 2, 100255, doi:10.1016/j.xcrm.2021.100255 (2021).

53 Shan, S. et al. Deep learning guided optimization of human antibody against SARS-CoV-2 variants with broad neutralization. Proc Natl Acad Sci U S A 119, e2122954119, doi:10.1073/pnas.2122954119 (2022).

54 Desautels, T. A. et al. Computationally restoring the potency of a clinical antibody against Omicron. Nature 629, 878–885, doi:10.1038/s41586-024-07385-1 (2024).

55 Shanker, V. R., Bruun, T. U. J., Hie, B. L. & Kim, P. S. Unsupervised evolution of protein and antibody complexes with a structure-informed language model. Science 385, 46–53, doi:10.1126/science.adk8946 (2024).

56 Hsieh, C. L. et al. Structure-based design of prefusion-stabilized SARS-CoV-2 spikes. Science 369, 1501–1505, doi:10.1126/science.abd0826 (2020).

57 Xiong, X. et al. A thermostable, closed SARS-CoV-2 spike protein trimer. Nat Struct Mol Biol 27, 934–941, doi:10.1038/s41594-020-0478-5 (2020).

58 Qu, K. et al. Engineered disulfide reveals structural dynamics of locked SARS-CoV-2 spike. PLoS Pathog 18, e1010583, doi:10.1371/journal.ppat.1010583 (2022).

59 Yuan, H. et al. Structures and receptor binding activities of merbecovirus spike proteins reveal key signatures for human DPP4 adaptation. Sci Adv 11, eadv7296, doi:10.1126/sciadv.adv7296 (2025).

60 Kucukelbir, A., Sigworth, F. J. & Tagare, H. D. Quantifying the local resolution of cryo-EM density maps. Nat Methods 11, 63–65, doi:10.1038/nmeth.2727 (2014).

61 Pettersen, E. F. et al. UCSF Chimera--a visualization system for exploratory research and analysis. J Comput Chem 25, 1605–1612, doi:10.1002/jcc.20084 (2004).

62 Emsley, P., Lohkamp, B., Scott, W. G. & Cowtan, K. Features and development of Coot. Acta Crystallogr D Biol Crystallogr 66, 486–501, doi:10.1107/S0907444910007493 (2010).

63 Afonine, P. V. et al. Real-space refinement in PHENIX for cryo-EM and crystallography. Acta Crystallogr D Struct Biol 74, 531–544, doi:10.1107/s2059798318006551 (2018).

64 Suzek, B. E., Wang, Y., Huang, H., McGarvey, P. B. & Wu, C. H. UniRef clusters: a comprehensive and scalable alternative for improving sequence similarity searches. Bioinformatics 31, 926–932, doi:10.1093/bioinformatics/btu739 (2015).

65 Cao, Y. et al. Omicron escapes the majority of existing SARS-CoV-2 neutralizing antibodies. Nature 602, 657–663, doi:10.1038/s41586-021-04385-3 (2022).

66 Wang, Z. et al. Memory B cell responses to Omicron subvariants after SARS-CoV-2 mRNA breakthrough infection in humans. J Exp Med 219, doi:10.1084/jem.20221006 (2022).

67 Witte, L. et al. Epistasis lowers the genetic barrier to SARS-CoV-2 neutralizing antibody escape. Nat Commun 14, 302, doi:10.1038/s41467-023-35927-0 (2023).

68 Kaku, C. I. et al. Recall of preexisting cross-reactive B cell memory after Omicron BA.1 breakthrough infection. Sci Immunol 7, eabq3511, doi:10.1126/sciimmunol.abq3511 (2022).

69 Yan, Q. et al. AI recognizes convergent somatic hypermutation signatures to allow the discovery of variant-resilient broadly neutralizing antibodies. bioRxiv, 2025.2012.2010.693372, doi:10.64898/2025.12.10.693372 (2025).

