## Supplemental Figures 1-16 and Supplemental tables 1-9 for "Structural determinants of IGHV1-69 public antibodies conferring resilience to SARS-CoV-2 antigenic escape"

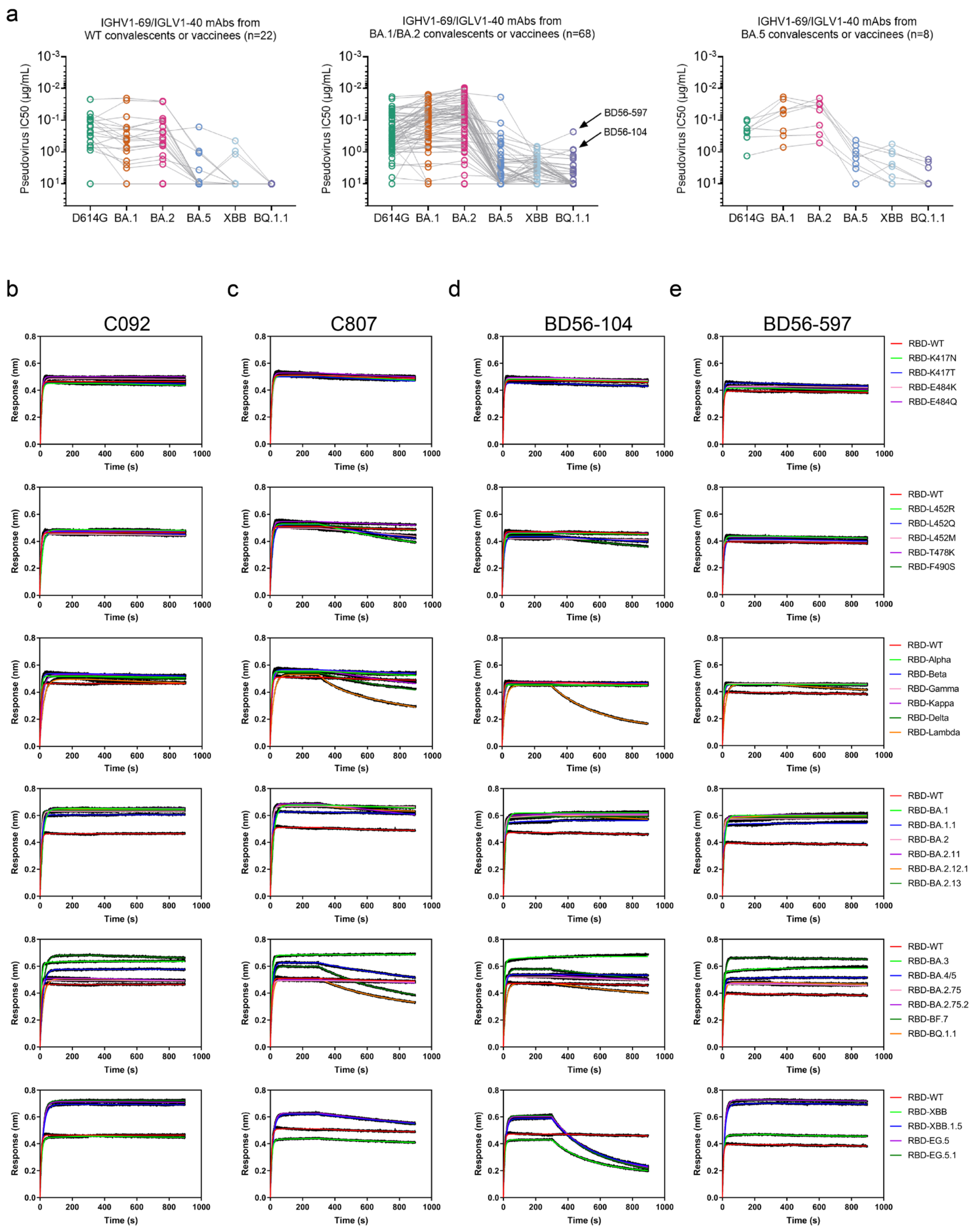


**Fig. S1. Binding of mutation-tolerant antibodies to the RBDs of SARS-CoV-2 variants.** **a**, Identification of antibodies BD56-104 and BD56-597 that may tolerate the $L452_{SARS2}$ mutation from the reported IGHV1-69/IGLV1-40 encoded antibody database. **b-e**, Binding curves of C092 (**b**), C807 (**c**), BD56-104 (**d**), and BD56-597 (**e**) to a panel of SARS-CoV-2 RBDs as analysed by BLI. Antibodies were immobilized onto Protein A biosensors and submerged into RBD solutions at a concentration of 200 nM. Detailed binding kinetic parameters are summarized in **Table S1**. The $K_D$ values and normalized $K_D$ fold changes are shown in **Fig. 1c**.

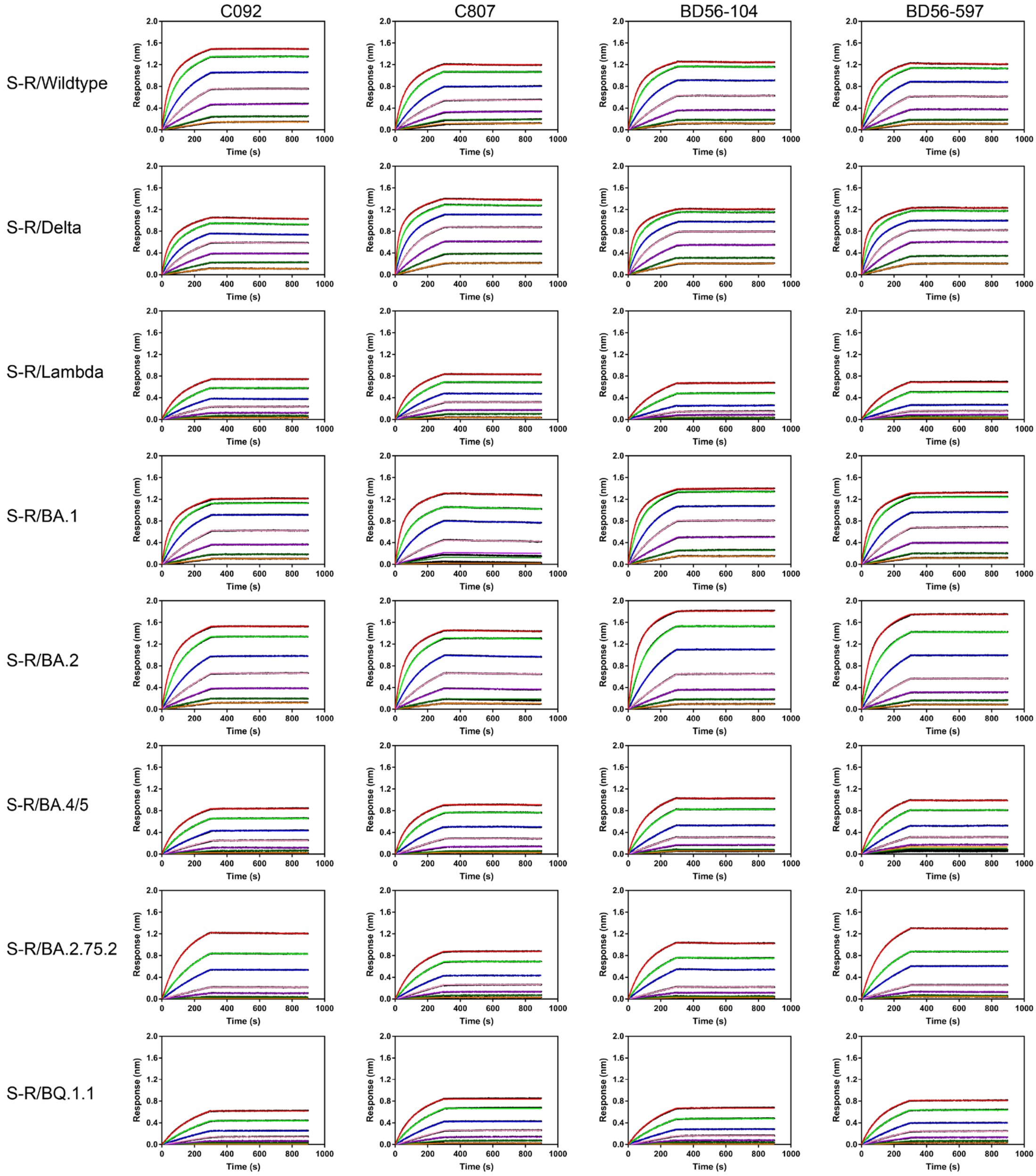


**Fig. S2. Binding of mutation-tolerant antibodies to the S-trimers of SARS-CoV-2 variants.**

**a**, Binding curves of C092, C807, BD56-104, and BD56-597 to SARS-CoV-2 wild-type, Delta, Lambda, Omicron BA.1, BA.2, BA.4/5, BA.2.75.2, and BQ.1.1 spikes are shown. Binding assays were performed using BLI, with antibodies immobilized onto Protein A sensors and surmeged into a dilution series of spike proteins (200 to 3.125 nM). Binding kinetics parameters are summarized in **Table S2**.

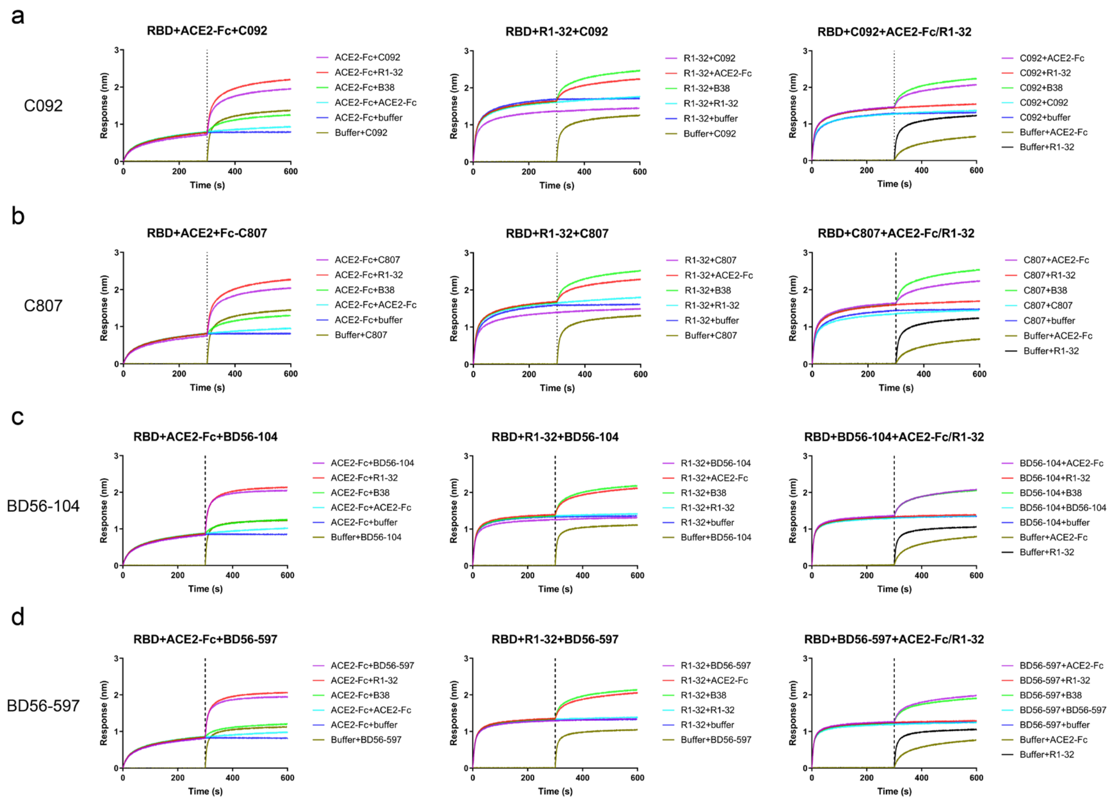


**Fig. S3. Competition assays between R1-32-like antibodies and ACE2-Fc or R1-32.** **a-d**, Competition binding to SARS-CoV-2 RBD between C092 (**a**), C807 (**b**), BD56-104 (**c**) and BD56-597 (**d**) and ACE2-Fc or R1-32 was assessed by BLI. SARS-CoV-2 RBD was immobilized onto HIS1K biosensors, and immobilized RBD was saturated with ACE2-Fc (left panels) or R1-32 (middle panels) before incubation with C092, C807, BD56-104, or BD56-597 (purple line). Binding of the immobilized RBD to only C092, C807, BD56-104, or BD56-597 (olive line) was assayed as controls. Conversely, the biosensors immobilized with the RBD were saturated with C092, C807, BD56-104, or BD56-597 (right panels) before incubation with ACE2-Fc (purple line) or R1-32 (red line). Binding of the immobilized RBD to only ACE2-Fc (olive line) or R1-32 (black line) was assayed as controls.

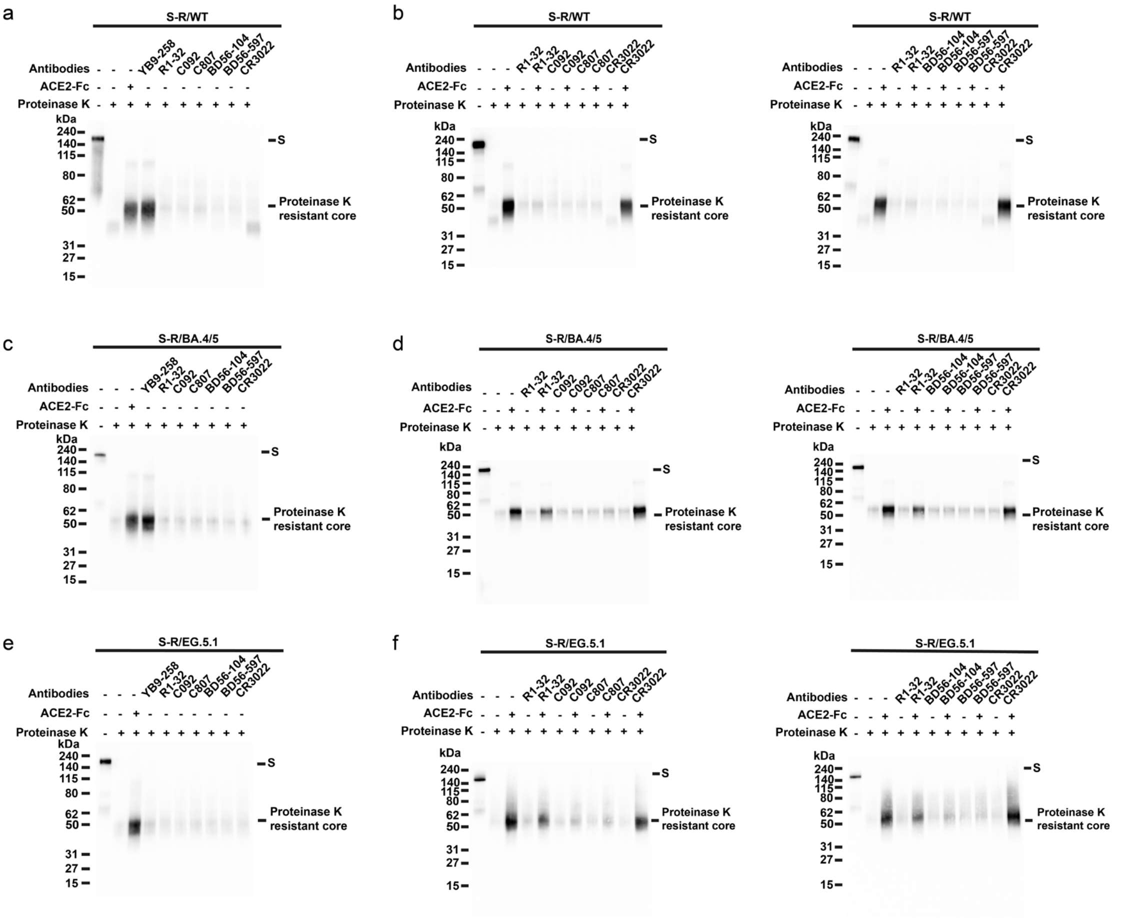


**Fig. S4. Mutation-tolerant antibodies inhibit the conformational change of SARS-CoV-2 S-trimers.**

**a-b**, Inhibition effect of R1-32-like antibodies on the conformational change of the SARS-CoV-2 wild-type spike (S-R/WT). S-R/WT (7.09 µM) was incubated with ACE2-Fc or antibodies (S-protomer:ACE2-Fc/IgG = 1:1.1 molar ratio) for 1 h (**a**). Samples were analyzed by western blotting to detect the generation of the 55 kDa proteinase K-resistant core, which is a signature of the post-fusion S2 structure. Only ACE2-Fc and class 1 antibody YB9-258 induced post-fusion structures. Inhibition of conformational change by non-ACE2 competing antibodies was assayed by antibody preincubation (1 h) before further incubation (1 h) with ACE2-Fc (**b**). Only R1-32-like antibodies were able to abolish fusogenic spike conformational change. **c-d**, Inhibition effect of R1-32-like antibodies on the conformational change of the SARS-CoV-2 BA.4/5 S-trimer (S-R/BA.4/5). **e-f**, Inhibition effect of R1-32-like antibodies on the conformational change of the SARS-CoV-2 EG.5.1 S-trimer (S-R/EG.5.1).

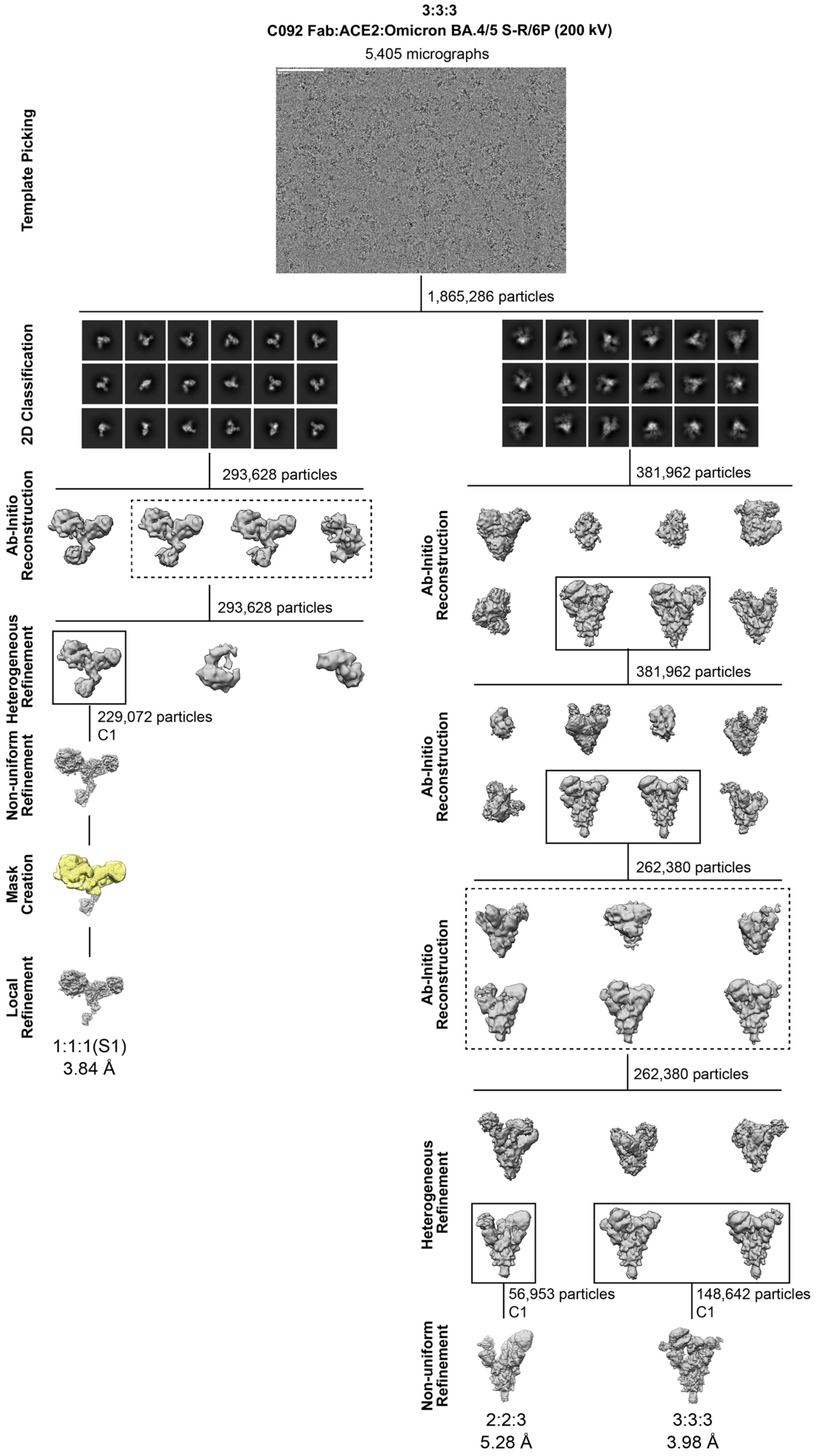


**Fig. S5. Cryo-EM data processing for the C092 Fab:ACE2:BA.4/5 S-R/6P dataset.**

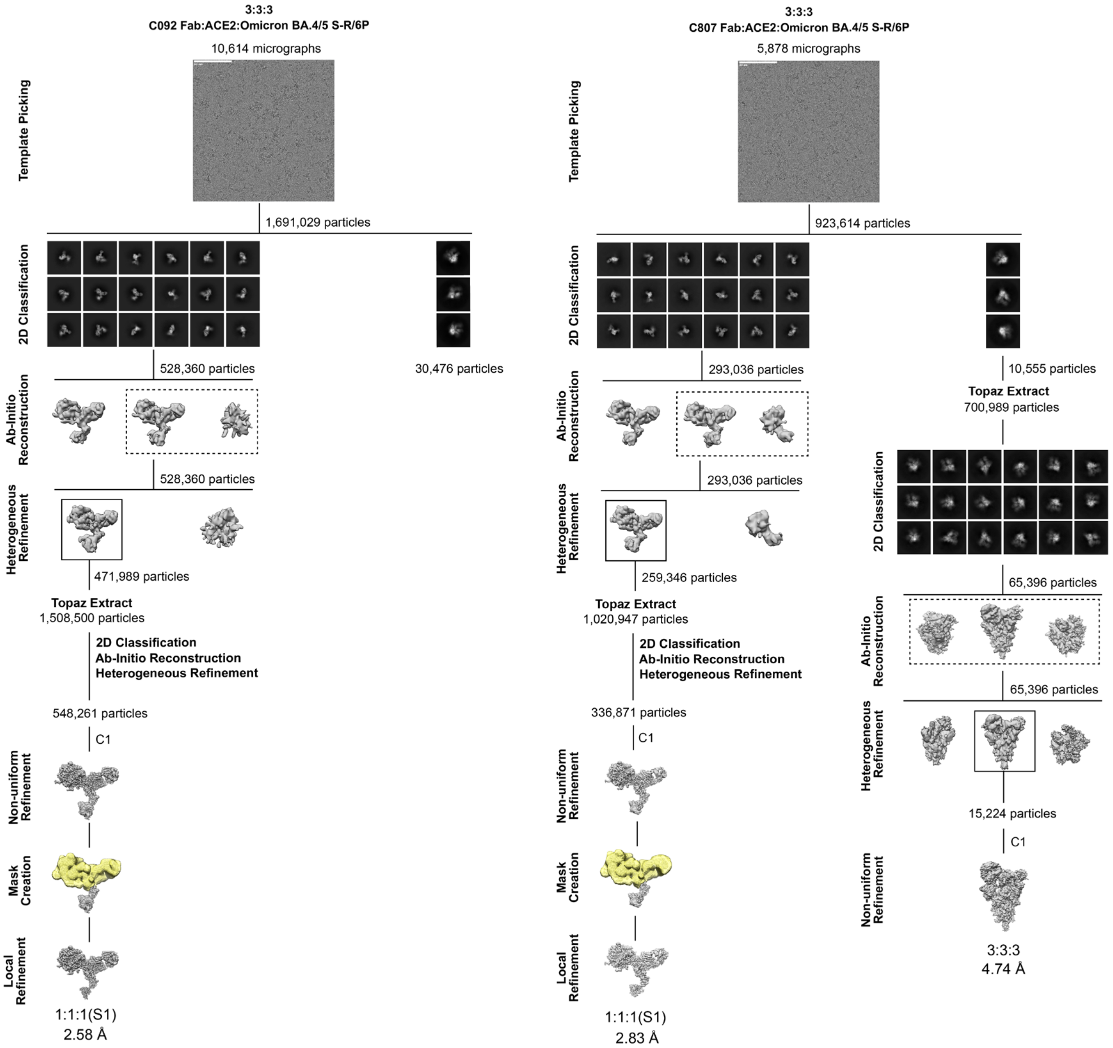


**Fig. S6. Cryo-EM data processing for the C092 Fab:ACE2:BA.4/5 S-R/6P and the C807 Fab:ACE2:BA.4/5 S-R/6P datasets.**

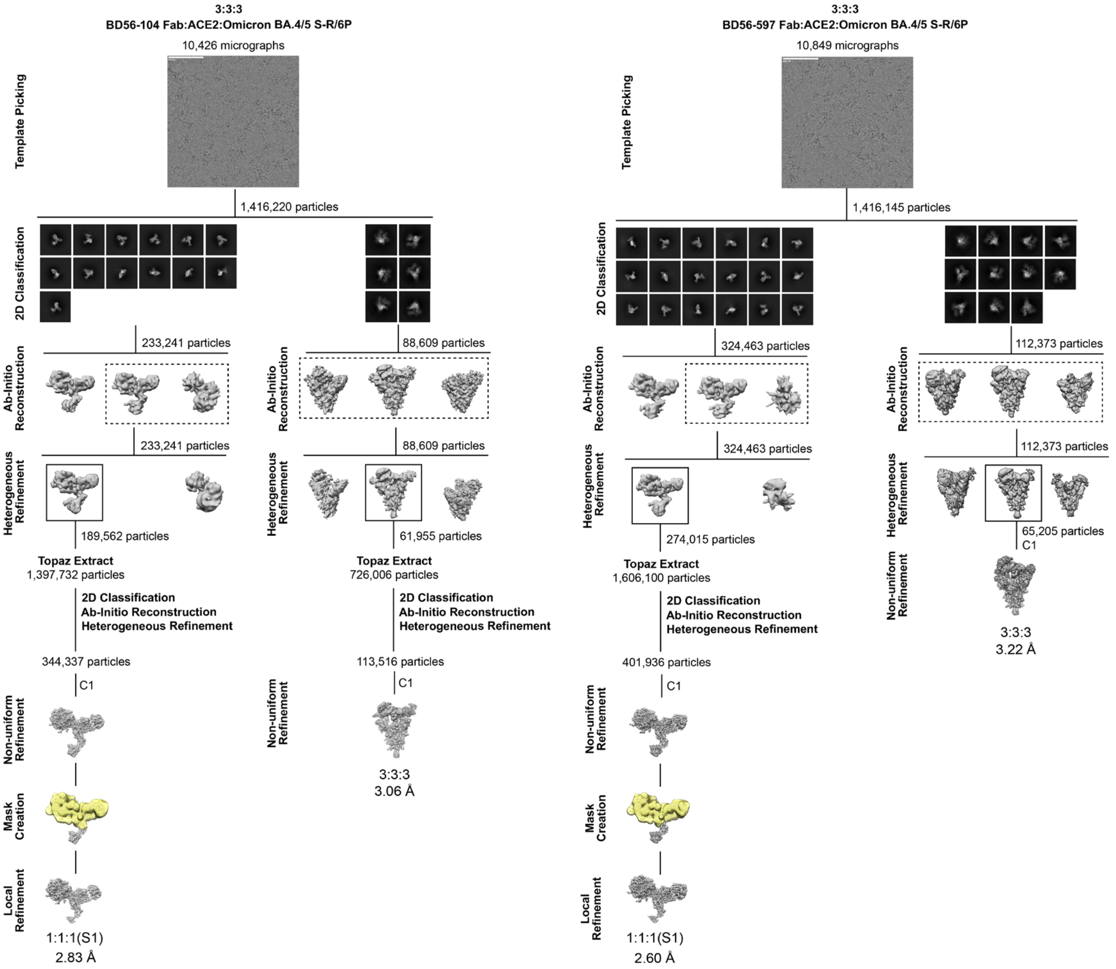


**Fig. S7. Cryo-EM data processing for the BD56-104 Fab:ACE2:BA.4/5 S-R/6P and the BD56-597 Fab:ACE2:BA.4/5 S-R/6P datasets.**

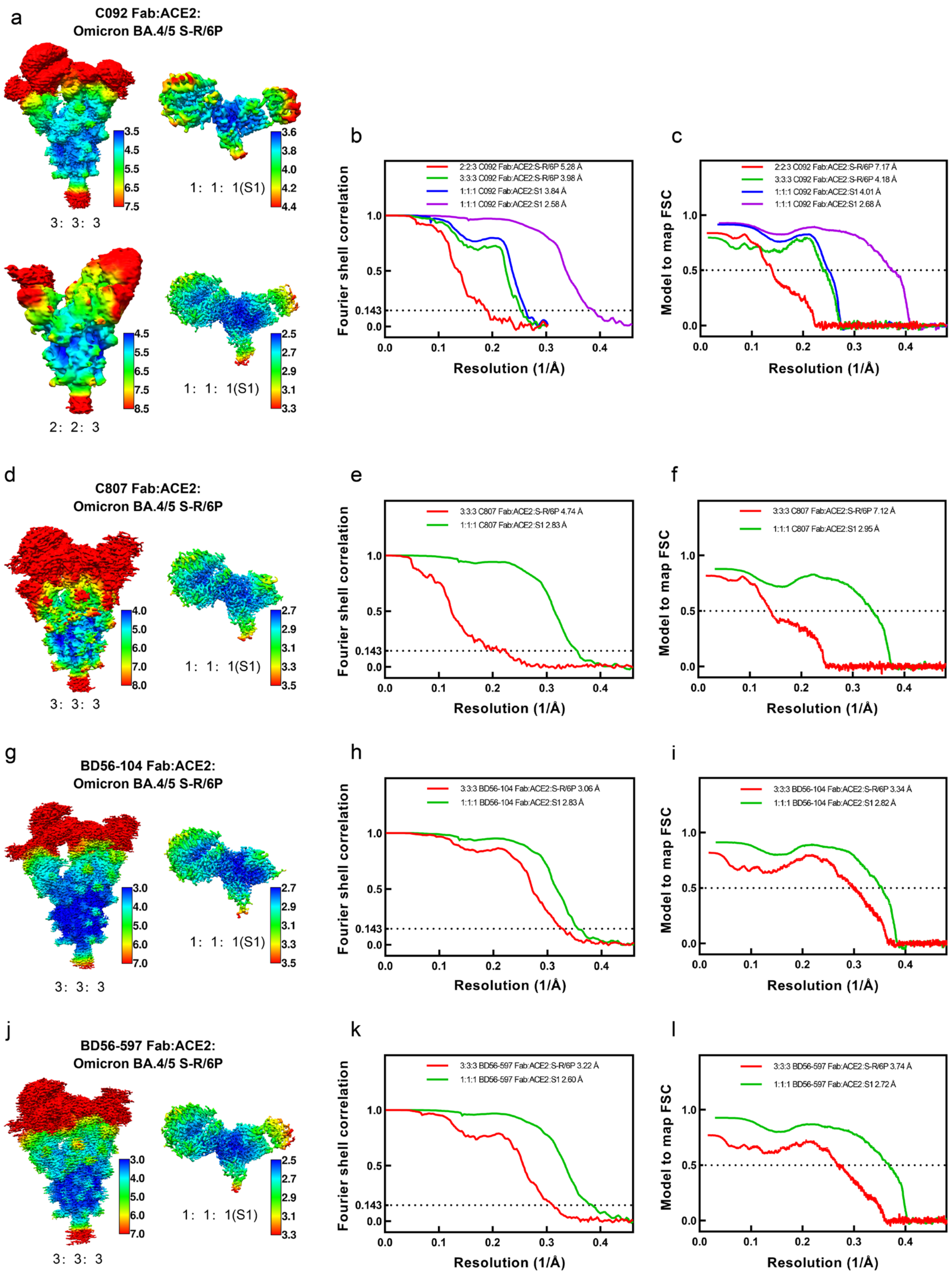


**Fig. S8. Resolution assessment of cryo-EM structures.**

**a, d, g, and j**, Local resolution maps of C092 Fab (**a**), C807 Fab (**d**), BD56-104 Fab (**g**), and BD56-597 Fab (**j**) in complex with ACE2 and Omicron BA.4/5 S-R/6P. **b, e, h, and k**, Global resolution assessment by Fourier shell correlation (FSC) at the 0.143 criterion. **c, f, i, and l**, Correlations of model vs map by FSC at the 0.5 criterion.

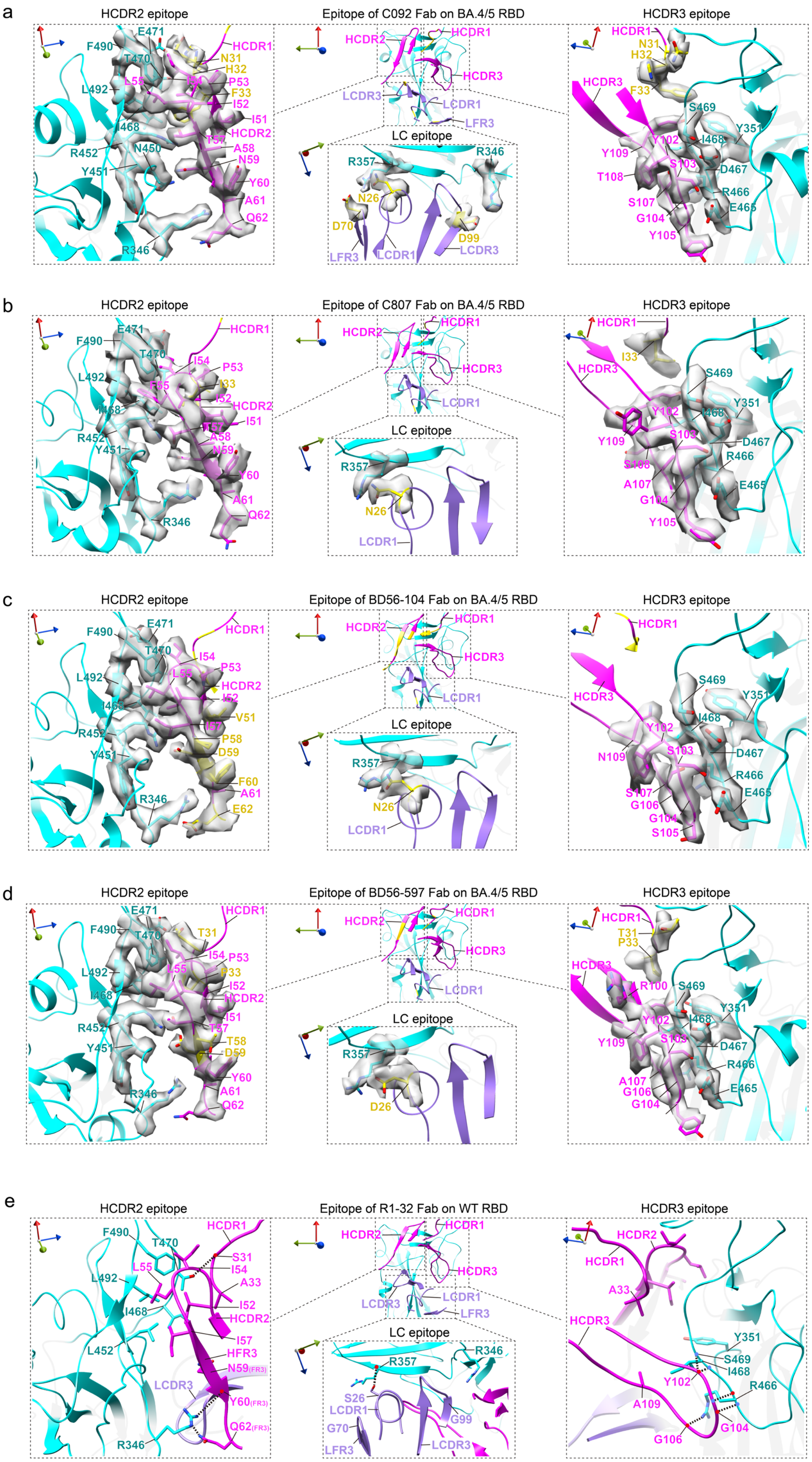

a
HCDR2 epitope
Epitope of C092 Fab on BA.4/5 RBD
HCDR3 epitope
LC epitope
F490
E471
T470
L492
I468
R452
N450
Y451
R346
HCDR1
N31
H32
P53
F33
I52
I51
HCDR2
T57
A58
N59
Y60
A61
Q62
L55
I54
HCDR3
LCDR3
LCDR1
LFR3
R357
R346
N26
D70
D99
S469
Y351
I468
D467
R466
E465
Y102
S103
Y109
T108
S107
G104
Y105
b
HCDR2 epitope
Epitope of C807 Fab on BA.4/5 RBD
HCDR3 epitope
LC epitope
F490
E471
T470
L492
I468
R452
Y451
R346
HCDR1
I54
P53
L55
I33
I52
HCDR2
I51
T57
A58
N59
Y60
A61
Q62
HCDR3
LCDR1
R357
N26
I33
S469
Y351
I468
Y102
S103
D467
R466
E465
Y109
S108
A107
G104
Y105
c
HCDR2 epitope
Epitope of BD56-104 Fab on BA.4/5 RBD
HCDR3 epitope
LC epitope
F490
E471
T470
L492
I468
R452
Y451
R346
HCDR1
I54
P53
L55
HCDR2
I52
V51
T57
P58
D59
F60
A61
E62
HCDR3
LCDR1
R357
N26
S469
Y351
I468
Y102
S103
N109
D467
R466
E465
S107
G106
G104
S105
d
HCDR2 epitope
Epitope of BD56-597 Fab on BA.4/5 RBD
HCDR3 epitope
LC epitope
E471
F490
T470
T31
L492
I468
R452
Y451
R346
HCDR1
I54
P53
L55
P33
I52
HCDR2
I51
T57
T58
D59
Y60
A61
Q62
HCDR3
LCDR1
R357
D26
T31
P33
R100
S469
Y351
Y102
I468
S103
Y109
D467
R466
E465
A107
G106
G104
e
HCDR2 epitope
Epitope of R1-32 Fab on WT RBD
HCDR3 epitope
LC epitope
F490
T470
HCDR1
L55
S31
I54
A33
L492
I52
I468
HCDR2
I57
L452
HFR3
N59(FR3)
LCDR3
Y60(FR3)
Q62(FR3)
R346
HCDR3
LCDR3
LCDR1
LFR3
R346
R357
S26
G70
G99
A33
Y351
S469
I468
Y102
R466
A109
G104
G106

**Fig. S9. Representative cryo-EM densities showing epitope binding by R1-32-like antibodies.**

**a-d**, Representative densities (grey surfaces) of the C092 (**a**), C807 (**b**), BD56-104 (**c**), and BD56-597 (**d**) epitopes and CDR loops. **e**, R1-32 HCDR2, HCDR3, and LC epitopes are shown from rotated views (PDB: 7YDI) for comparison. RBD, R1-32-H, and R1-32-L residues are colored in cyan, magenta, and purple, respectively. The backbone carbonyl oxygens and amide nitrogens are indicated by red and blue dots, respectively. Hydrogen bonds are shown as black dash lines.

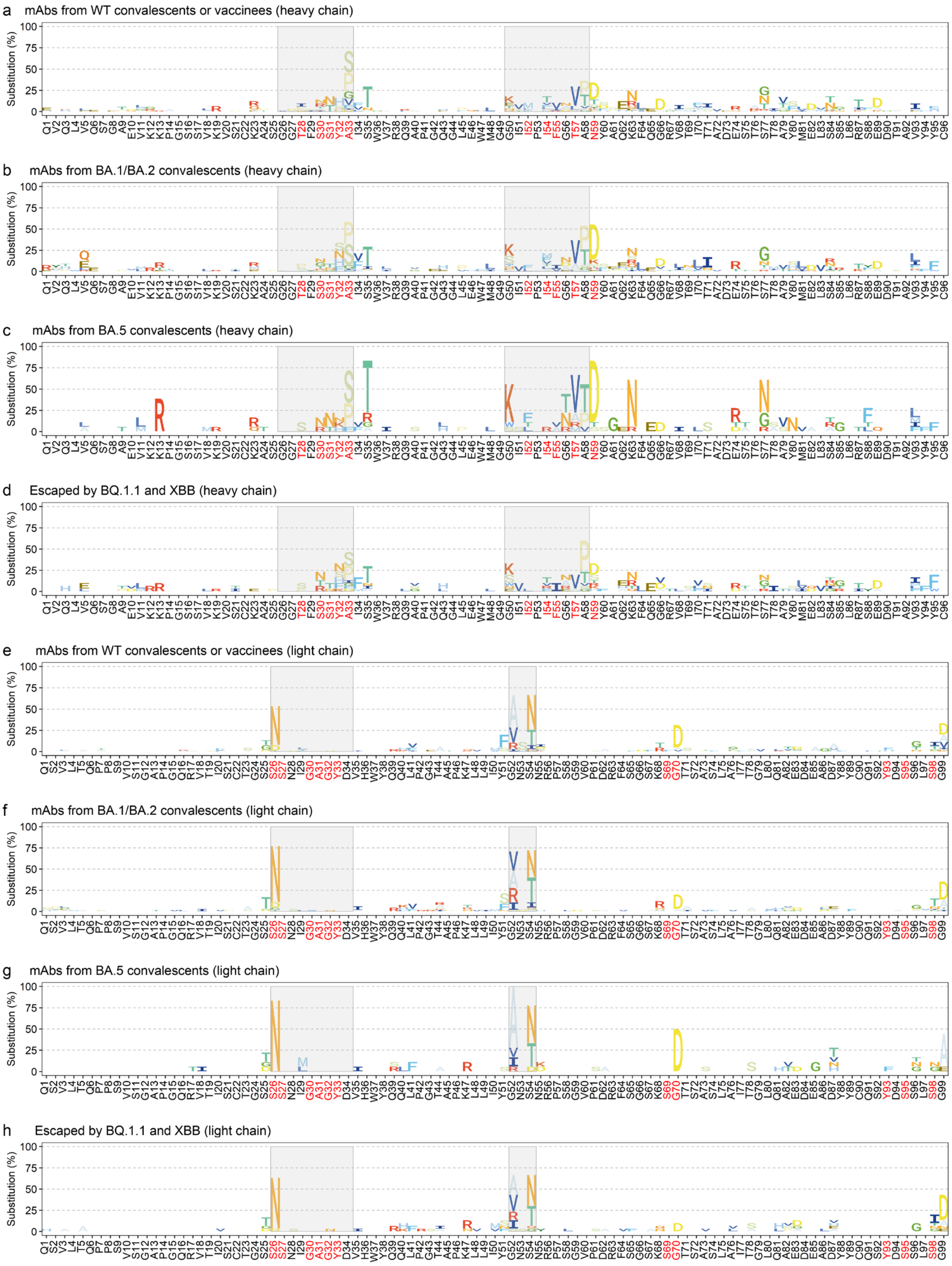


**Fig. S10. Frequency of somatic hypermutations (SHMs) in antibodies isolated from different exposure histories.**

**a-c**, Frequency of SHMs in the heavy chains of antibodies isolated from WT convalescents or vaccinees (**a**), BA.1/BA.2 convalescents (**b**), and BA.5 convalescents (**c**). **d**, Frequency of SHMs in the heavy chains of antibodies escaped by BQ.1.1 and XBB variants. **e-g**, Frequency of SHMs in the light chains of antibodies isolated from WT convalescents or vaccinees (**e**), BA.1/BA.2 convalescents (**f**), and BA.5 convalescents (**g**). **h**, Frequency of SHMs in the light chains of antibodies escaped by BQ.1.1 and XBB variants.

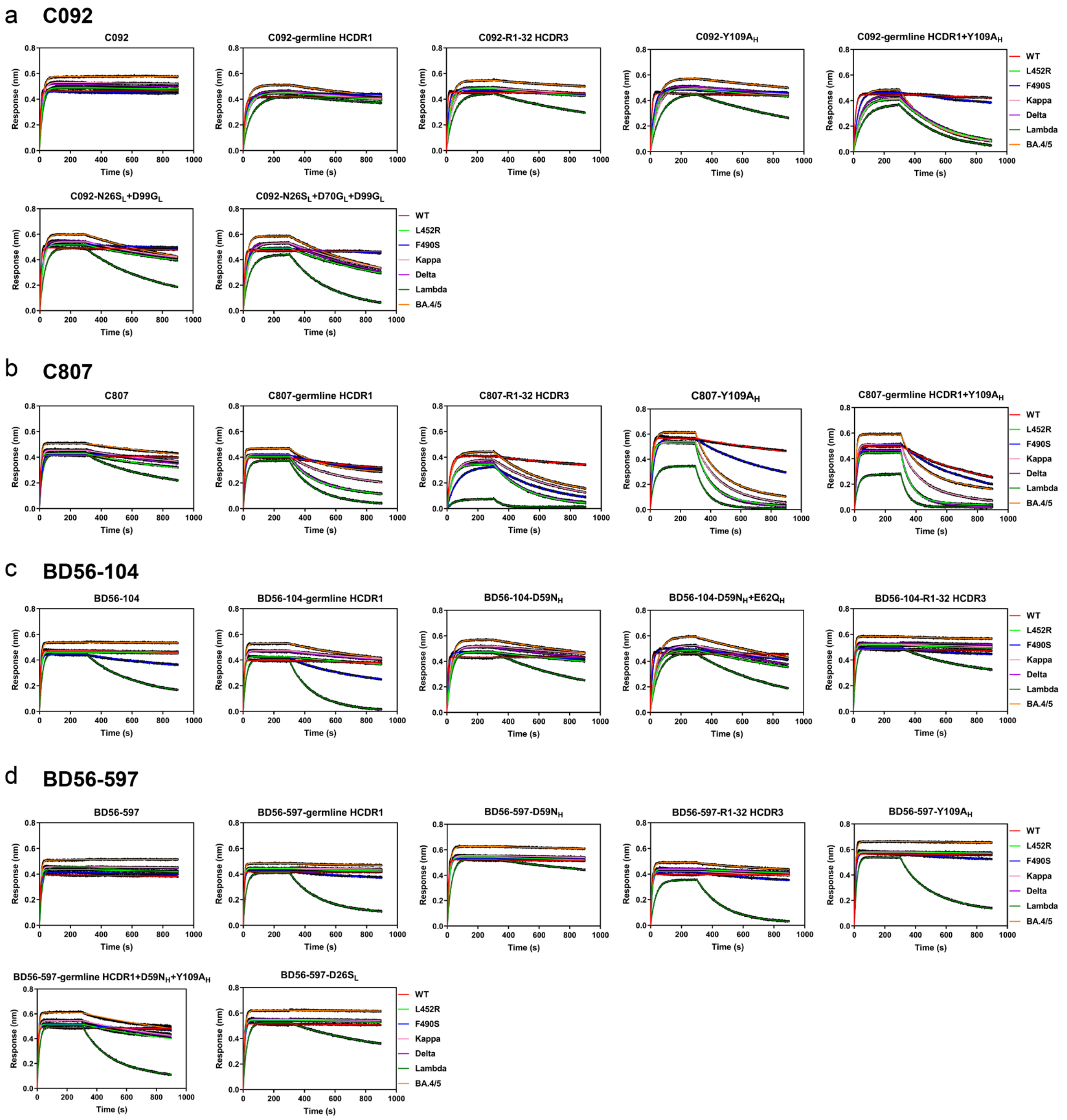


**Fig. S11. Binding assays of constructed R1-32-like antibody mutants to the RBDs.** **a-d**, Binding curves of mutants derived from C092 (**a**), C807 (**b**), BD56-104 (**c**), and BD56-597 (**d**) to the RBDs of SARS-CoV-2 wild-type, L452R, F490S, Kappa, Delta, Lambda and BA.4/5. Binding assays were performed by BLI, with detailed binding kinetics parameters summarized in **Table S3**. Binding affinity changes to different mutants are shown in **Fig. 5b-e**.

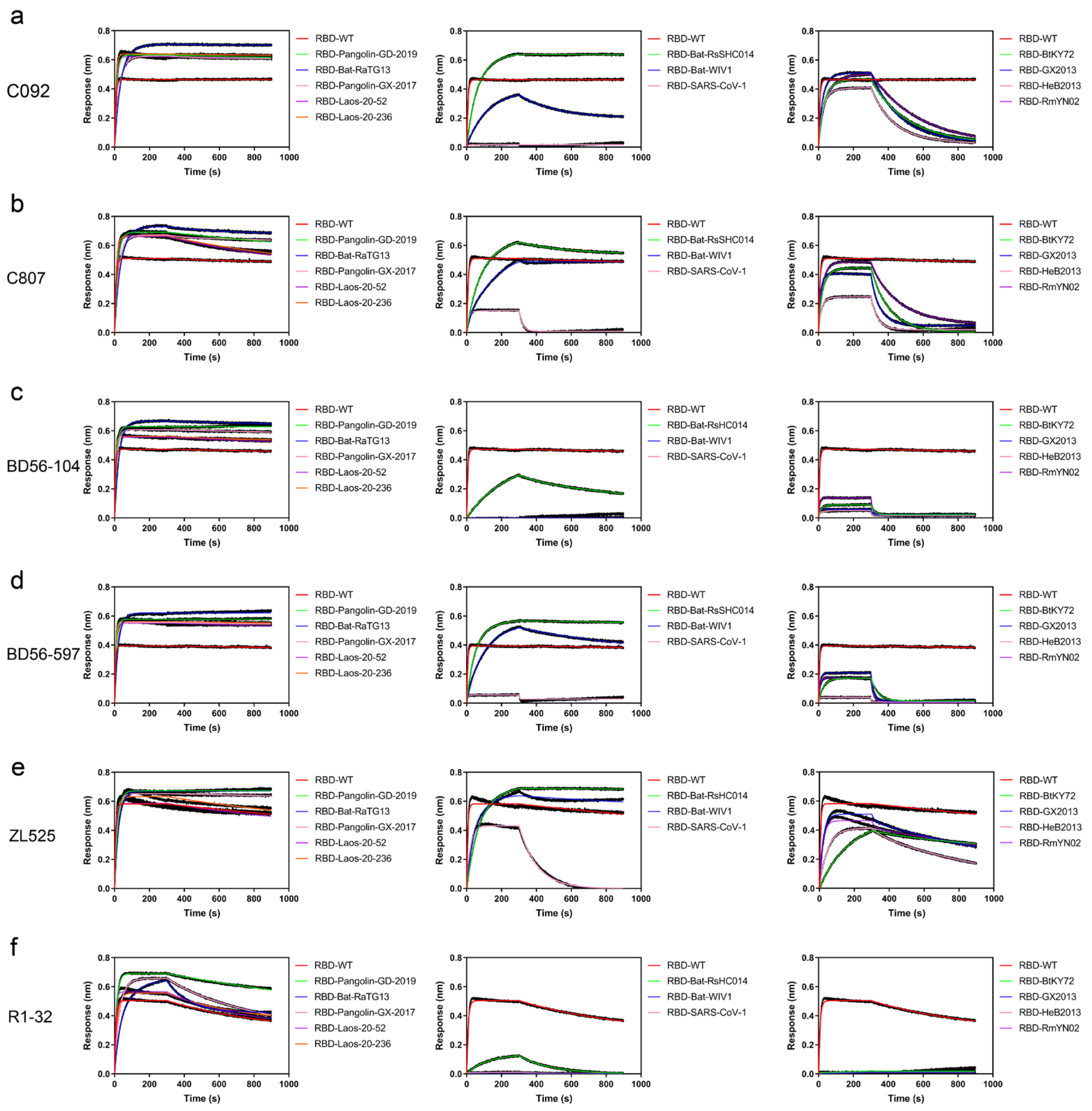


**Fig. S12. Binding of mutation-tolerant antibodies to the RBDs of SARSr-CoVs.**
**a-f**, Binding curves of C092 (**a**), C807 (**b**), BD56-104 (**c**), BD56-597 (**d**), ZL525 (**e**), and R1-32 (**f**) to the RBDs of SARS-related coronaviruses (SARSr-CoVs). Binding assays were performed by BLI, with RBDs at a fixed concentration of 200 nM. The $K_D$ values are shown in **Fig. 5a**. Detailed binding kinetic parameters are summarized in **Table S5**.

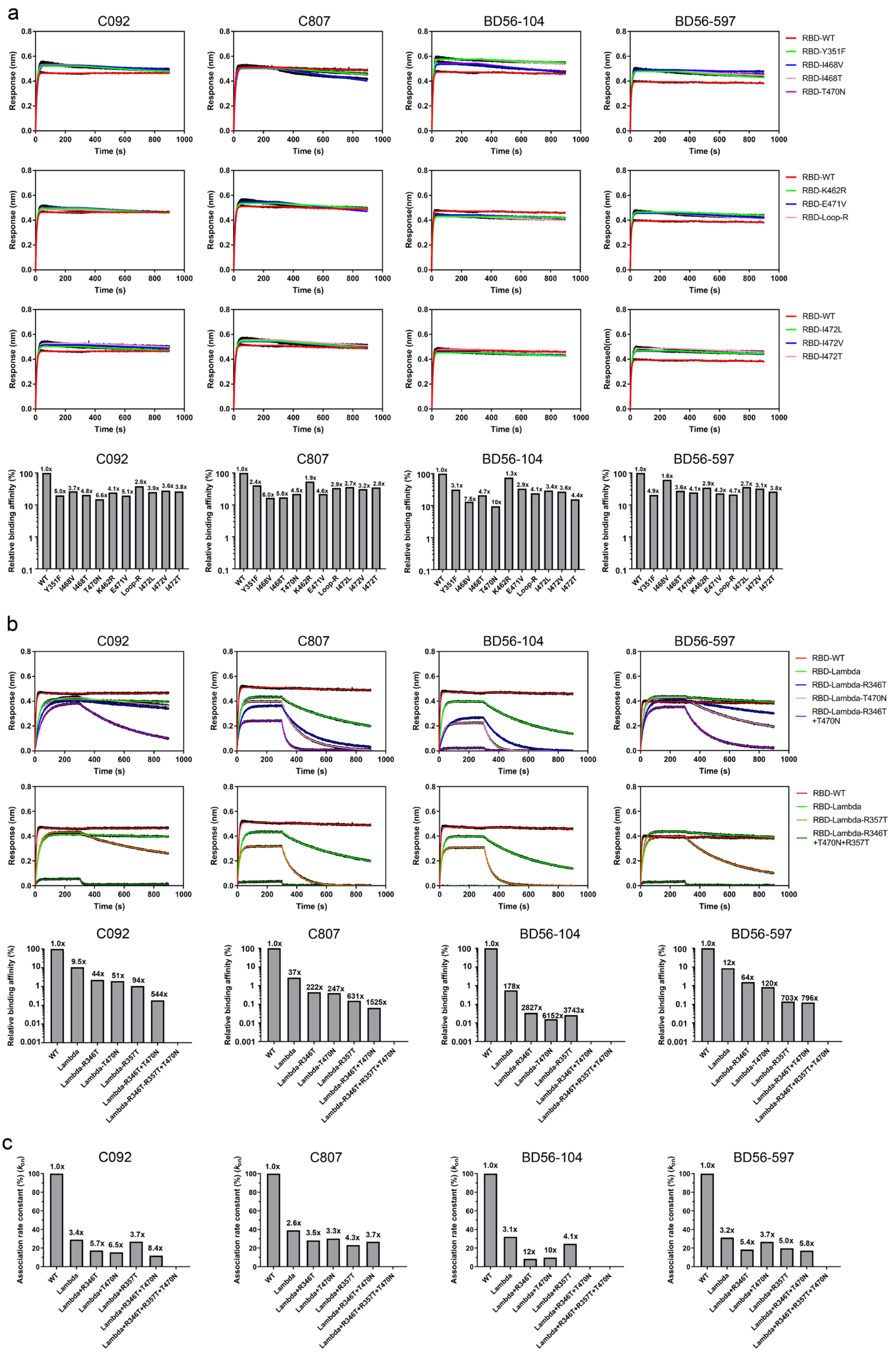

a
C092
C807
BD56-104
BD56-597
Response (nm)
Time (s)
RBD-WT
RBD-Y351F
RBD-I468V
RBD-I468T
RBD-T470N
RBD-WT
RBD-K462R
RBD-E471V
RBD-Loop-R
RBD-WT
RBD-I472L
RBD-I472V
RBD-I472T
Relative binding affinity (%)
WT
Y351F
I468V
I468T
T470N
K462R
E471V
Loop-R
I472L
I472V
I472T
b
RBD-WT
RBD-Lambda
RBD-Lambda-R346T
RBD-Lambda-T470N
RBD-Lambda-R346T +T470N
RBD-WT
RBD-Lambda
RBD-Lambda-R357T
RBD-Lambda-R346T +T470N+R357T
WT
Lambda
Lambda-R346T
Lambda-T470N
Lambda-R357T
Lambda-R346T+T470N
Lambda-R346T+R357T+T470N
c
Association rate constant (%) ($k_{on}$)
WT
Lambda
Lambda+R346T
Lambda+T470N
Lambda+R357T
Lambda+R346T+T470N
Lambda+R346T+R357T+T470N

**Fig. S13. RBD mutations and their effects on the binding of mutation-tolerant antibodies.**

**a**, Binding of R1-32-like antibodies to RBDs with single amino acid substitutions. The Loop-R indicates the replacement of the SARS-CoV-2 ACE2 binding loop (474-486) with the corresponding SARS-CoV-1 residues (461-472). Bottom panels: Relative affinity changes are normalized to the $K_D$s for wild-type RBD (set as 100%) and shown as bar graphs. Fold changes in $K_D$ relative to the wild-type RBD are indicated above the bars. **b**, Binding affinities of R1-32-like antibodies to single and combination mutations constructed on the Lambda RBD. Bottom panels: Relative affinity changes are shown as bar graphs. Fold changes in $K_D$ are indicated above the bars. **c**, Relative changes in association rate constant ($k_{on}$) are normalized to the $k_{on}$ for wild-type RBD (set as 100%) and are shown as bar graphs. Fold changes in $k_{on}$ are indicated above the bars. $K_D$ and $k_{on}$ values derived from fitting BLI binding curves in (**a**) and (**b**) are summarized in **Tables S6 and S7**.

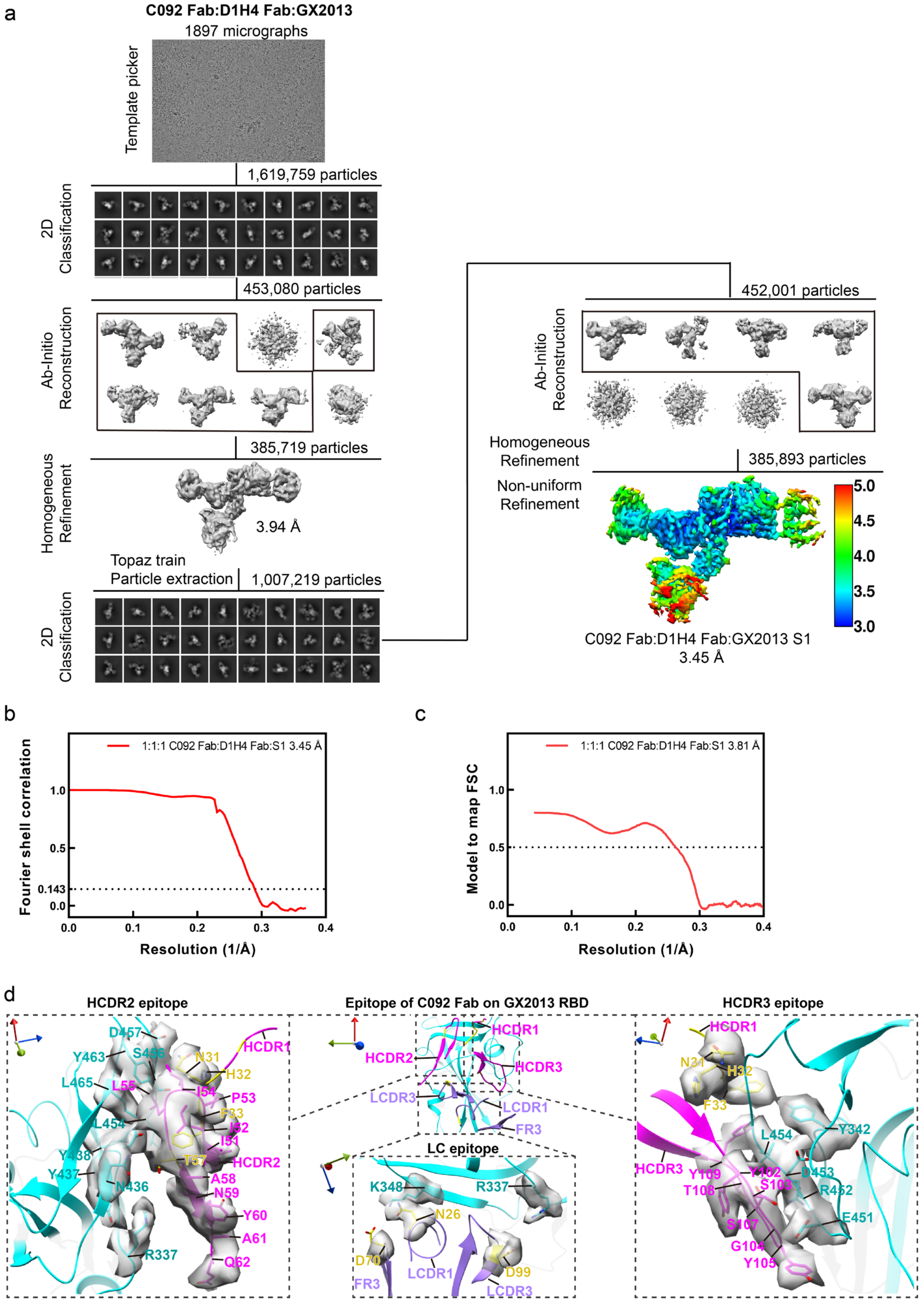


**Fig. S14. Cryo-EM structure determination of the C092 Fab:D1H4 Fab:GX2013 S1 complex.**

**a**, Cryo-EM data processing pipeline for the C092 Fab:D1H4 Fab:GX2013/5P dataset and local resolution assessment of the C092 Fab:D1H4 Fab:GX2013 S1 complex strucuture. **b**, Global resolution assessment by Fourier shell correlation (FSC) at the 0.143 criterion. **c**,

Correlations of model vs map by FSC at the 0.5 criterion. **d**, Representative cryo-EM densities showing the C092 CDR loops and the C092 epitope on the GX2013 RBD.

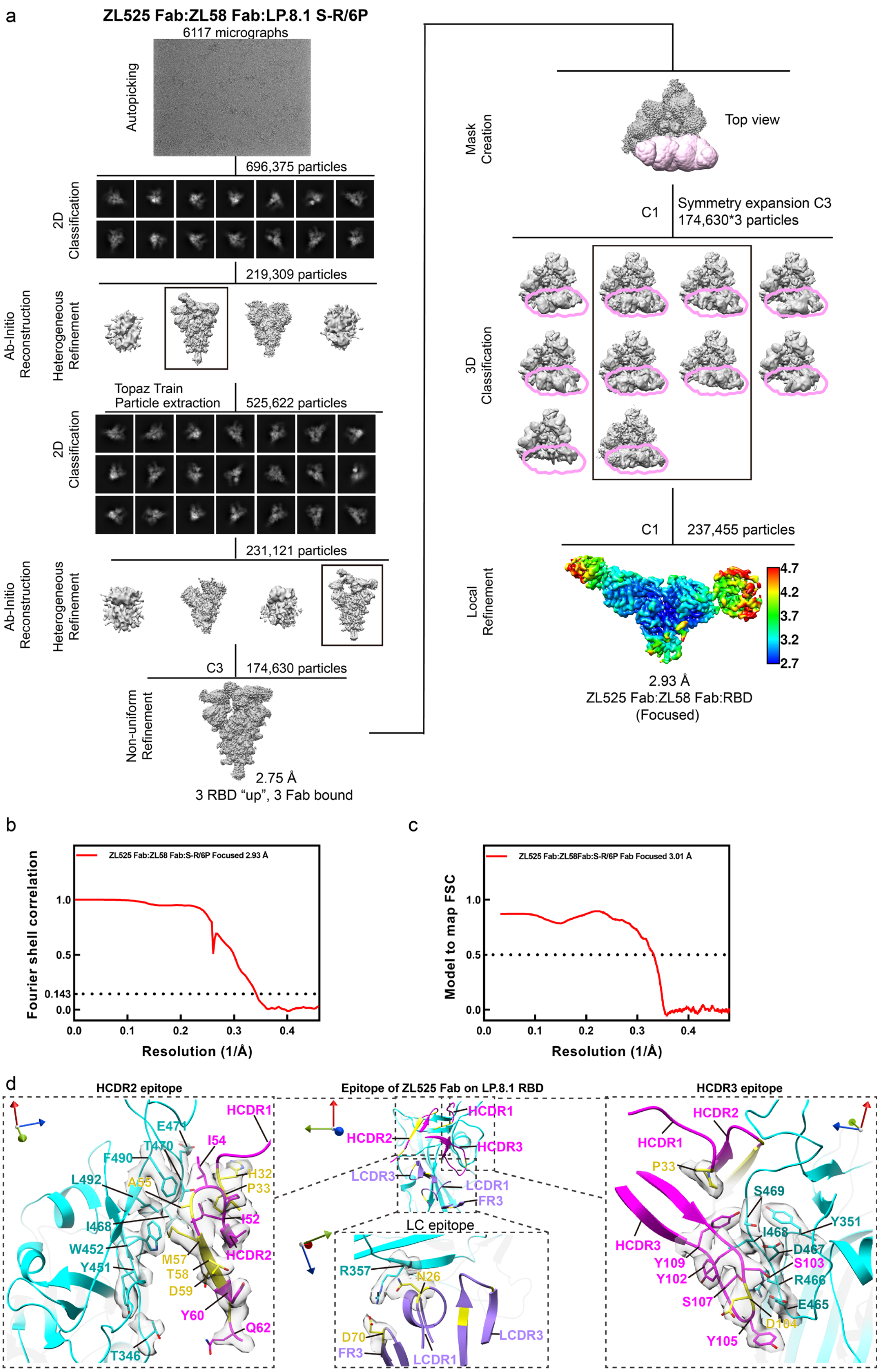
a
ZL525 Fab:ZL58 Fab:LP.8.1 S-R/6P
6117 micrographs
Autopicking
696,375 particles
2D Classification
219,309 particles
Ab-Initio Reconstruction
Heterogeneous Refinement
Topaz Train
Particle extraction
525,622 particles
2D Classification
231,121 particles
Ab-Initio Reconstruction
Heterogeneous Refinement
C3
174,630 particles
Non-uniform Refinement
2.75 Å
3 RBD "up", 3 Fab bound
Mask Creation
Top view
C1
Symmetry expansion C3
174,630*3 particles
3D Classification
C1
237,455 particles
Local Refinement
4.7
4.2
3.7
3.2
2.7
2.93 Å
ZL525 Fab:ZL58 Fab:RBD
(Focused)
b
ZL525 Fab:ZL58 Fab:S-R/6P Focused 2.93 Å
Fourier shell correlation
1.0
0.5
0.143
0.0
0.0
0.1
0.2
0.3
0.4
Resolution (1/Å)
c
ZL525 Fab:ZL58Fab:S-R/6P Fab Focused 3.01 Å
Model to map FSC
1.0
0.5
0.0
0.0
0.1
0.2
0.3
0.4
Resolution (1/Å)
d
HCDR2 epitope
HCDR1
E471
T470
I54
F490
H32
L492
A55
P33
I52
I468
W452
Y451
M57
T58
HCDR2
D59
Y60
Q62
T346
Epitope of ZL525 Fab on LP.8.1 RBD
HCDR1
HCDR2
HCDR3
LCDR3
LCDR1
FR3
LC epitope
R357
N26
D70
FR3
LCDR1
LCDR3
HCDR3 epitope
HCDR2
HCDR1
P33
S469
Y351
I468
HCDR3
D467
Y109
S103
Y102
R466
S107
E465
D104
Y105

**Fig. S15. Cryo-EM structure determination of the ZL525 Fab:ZL58 Fab:LP.8.1 S-R/6P complex.**

**a**, Cryo-EM data processing pipelines for the ZL525 Fab:ZL58 Fab:LP.8.1 S-R/6P dataset and local resolution assessment of the ZL525 Fab:ZL58 Fab:LP.8.1 RBD complex structure. **b**, Global resolution assessment by Fourier shell correlation (FSC) at the 0.143 criterion. **c**, Correlations of model vs map by FSC at the 0.5 criterion. **d**, Representative cryo-EM densities showing the ZL525 CDR loops and epitope.

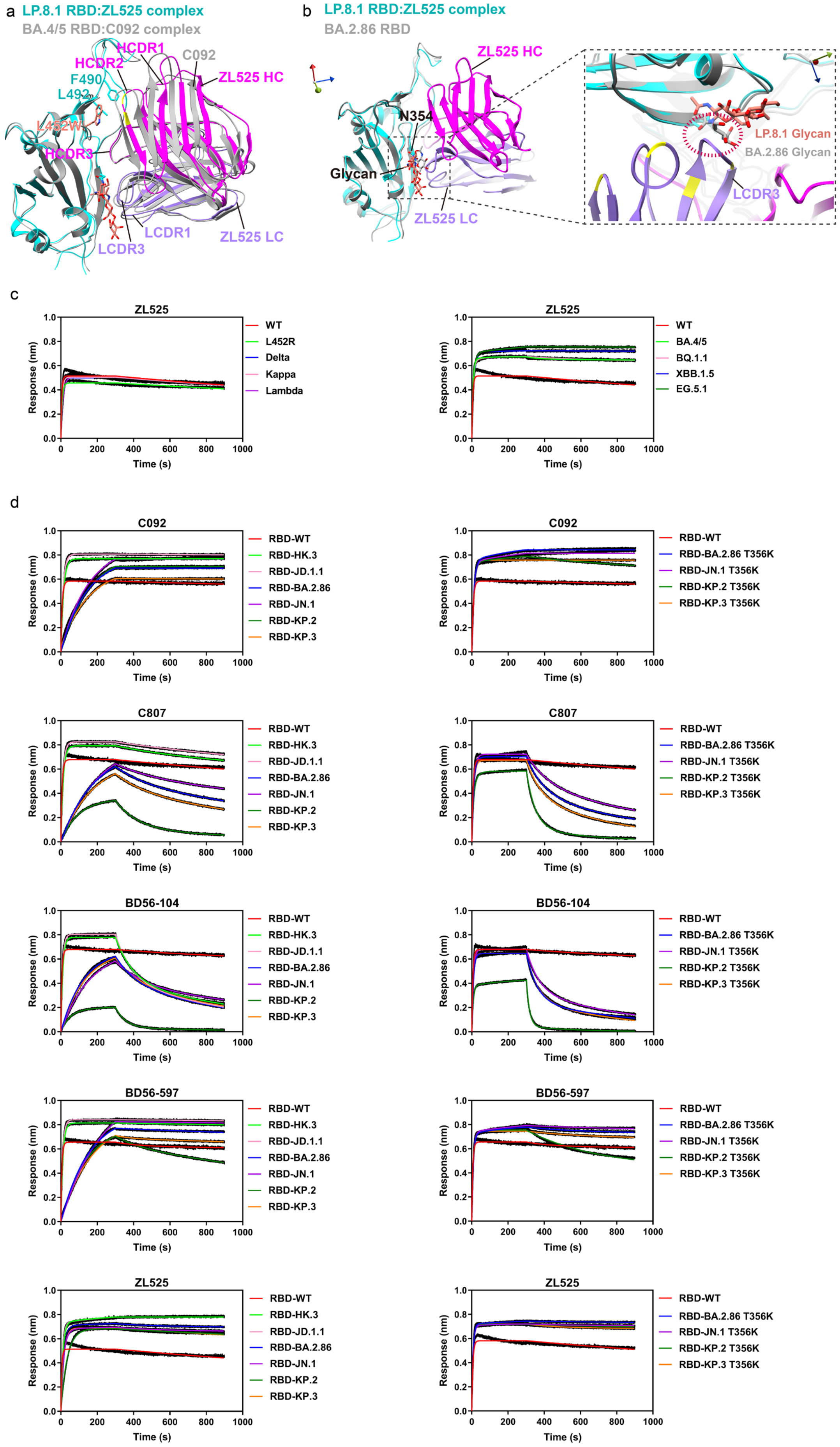
a
LP.8.1 RBD:ZL525 complex
BA.4/5 RBD:C092 complex
HCDR1
HCDR2
C092
ZL525 HC
F490
L492
L452W
HCDR3
LCDR1
ZL525 LC
LCDR3
b
LP.8.1 RBD:ZL525 complex
BA.2.86 RBD
ZL525 HC
N354
Glycan
ZL525 LC
LP.8.1 Glycan
BA.2.86 Glycan
LCDR3
c
ZL525
WT
L452R
Delta
Kappa
Lambda
BA.4/5
BQ.1.1
XBB.1.5
EG.5.1
Response (nm)
Time (s)
d
C092
C807
BD56-104
BD56-597
ZL525
RBD-WT
RBD-HK.3
RBD-JD.1.1
RBD-BA.2.86
RBD-JN.1
RBD-KP.2
RBD-KP.3
RBD-BA.2.86 T356K
RBD-JN.1 T356K
RBD-KP.2 T356K
RBD-KP.3 T356K
Response (nm)
Time (s)

**Fig. S16. Structural and functional characterization of N354 glycosylation on SARS-CoV-2 RBD.**

**a**, Structural superposition of the LP.8.1 RBD:ZL525 Fab and BA.4/5 RBD:C092 Fab complexes, using the LP.8.1 RBD as the reference. The LP.8.1 RBD:ZL525 Fab complex is shown in color, and the BA.4/5 RBD:C092 Fab complex is depicted in grey. **b,** Mapping of the N354 glycosylation site on the RBD. The structure of the BA.2.86 RBD is colored grey. **c**, Binding curves of ZL525 to a panel of SARS-CoV-2 RBDs. The $K_D$ values and normalized $K_D$ fold changes are shown in **Fig. 6e**. **d**, Binding curves of C092, C807, BD56-104, BD56-597, and ZL525 against RBDs from newly emergent variants and engineered T356K mutants. Binding assays were performed by BLI. Detailed binding kinetic parameters are summarized in **Table S8**.

### Table S1. Kinetic parameters of antibody binding to variant RBDs (related to Fig. S1).

| | **C092** | | | **C807** | | | **BD56-104** | | | **BD56-597** | | |
|---|---|---|---|---|---|---|---|---|---|---|---|---|
| | $k_{on}$ ($M^{-1}$ $s^{-1}$) | $k_{off}$ ($s^{-1}$) | $K_D$ (nM) | $k_{on}$ ($M^{-1}$ $s^{-1}$) | $k_{off}$ ($s^{-1}$) | $K_D$ (nM) | $k_{on}$ ($M^{-1}$ $s^{-1}$) | $k_{off}$ ($s^{-1}$) | $K_D$ (nM) | $k_{on}$ ($M^{-1}$ $s^{-1}$) | $k_{off}$ ($s^{-1}$) | $K_D$ (nM) |
| WT | $7.62\times10^{5}$ | $3.53\times10^{-5}$ | 0.046 | $6.56\times10^{5}$ | $7.74\times10^{-5}$ | 0.118 | $8.62\times10^{5}$ | $3.29\times10^{-5}$ | 0.038 | $9.29\times10^{5}$ | $4.46\times10^{-5}$ | 0.048 |
| K417N | $5.91\times10^{5}$ | $5.50\times10^{-5}$ | 0.093 | $5.22\times10^{5}$ | $1.15\times10^{-4}$ | 0.221 | $8.60\times10^{5}$ | $6.31\times10^{-5}$ | 0.073 | $9.88\times10^{5}$ | $4.33\times10^{-5}$ | 0.044 |
| K417T | $7.99\times10^{5}$ | $1.55\times10^{-5}$ | 0.019 | $7.03\times10^{5}$ | $1.11\times10^{-4}$ | 0.157 | $8.98\times10^{5}$ | $8.77\times10^{-5}$ | 0.098 | $8.08\times10^{5}$ | $4.64\times10^{-5}$ | 0.057 |
| E484K | $4.68\times10^{5}$ | $1.94\times10^{-5}$ | 0.042 | $4.29\times10^{5}$ | $8.47\times10^{-5}$ | 0.198 | $7.82\times10^{5}$ | $5.50\times10^{-5}$ | 0.070 | $8.01\times10^{5}$ | $3.79\times10^{-5}$ | 0.047 |
| E484Q | $6.27\times10^{5}$ | $2.98\times10^{-5}$ | 0.048 | $5.72\times10^{5}$ | $9.58\times10^{-5}$ | 0.168 | $8.38\times10^{5}$ | $7.33\times10^{-5}$ | 0.088 | $9.38\times10^{5}$ | $5.87\times10^{-5}$ | 0.063 |
| L452R | $3.04\times10^{5}$ | $2.19\times10^{-5}$ | 0.072 | $4.75\times10^{5}$ | $5.14\times10^{-4}$ | 1.080 | $5.53\times10^{5}$ | $2.66\times10^{-5}$ | 0.048 | $6.66\times10^{5}$ | $5.29\times10^{-5}$ | 0.079 |
| L452Q | $3.30\times10^{5}$ | $6.59\times10^{-5}$ | 0.200 | $3.26\times10^{5}$ | $3.84\times10^{-4}$ | 1.180 | $4.42\times10^{5}$ | $1.54\times10^{-4}$ | 0.349 | $5.20\times10^{5}$ | $5.63\times10^{-5}$ | 0.108 |
| L452M | $8.81\times10^{5}$ | $3.50\times10^{-5}$ | 0.040 | $8.60\times10^{5}$ | $2.11\times10^{-4}$ | 0.246 | $1.20\times10^{6}$ | $3.21\times10^{-5}$ | 0.027 | $1.28\times10^{6}$ | $3.92\times10^{-5}$ | 0.031 |
| T478K | $5.48\times10^{5}$ | $3.17\times10^{-5}$ | 0.058 | $5.16\times10^{5}$ | $8.48\times10^{-5}$ | 0.164 | $6.69\times10^{5}$ | $4.91\times10^{-5}$ | 0.073 | $8.05\times10^{5}$ | $3.98\times10^{-5}$ | 0.050 |
| F490S | $3.75\times10^{5}$ | $4.78\times10^{-5}$ | 0.128 | $3.43\times10^{5}$ | $1.50\times10^{-4}$ | 0.437 | $5.90\times10^{5}$ | $3.49\times10^{-4}$ | 0.592 | $5.71\times10^{5}$ | $6.02\times10^{-5}$ | 0.105 |
| Alpha | $6.82\times10^{5}$ | $3.41\times10^{-5}$ | 0.050 | $6.95\times10^{5}$ | $9.91\times10^{-5}$ | 0.142 | $8.57\times10^{5}$ | $2.31\times10^{-5}$ | 0.027 | $1.04\times10^{6}$ | $5.17\times10^{-5}$ | 0.050 |
| Beta | $5.82\times10^{5}$ | $5.43\times10^{-5}$ | 0.093 | $5.04\times10^{5}$ | $8.07\times10^{-5}$ | 0.160 | $6.92\times10^{5}$ | $3.20\times10^{-5}$ | 0.046 | $7.24\times10^{5}$ | $1.03\times10^{-5}$ | 0.014 |
| Gamma | $6.06\times10^{5}$ | $5.81\times10^{-5}$ | 0.096 | $5.22\times10^{5}$ | $7.97\times10^{-5}$ | 0.153 | $7.67\times10^{5}$ | $2.76\times10^{-5}$ | 0.036 | $7.73\times10^{5}$ | $1.23\times10^{-5}$ | 0.016 |
| Kappa | $2.90\times10^{5}$ | $3.68\times10^{-5}$ | 0.127 | $4.18\times10^{5}$ | $3.40\times10^{-4}$ | 0.813 | $5.31\times10^{5}$ | $4.81\times10^{-5}$ | 0.091 | $6.14\times10^{5}$ | $2.27\times10^{-5}$ | 0.037 |
| Delta | $3.64\times10^{5}$ | $6.52\times10^{-5}$ | 0.179 | $5.14\times10^{5}$ | $4.65\times10^{-4}$ | 0.904 | $5.10\times10^{5}$ | $4.16\times10^{-5}$ | 0.082 | $6.22\times10^{5}$ | $2.17\times10^{-5}$ | 0.035 |
| Lambda | $2.22\times10^{5}$ | $9.67\times10^{-5}$ | 0.436 | $2.56\times10^{5}$ | $1.12\times10^{-3}$ | 4.360 | $2.77\times10^{5}$ | $1.87\times10^{-3}$ | 6.760 | $2.90\times10^{5}$ | $1.62\times10^{-4}$ | 0.560 |
| BA.1 | $2.86\times10^{5}$ | $4.06\times10^{-5}$ | 0.142 | $2.57\times10^{5}$ | $5.76\times10^{-5}$ | 0.224 | $3.60\times10^{5}$ | $4.40\times10^{-5}$ | 0.122 | $3.05\times10^{5}$ | $2.04\times10^{-5}$ | 0.067 |
| BA.1.1 | $3.75\times10^{5}$ | $5.81\times10^{-6}$ | 0.016 | $4.56\times10^{5}$ | $3.68\times10^{-5}$ | 0.081 | $8.60\times10^{5}$ | $2.03\times10^{-5}$ | 0.024 | $9.03\times10^{5}$ | $2.42\times10^{-5}$ | 0.027 |
| BA.2 | $9.16\times10^{5}$ | $2.00\times10^{-5}$ | 0.022 | $5.79\times10^{5}$ | $7.86\times10^{-6}$ | 0.014 | $1.27\times10^{6}$ | $2.65\times10^{-5}$ | 0.021 | $1.03\times10^{6}$ | $4.78\times10^{-5}$ | 0.046 |
| BA.2.11 | $6.03\times10^{5}$ | $1.95\times10^{-5}$ | 0.032 | $4.48\times10^{5}$ | $2.11\times10^{-4}$ | 0.471 | $8.73\times10^{5}$ | $4.27\times10^{-5}$ | 0.049 | $8.81\times10^{5}$ | $1.37\times10^{-5}$ | 0.016 |
| BA.2.12.1 | $7.73\times10^{5}$ | $1.65\times10^{-4}$ | 0.213 | $5.65\times10^{5}$ | $1.21\times10^{-4}$ | 0.215 | $7.25\times10^{5}$ | $6.30\times10^{-5}$ | 0.087 | $8.44\times10^{5}$ | $1.76\times10^{-5}$ | 0.021 |
| BA.2.13 | $6.95\times10^{5}$ | $1.51\times10^{-5}$ | 0.022 | $6.20\times10^{5}$ | $2.70\times10^{-5}$ | 0.044 | $1.23\times10^{6}$ | $2.50\times10^{-5}$ | 0.020 | $1.19\times10^{6}$ | $2.19\times10^{-5}$ | 0.019 |
| BA.3 | $8.79\times10^{5}$ | $2.84\times10^{-5}$ | 0.032 | $9.26\times10^{5}$ | $4.53\times10^{-5}$ | 0.049 | $1.02\times10^{6}$ | $3.17\times10^{-5}$ | 0.031 | $1.09\times10^{6}$ | $3.38\times10^{-5}$ | 0.031 |
| BA.4/5 | $3.87\times10^{5}$ | $5.12\times10^{-5}$ | 0.132 | $4.47\times10^{5}$ | $3.31\times10^{-4}$ | 0.741 | $5.63\times10^{5}$ | $1.62\times10^{-4}$ | 0.029 | $6.61\times10^{5}$ | $3.36\times10^{-5}$ | 0.051 |
| BA.2.75 | $7.48\times10^{5}$ | $1.29\times10^{-5}$ | 0.017 | $6.62\times10^{5}$ | $3.17\times10^{-5}$ | 0.048 | $8.73\times10^{5}$ | $8.66\times10^{-5}$ | 0.099 | $8.35\times10^{5}$ | $4.02\times10^{-5}$ | 0.048 |
| BA.2.75.2 | $6.77\times10^{5}$ | $5.75\times10^{-5}$ | 0.085 | $5.47\times10^{5}$ | $8.10\times10^{-5}$ | 0.148 | $6.90\times10^{5}$ | $1.15\times10^{-4}$ | 0.167 | $7.25\times10^{5}$ | $5.72\times10^{-5}$ | 0.079 |
| BF.7 | $2.53\times10^{5}$ | $7.42\times10^{-5}$ | 0.293 | $4.13\times10^{5}$ | $7.88\times10^{-4}$ | 1.910 | $3.89\times10^{5}$ | $2.32\times10^{-4}$ | 0.598 | $4.53\times10^{5}$ | $2.77\times10^{-5}$ | 0.061 |
| BQ.1.1 | $2.54\times10^{5}$ | $5.07\times10^{-5}$ | 0.200 | $3.92\times10^{5}$ | $7.86\times10^{-4}$ | 2.010 | $3.27\times10^{5}$ | $2.93\times10^{-4}$ | 0.897 | $4.78\times10^{5}$ | $4.66\times10^{-5}$ | 0.097 |
| XBB | $5.27\times10^{5}$ | $2.93\times10^{-5}$ | 0.056 | $5.74\times10^{5}$ | $1.11\times10^{-4}$ | 0.194 | $7.23\times10^{5}$ | $1.52\times10^{-3}$ | 2.110 | $6.33\times10^{5}$ | $4.43\times10^{-5}$ | 0.070 |
| XBB.1.5 | $3.63\times10^{5}$ | $2.66\times10^{-5}$ | 0.073 | $3.15\times10^{5}$ | $2.01\times10^{-4}$ | 0.640 | $4.12\times10^{5}$ | $1.79\times10^{-3}$ | 4.350 | $3.63\times10^{5}$ | $1.23\times10^{-5}$ | 0.034 |
| EG.5 | $2.99\times10^{5}$ | $1.67\times10^{-5}$ | 0.056 | $3.39\times10^{5}$ | $2.21\times10^{-4}$ | 0.653 | $4.43\times10^{5}$ | $1.88\times10^{-3}$ | 4.250 | $3.92\times10^{5}$ | $1.70\times10^{-5}$ | 0.043 |
| EG.5.1 | $3.51\times10^{5}$ | $2.07\times10^{-5}$ | 0.059 | $3.51\times10^{5}$ | $2.37\times10^{-4}$ | 0.677 | $4.59\times10^{5}$ | $2.02\times10^{-3}$ | 4.400 | $3.96\times10^{5}$ | $2.02\times10^{-5}$ | 0.051 |

### Table S2. Kinetic parameters of antibody binding to different SARS-CoV-2 S-proteins (related to Fig. S2).

| | C092 | | | C807 | | | BD56-104 | | | BD56-597 | | |
|---|---|---|---|---|---|---|---|---|---|---|---|---|
| | $k_{on}$ ($M^{-1}$ $s^{-1}$) | $k_{off}$ ($s^{-1}$) | $K_D$ (nM) | $k_{on}$ ($M^{-1}$ $s^{-1}$) | $k_{off}$ ($s^{-1}$) | $K_D$ (nM) | $k_{on}$ ($M^{-1}$ $s^{-1}$) | $k_{off}$ ($s^{-1}$) | $K_D$ (nM) | $k_{on}$ ($M^{-1}$ $s^{-1}$) | $k_{off}$ ($s^{-1}$) | $K_D$ (nM) |
| **S-R/WT** | $2.65\times10^{4}$ ($k_{on1}$)<br>$1.57\times10^{5}$ ($k_{on2}$) | $3.13\times10^{-7}$ ($k_{off1}$)<br>$< 1.00\times10^{-7}$ ($k_{off2}$) | $\leqslant 0.0118$ ($k_{off1}/k_{on1}$) | $3.24\times10^{4}$ ($k_{on1}$)<br>$1.46\times10^{5}$ ($k_{on2}$) | $1.09\times10^{-5}$ ($k_{off1}$)<br>$< 1.00\times10^{-7}$ ($k_{off2}$) | 0.335 ($k_{off1}/k_{on1}$)<br>$\leqslant 0.0744$ ($k_{off1}/k_{on2}$) | $3.08\times10^{4}$ ($k_{on1}$)<br>$1.37\times10^{5}$ ($k_{on2}$) | $1.97\times10^{-5}$ ($k_{off1}$)<br>$< 1.00\times10^{-7}$ ($k_{off2}$) | 0.640 ($k_{off1}/k_{on1}$)<br>0.144 ($k_{off1}/k_{on2}$)<br>$< 0.00325$ ($k_{off2}/k_{on1}$) | $2.70\times10^{4}$ ($k_{on1}$)<br>$1.26\times10^{5}$ ($k_{on2}$) | $2.22\times10^{-5}$ ($k_{off1}$)<br>$3.86\times10^{-6}$ ($k_{off2}$) | 0.820 ($k_{off1}/k_{on1}$)<br>0.177 ($k_{off1}/k_{on2}$)<br>0.143 ($k_{off2}/k_{on1}$)<br>0.0307 ($k_{off2}/k_{on2}$) |
| **S-R/Delta** | $2.86\times10^{4}$ ($k_{on1}$)<br>$1.83\times10^{5}$ ($k_{on2}$) | $5.90\times10^{-5}$ ($k_{off1}$)<br>$8.84\times10^{-6}$ ($k_{off2}$) | 2.06 ($k_{off1}/k_{on1}$)<br>0.323 ($k_{off1}/k_{on2}$)<br>0.309 ($k_{off2}/k_{on1}$)<br>0.0484 ($k_{off2}/k_{on2}$) | $3.77\times10^{4}$ ($k_{on1}$)<br>$3.27\times10^{5}$ ($k_{on2}$) | $1.85\times10^{-5}$ ($k_{off1}$)<br>$8.10\times10^{-7}$ ($k_{off2}$) | 0.490 ($k_{off1}/k_{on1}$)<br>$\leqslant 0.0564$ ($k_{off1}/k_{on2}$) | $3.02\times10^{4}$ ($k_{on1}$)<br>$2.54\times10^{5}$ ($k_{on2}$) | $1.42\times10^{-6}$ ($k_{off1}$)<br>$5.23\times10^{-7}$ ($k_{off2}$) | $\leqslant 0.047$ ($k_{off1}/k_{on1}$) | $2.92\times10^{4}$ ($k_{on1}$)<br>$2.57\times10^{5}$ ($k_{on2}$) | $3.08\times10^{-7}$ ($k_{off1}$)<br>$2.57\times10^{-7}$ ($k_{off2}$) | $\leqslant 0.0105$ ($k_{off1}/k_{on1}$) |
| **S-R/Lambda** | $1.20\times10^{4}$ ($k_{on1}$)<br>$7.11\times10^{4}$ ($k_{on2}$) | $1.49\times10^{-5}$ ($k_{off1}$)<br>$< 1.00\times10^{-7}$ ($k_{off2}$) | 1.25 ($k_{off1}/k_{on1}$)<br>0.21 ($k_{off1}/k_{on2}$)<br>$< 0.00835$ ($k_{off2}/k_{on1}$) | $1.31\times10^{4}$ ($k_{on1}$)<br>$8.29\times10^{4}$ ($k_{on2}$) | $1.39\times10^{-5}$ ($k_{off1}$)<br>$< 1.00\times10^{-7}$ ($k_{off2}$) | 1.06 ($k_{off1}/k_{on1}$)<br>0.168 ($k_{off1}/k_{on2}$)<br>$< 0.00762$ ($k_{off2}/k_{on1}$) | $1.78\times10^{4}$ ($k_{on1}$)<br>$3.01\times10^{4}$ ($k_{on2}$) | $5.39\times10^{-7}$ ($k_{off1}$)<br>$< 1.00\times10^{-7}$ ($k_{off2}$) | $\leqslant 0.0302$ ($k_{off1}/k_{on1}$) | $1.50\times10^{4}$ ($k_{on1}$)<br>$3.92\times10^{4}$ ($k_{on2}$) | $1.28\times10^{-5}$ ($k_{off1}$)<br>$< 1.00\times10^{-7}$ ($k_{off2}$) | 0.857 ($k_{off1}/k_{on1}$)<br>0.327 ($k_{off1}/k_{on2}$)<br>$< 0.00668$ ($k_{off2}/k_{on1}$) |
| **S-R/BA.1** | $1.48\times10^{4}$ ($k_{on1}$)<br>$1.43\times10^{5}$ ($k_{on2}$) | $7.78\times10^{-7}$ ($k_{off1}$)<br>$< 1.00\times10^{-7}$ ($k_{off2}$) | $\leqslant 0.0527$ ($k_{off1}/k_{on1}$) | $1.63\times10^{4}$ ($k_{on1}$)<br>$1.44\times10^{5}$ ($k_{on2}$) | $8.94\times10^{-5}$ ($k_{off1}$)<br>$< 1.00\times10^{-7}$ ($k_{off2}$) | 5.49 ($k_{off1}/k_{on1}$)<br>0.622 ($k_{off1}/k_{on2}$)<br>$< 0.00614$ ($k_{off2}/k_{on1}$) | $2.02\times10^{4}$ ($k_{on1}$)<br>$1.39\times10^{5}$ ($k_{on2}$) | $4.36\times10^{-7}$ ($k_{off1}$)<br>$< 1.00\times10^{-7}$ ($k_{off2}$) | $\leqslant 0.0216$ ($k_{off1}/k_{on1}$) | $2.18\times10^{4}$ ($k_{on1}$)<br>$1.36\times10^{5}$ ($k_{on2}$) | $2.28\times10^{-7}$ ($k_{off1}$)<br>$< 1.00\times10^{-7}$ ($k_{off2}$) | $\leqslant 0.0104$ ($k_{off1}/k_{on1}$) |
| **S-R/BA.2** | $1.21\times10^{4}$ ($k_{on1}$)<br>$1.02\times10^{5}$ ($k_{on2}$) | $< 1.00\times10^{-7}$ ($k_{off1}$)<br>$< 1.00\times10^{-7}$ ($k_{off2}$) | $< 0.00829$ ($k_{off1}/k_{on1}$) | $2.61\times10^{4}$ ($k_{on1}$)<br>$1.49\times10^{5}$ ($k_{on2}$) | $5.01\times10^{-5}$ ($k_{off1}$)<br>$< 1.00\times10^{-7}$ ($k_{off2}$) | 1.92 ($k_{off1}/k_{on1}$)<br>0.337 ($k_{off1}/k_{on2}$)<br>$< 0.00384$ ($k_{off2}/k_{on1}$) | $3.49\times10^{4}$ ($k_{on1}$)<br>$1.41\times10^{5}$ ($k_{on2}$) | $6.19\times10^{-7}$ ($k_{off1}$)<br>$< 1.00\times10^{-7}$ ($k_{off2}$) | $\leqslant 0.0178$ ($k_{off1}/k_{on1}$) | $3.04\times10^{4}$ ($k_{on1}$)<br>$1.39\times10^{5}$ ($k_{on2}$) | $< 1.00\times10^{-7}$ ($k_{off1}$)<br>$< 1.00\times10^{-7}$ ($k_{off2}$) | $< 0.00329$ ($k_{off1}/k_{on1}$) |
| **S-R/BA.4/5** | $1.58\times10^{4}$ ($k_{on1}$)<br>$7.49\times10^{4}$ ($k_{on2}$) | $9.87\times10^{-7}$ ($k_{off1}$)<br>$< 1.00\times10^{-7}$ ($k_{off2}$) | $\leqslant 0.0623$ ($k_{off1}/k_{on1}$) | $2.04\times10^{4}$ ($k_{on1}$)<br>$1.01\times10^{5}$ ($k_{on2}$) | $1.41\times10^{-7}$ ($k_{off1}$)<br>$< 1.00\times10^{-7}$ ($k_{off2}$) | $\leqslant 0.00691$ ($k_{off1}/k_{on1}$) | $2.99\times10^{4}$ ($k_{on1}$)<br>$8.61\times10^{4}$ ($k_{on2}$) | $1.57\times10^{-6}$ ($k_{off1}$)<br>$1.28\times10^{-7}$ ($k_{off2}$) | $\leqslant 0.0525$ ($k_{off1}/k_{on1}$) | $2.49\times10^{4}$ ($k_{on1}$)<br>$7.63\times10^{4}$ ($k_{on2}$) | $1.42\times10^{-6}$ ($k_{off1}$)<br>$1.91\times10^{-7}$ ($k_{off2}$) | $\leqslant 0.0569$ ($k_{off1}/k_{on1}$) |
| **S-R/BA.2.75.2** | $2.34\times10^{4}$ ($k_{on1}$)<br>$5.14\times10^{4}$ ($k_{on2}$) | $2.16\times10^{-5}$ ($k_{off1}$)<br>$1.82\times10^{-5}$ ($k_{off2}$) | 0.926 ($k_{off1}/k_{on1}$)<br>0.421 ($k_{off1}/k_{on2}$)<br>0.778 ($k_{off2}/k_{on1}$)<br>0.354 ($k_{off2}/k_{on2}$) | $1.88\times10^{4}$ ($k_{on1}$)<br>$4.95\times10^{4}$ ($k_{on2}$) | $4.13\times10^{-7}$ ($k_{off1}$)<br>$< 1.00\times10^{-7}$ ($k_{off2}$) | $\leqslant 0.022$ ($k_{off1}/k_{on1}$) | $2.06\times10^{4}$ ($k_{on1}$)<br>$6.24\times10^{4}$ ($k_{on2}$) | $5.10\times10^{-5}$ ($k_{off1}$)<br>$1.21\times10^{-6}$ ($k_{off2}$) | 2.48 ($k_{off1}/k_{on1}$)<br>0.818 ($k_{off1}/k_{on2}$)<br>$\leqslant 0.0589$ ($k_{off2}/k_{on1}$) | $1.99\times10^{4}$ ($k_{on1}$)<br>$5.92\times10^{4}$ ($k_{on2}$) | $1.86\times10^{-7}$ ($k_{off1}$)<br>$< 1.00\times10^{-7}$ ($k_{off2}$) | $\leqslant 0.00938$ ($k_{off1}/k_{on1}$) |
| **S-R/BQ.1.1** | $2.21\times10^{4}$ ($k_{on1}$)<br>$2.40\times10^{4}$ ($k_{on2}$) | $2.12\times10^{-6}$ ($k_{off1}$)<br>$< 1.00\times10^{-7}$ ($k_{off2}$) | $\leqslant 0.0962$ ($k_{off1}/k_{on1}$) | $1.95\times10^{4}$ ($k_{on1}$)<br>$5.55\times10^{4}$ ($k_{on2}$) | $6.52\times10^{-6}$ ($k_{off1}$)<br>$< 1.00\times10^{-7}$ ($k_{off2}$) | 0.335 ($k_{off1}/k_{on1}$)<br>0.118 ($k_{off1}/k_{on2}$)<br>$< 0.00514$ ($k_{off2}/k_{on1}$) | $2.13\times10^{4}$ ($k_{on1}$)<br>$2.88\times10^{4}$ ($k_{on2}$) | $2.43\times10^{-7}$ ($k_{off1}$)<br>$< 1.00\times10^{-7}$ ($k_{off2}$) | $\leqslant 0.0114$ ($k_{off1}/k_{on1}$) | $1.77\times10^{4}$ ($k_{on1}$)<br>$5.43\times10^{4}$ ($k_{on2}$) | $1.17\times10^{-6}$ ($k_{off1}$)<br>$< 1.00\times10^{-7}$ ($k_{off2}$) | $\leqslant 0.0661$ ($k_{off1}/k_{on1}$) |

### Table S3. Kinetic parameters of constructed R1-32-like mutant antibodies binding to RBDs (related to Fig. S11).

| | C092 | | | C092-germline HCDR1 | | | C092-R1-32 HCDR3 | | | C092-Y109A$_H$ | | | C092-germline HCDR1+Y109A$_H$ | | | C092-N26S$_L$+D99G$_L$ | | | C092-N26S$_L$+D70G$_L$+D99G$_L$ | | |
|---|---|---|---|---|---|---|---|---|---|---|---|---|---|---|---|---|---|---|---|---|---|
| | $k_{on}$ (M$^{-1}$ s$^{-1}$) | $k_{off}$ (s$^{-1}$) | $K_D$ (nM) | $k_{on}$ (M$^{-1}$ s$^{-1}$) | $k_{off}$ (s$^{-1}$) | $K_D$ (nM) | $k_{on}$ (M$^{-1}$ s$^{-1}$) | $k_{off}$ (s$^{-1}$) | $K_D$ (nM) | $k_{on}$ (M$^{-1}$ s$^{-1}$) | $k_{off}$ (s$^{-1}$) | $K_D$ (nM) | $k_{on}$ (M$^{-1}$ s$^{-1}$) | $k_{off}$ (s$^{-1}$) | $K_D$ (nM) | $k_{on}$ (M$^{-1}$ s$^{-1}$) | $k_{off}$ (s$^{-1}$) | $K_D$ (nM) | $k_{on}$ (M$^{-1}$ s$^{-1}$) | $k_{off}$ (s$^{-1}$) | $K_D$ (nM) |
| WT | $7.62\times10^{5}$ | $3.53\times10^{-5}$ | 0.046 | $5.70\times10^{5}$ | $2.23\times10^{-5}$ | 0.039 | $9.32\times10^{5}$ | $5.50\times10^{-5}$ | 0.059 | $5.93\times10^{5}$ | $6.60\times10^{-5}$ | 0.111 | $3.76\times10^{5}$ | $1.18\times10^{-4}$ | 0.314 | $7.26\times10^{5}$ | $5.56\times10^{-5}$ | 0.077 | $6.26\times10^{5}$ | $2.84\times10^{-5}$ | 0.045 |
| L452R | $3.04\times10^{5}$ | $2.19\times10^{-5}$ | 0.072 | $1.84\times10^{5}$ | $2.02\times10^{-4}$ | 1.100 | $1.98\times10^{5}$ | $2.14\times10^{-4}$ | 1.080 | $1.46\times10^{5}$ | $2.10\times10^{-4}$ | 1.440 | $9.88\times10^{4}$ | $2.68\times10^{-3}$ | 27.100 | $3.00\times10^{5}$ | $4.58\times10^{-4}$ | 1.530 | $2.11\times10^{5}$ | $8.72\times10^{-4}$ | 4.130 |
| F490S | $3.75\times10^{5}$ | $4.78\times10^{-5}$ | 0.128 | $5.76\times10^{5}$ | $9.57\times10^{-4}$ | 0.241 | $3.59\times10^{5}$ | $9.56\times10^{-5}$ | 0.266 | $2.13\times10^{5}$ | $8.40\times10^{-5}$ | 0.394 | $1.72\times10^{5}$ | $3.26\times10^{-4}$ | 1.890 | $3.29\times10^{5}$ | $8.53\times10^{-5}$ | 0.260 | $2.63\times10^{5}$ | $1.18\times10^{-4}$ | 0.450 |
| Kappa | $2.90\times10^{5}$ | $3.68\times10^{-5}$ | 0.127 | $1.54\times10^{5}$ | $1.82\times10^{-4}$ | 1.180 | $1.44\times10^{5}$ | $1.61\times10^{-4}$ | 1.120 | $1.11\times10^{5}$ | $1.81\times10^{-4}$ | 1.630 | $8.14\times10^{4}$ | $2.85\times10^{-3}$ | 34.900 | $2.32\times10^{5}$ | $4.35\times10^{-4}$ | 1.870 | $1.53\times10^{5}$ | $7.95\times10^{-4}$ | 5.220 |
| Delta | $3.64\times10^{5}$ | $6.52\times10^{-5}$ | 0.179 | $1.78\times10^{5}$ | $2.07\times10^{-4}$ | 1.160 | $1.87\times10^{5}$ | $1.91\times10^{-4}$ | 1.020 | $1.42\times10^{5}$ | $1.98\times10^{-4}$ | 1.400 | $9.93\times10^{4}$ | $2.85\times10^{-3}$ | 28.700 | $2.97\times10^{5}$ | $4.88\times10^{-4}$ | 1.640 | $2.13\times10^{5}$ | $9.30\times10^{-4}$ | 4.360 |
| Lambda | $2.22\times10^{5}$ | $9.67\times10^{-5}$ | 0.436 | $8.93\times10^{4}$ | $2.24\times10^{-4}$ | 2.510 | $1.00\times10^{5}$ | $6.88\times10^{-4}$ | 6.880 | $6.98\times10^{4}$ | $8.90\times10^{-4}$ | 12.800 | $6.22\times10^{4}$ | $3.47\times10^{-3}$ | 55.800 | $1.51\times10^{5}$ | $1.69\times10^{-3}$ | 11.200 | $1.05\times10^{5}$ | $3.36\times10^{-3}$ | 31.900 |
| BA.4/5 | $3.87\times10^{5}$ | $5.12\times10^{-5}$ | 0.132 | $2.15\times10^{5}$ | $3.06\times10^{-4}$ | 1.430 | $1.75\times10^{5}$ | $1.55\times10^{-4}$ | 0.881 | $1.18\times10^{5}$ | $2.24\times10^{-4}$ | 1.890 | $1.41\times10^{5}$ | $3.39\times10^{-3}$ | 24.100 | $2.46\times10^{5}$ | $5.90\times10^{-4}$ | 2.400 | $2.49\times10^{5}$ | $9.76\times10^{-4}$ | 3.920 |

| | C807 | | | C807-germline HCDR1 | | | C807-R1-32 HCDR3 | | | C807-Y109A$_H$ | | | C807-germline HCDR1+Y109A$_H$ | | |
|---|---|---|---|---|---|---|---|---|---|---|---|---|---|---|---|
| | $k_{on}$ (M$^{-1}$ s$^{-1}$) | $k_{off}$ (s$^{-1}$) | $K_D$ (nM) | $k_{on}$ (M$^{-1}$ s$^{-1}$) | $k_{off}$ (s$^{-1}$) | $K_D$ (nM) | $k_{on}$ (M$^{-1}$ s$^{-1}$) | $k_{off}$ (s$^{-1}$) | $K_D$ (nM) | $k_{on}$ (M$^{-1}$ s$^{-1}$) | $k_{off}$ (s$^{-1}$) | $K_D$ (nM) | $k_{on}$ (M$^{-1}$ s$^{-1}$) | $k_{off}$ (s$^{-1}$) | $K_D$ (nM) |
| WT | $6.56\times10^{5}$ | $7.74\times10^{-5}$ | 0.118 | $6.09\times10^{5}$ | $3.88\times10^{-4}$ | 0.637 | $2.65\times10^{5}$ | $3.16\times10^{-4}$ | 1.190 | $4.89\times10^{5}$ | $3.53\times10^{-4}$ | 0.720 | $3.87\times10^{5}$ | $1.17\times10^{-3}$ | 3.020 |
| L452R | $4.75\times10^{5}$ | $5.14\times10^{-4}$ | 1.080 | $6.52\times10^{5}$ | $2.68\times10^{-3}$ | 4.120 | $1.33\times10^{5}$ | $3.87\times10^{-3}$ | 29.000 | $3.35\times10^{5}$ | $7.33\times10^{-3}$ | 21.900 | $3.80\times10^{5}$ | $9.18\times10^{-3}$ | 24.200 |
| F490S | $3.43\times10^{5}$ | $1.50\times10^{-4}$ | 0.437 | $4.20\times10^{5}$ | $5.82\times10^{-4}$ | 1.390 | $6.60\times10^{4}$ | $2.23\times10^{-3}$ | 33.700 | $2.07\times10^{5}$ | $1.16\times10^{-3}$ | 5.600 | $2.44\times10^{5}$ | $1.63\times10^{-3}$ | 6.670 |
| Kappa | $4.18\times10^{5}$ | $3.40\times10^{-4}$ | 0.813 | $6.84\times10^{5}$ | $1.37\times10^{-3}$ | 2.000 | $1.12\times10^{5}$ | $1.90\times10^{-3}$ | 17.000 | $2.58\times10^{5}$ | $4.75\times10^{-3}$ | 18.400 | $4.29\times10^{5}$ | $4.46\times10^{-3}$ | 10.400 |
| Delta | $5.14\times10^{5}$ | $4.65\times10^{-4}$ | 0.904 | $6.46\times10^{5}$ | $2.71\times10^{-3}$ | 4.200 | $1.31\times10^{5}$ | $3.80\times10^{-3}$ | 29.000 | $3.22\times10^{5}$ | $7.74\times10^{-3}$ | 24.100 | $3.61\times10^{5}$ | $1.01\times10^{-2}$ | 27.800 |
| Lambda | $2.56\times10^{5}$ | $1.12\times10^{-3}$ | 4.360 | $2.26\times10^{5}$ | $4.64\times10^{-3}$ | 20.500 | $5.11\times10^{4}$ | $1.87\times10^{-2}$ | 366.000 | $1.17\times10^{5}$ | $1.17\times10^{-2}$ | 99.700 | $1.49\times10^{5}$ | $1.50\times10^{-2}$ | 100.000 |
| BA.4/5 | $4.47\times10^{5}$ | $3.31\times10^{-4}$ | 0.741 | $8.10\times10^{5}$ | $9.54\times10^{-4}$ | 1.180 | $1.78\times10^{5}$ | $1.87\times10^{-3}$ | 10.500 | $2.91\times10^{5}$ | $3.83\times10^{-3}$ | 13.200 | $5.62\times10^{5}$ | $2.82\times10^{-3}$ | 5.020 |

| | BD56-104 | | | BD56-104-germline HCDR1 | | | BD56-104-D59N$_H$ | | | BD56-104-D59N$_H$+E62Q$_H$ | | | BD56-104-R1-32 HCDR3 | | |
|---|---|---|---|---|---|---|---|---|---|---|---|---|---|---|---|
| | $k_{on}$ (M$^{-1}$ s$^{-1}$) | $k_{off}$ (s$^{-1}$) | $K_D$ (nM) | $k_{on}$ (M$^{-1}$ s$^{-1}$) | $k_{off}$ (s$^{-1}$) | $K_D$ (nM) | $k_{on}$ (M$^{-1}$ s$^{-1}$) | $k_{off}$ (s$^{-1}$) | $K_D$ (nM) | $k_{on}$ (M$^{-1}$ s$^{-1}$) | $k_{off}$ (s$^{-1}$) | $K_D$ (nM) | $k_{on}$ (M$^{-1}$ s$^{-1}$) | $k_{off}$ (s$^{-1}$) | $K_D$ (nM) |
| WT | $8.62\times10^{5}$ | $3.29\times10^{-5}$ | 0.038 | $8.95\times10^{5}$ | $7.36\times10^{-5}$ | 0.082 | $9.17\times10^{5}$ | $4.39\times10^{-5}$ | 0.048 | $7.00\times10^{5}$ | $1.80\times10^{-5}$ | 0.026 | $1.13\times10^{6}$ | $2.55\times10^{-5}$ | 0.023 |
| L452R | $5.53\times10^{5}$ | $2.66\times10^{-5}$ | 0.048 | $5.03\times10^{5}$ | $2.46\times10^{-4}$ | 0.489 | $1.97\times10^{5}$ | $2.80\times10^{-4}$ | 1.420 | $1.23\times10^{5}$ | $5.27\times10^{-4}$ | 4.270 | $6.34\times10^{5}$ | $8.21\times10^{-5}$ | 0.130 |
| F490S | $5.90\times10^{5}$ | $3.49\times10^{-4}$ | 0.592 | $5.76\times10^{5}$ | $9.57\times10^{-4}$ | 1.660 | $4.61\times10^{5}$ | $2.13\times10^{-4}$ | 0.462 | $3.33\times10^{5}$ | $3.12\times10^{-4}$ | 0.936 | $6.86\times10^{4}$ | $1.86\times10^{-4}$ | 0.271 |
| Kappa | $5.31\times10^{5}$ | $4.81\times10^{-5}$ | 0.091 | $4.88\times10^{5}$ | $2.56\times10^{-4}$ | 0.524 | $1.68\times10^{5}$ | $2.06\times10^{-4}$ | 1.220 | $1.01\times10^{5}$ | $4.04\times10^{-4}$ | 4.000 | $5.71\times10^{5}$ | $5.69\times10^{-5}$ | 0.100 |
| Delta | $5.10\times10^{5}$ | $4.16\times10^{-5}$ | 0.082 | $4.93\times10^{5}$ | $2.57\times10^{-4}$ | 0.521 | $1.90\times10^{5}$ | $2.84\times10^{-4}$ | 1.490 | $1.20\times10^{5}$ | $5.56\times10^{-4}$ | 4.640 | $5.80\times10^{5}$ | $2.90\times10^{-5}$ | 0.050 |
| Lambda | $2.77\times10^{5}$ | $1.87\times10^{-3}$ | 6.760 | $2.30\times10^{5}$ | $6.04\times10^{-3}$ | 26.300 | $1.59\times10^{5}$ | $1.11\times10^{-3}$ | 17.100 | $8.78\times10^{4}$ | $1.54\times10^{-3}$ | 17.600 | $2.89\times10^{5}$ | $7.89\times10^{-4}$ | 2.730 |
| BA.4/5 | $5.63\times10^{5}$ | $1.62\times10^{-4}$ | 0.029 | $5.18\times10^{5}$ | $4.24\times10^{-4}$ | 0.819 | $1.86\times10^{5}$ | $3.32\times10^{-4}$ | 1.790 | $1.18\times10^{5}$ | $5.47\times10^{-4}$ | 4.650 | $6.56\times10^{5}$ | $5.58\times10^{-5}$ | 0.085 |

| | BD56-597 | | | BD56-597-germline HCDR1 | | | BD56-597-D59N$_H$ | | | BD56-597-R1-32 HCDR3 | | | BD56-597-Y109A$_H$ | | | BD56-597-germline HCDR1-D59N$_H$+Y109A$_H$ | | | BD56-597-D26S$_L$ | | |
|---|---|---|---|---|---|---|---|---|---|---|---|---|---|---|---|---|---|---|---|---|---|
| | $k_{on}$ (M$^{-1}$ s$^{-1}$) | $k_{off}$ (s$^{-1}$) | $K_D$ (nM) | $k_{on}$ (M$^{-1}$ s$^{-1}$) | $k_{off}$ (s$^{-1}$) | $K_D$ (nM) | $k_{on}$ (M$^{-1}$ s$^{-1}$) | $k_{off}$ (s$^{-1}$) | $K_D$ (nM) | $k_{on}$ (M$^{-1}$ s$^{-1}$) | $k_{off}$ (s$^{-1}$) | $K_D$ (nM) | $k_{on}$ (M$^{-1}$ s$^{-1}$) | $k_{off}$ (s$^{-1}$) | $K_D$ (nM) | $k_{on}$ (M$^{-1}$ s$^{-1}$) | $k_{off}$ (s$^{-1}$) | $K_D$ (nM) | $k_{on}$ (M$^{-1}$ s$^{-1}$) | $k_{off}$ (s$^{-1}$) | $K_D$ (nM) |
| WT | $9.29\times10^{5}$ | $4.46\times10^{-5}$ | 0.048 | $9.35\times10^{5}$ | $2.71\times10^{-5}$ | 0.029 | $7.29\times10^{5}$ | $2.36\times10^{-5}$ | 0.032 | $9.11\times10^{5}$ | $5.74\times10^{-5}$ | 0.063 | $8.21\times10^{5}$ | $2.69\times10^{-5}$ | 0.033 | $9.50\times10^{5}$ | $1.55\times10^{-5}$ | 0.016 | $8.10\times10^{5}$ | $2.02\times10^{-5}$ | 0.025 |
| L452R | $6.66\times10^{5}$ | $5.29\times10^{-5}$ | 0.079 | $7.92\times10^{5}$ | $7.14\times10^{-5}$ | 0.090 | $4.09\times10^{5}$ | $7.11\times10^{-5}$ | 0.174 | $4.91\times10^{5}$ | $1.17\times10^{-4}$ | 0.239 | $7.09\times10^{5}$ | $3.34\times10^{-5}$ | 0.047 | $6.24\times10^{5}$ | $4.39\times10^{-4}$ | 0.703 | $5.45\times10^{5}$ | $4.82\times10^{-5}$ | 0.089 |
| F490S | $5.71\times10^{5}$ | $6.02\times10^{-5}$ | 0.105 | $7.00\times10^{5}$ | $2.40\times10^{-4}$ | 0.343 | $3.99\times10^{5}$ | $2.39\times10^{-5}$ | 0.060 | $4.84\times10^{5}$ | $3.01\times10^{-4}$ | 0.621 | $5.65\times10^{5}$ | $1.38\times10^{-4}$ | 0.244 | $6.19\times10^{5}$ | $1.48\times10^{-4}$ | 0.239 | $4.59\times10^{5}$ | $2.05\times10^{-5}$ | 0.045 |
| Kappa | $6.14\times10^{5}$ | $2.27\times10^{-5}$ | 0.037 | $7.34\times10^{5}$ | $3.05\times10^{-5}$ | 0.042 | $3.58\times10^{5}$ | $3.60\times10^{-5}$ | 0.101 | $4.60\times10^{5}$ | $1.34\times10^{-4}$ | 0.292 | $6.75\times10^{5}$ | $2.56\times10^{-5}$ | 0.038 | $5.57\times10^{5}$ | $4.19\times10^{-4}$ | 0.753 | $4.87\times10^{5}$ | $2.42\times10^{-5}$ | 0.050 |
| Delta | $6.22\times10^{5}$ | $2.17\times10^{-5}$ | 0.035 | $7.77\times10^{5}$ | $6.02\times10^{-5}$ | 0.078 | $3.87\times10^{5}$ | $5.34\times10^{-5}$ | 0.138 | $4.82\times10^{5}$ | $1.21\times10^{-4}$ | 0.251 | $6.96\times10^{5}$ | $2.94\times10^{-5}$ | 0.042 | $6.04\times10^{5}$ | $4.51\times10^{-4}$ | 0.746 | $5.21\times10^{5}$ | $2.99\times10^{-5}$ | 0.057 |
| Lambda | $2.90\times10^{5}$ | $1.62\times10^{-4}$ | 0.560 | $2.69\times10^{5}$ | $2.82\times10^{-3}$ | 10.500 | $1.82\times10^{5}$ | $3.02\times10^{-4}$ | 1.660 | $1.29\times10^{5}$ | $4.86\times10^{-3}$ | 37.700 | $3.16\times10^{5}$ | $2.82\times10^{-3}$ | 8.900 | $3.07\times10^{5}$ | $3.12\times10^{-3}$ | 10.100 | $2.12\times10^{5}$ | $6.67\times10^{-4}$ | 3.150 |
| BA.4/5 | $6.61\times10^{5}$ | $3.36\times10^{-5}$ | 0.051 | $8.21\times10^{5}$ | $5.73\times10^{-5}$ | 0.070 | $3.54\times10^{5}$ | $4.94\times10^{-5}$ | 0.139 | $4.92\times10^{5}$ | $2.03\times10^{-4}$ | 0.412 | $5.99\times10^{5}$ | $2.24\times10^{-5}$ | 0.037 | $6.21\times10^{5}$ | $3.89\times10^{-4}$ | 0.627 | $4.73\times10^{5}$ | $1.65\times10^{-5}$ | 0.035 |

**Table S4. Kinetic parameters of constructed R1-32-AAM antibody binding to SARS-CoV-2 RBDs (related to Fig. 4f).**

| | R1-32 | | | R1-32-AAM | | |
|---|---|---|---|---|---|---|
| | $k_{on}$ ($M^{-1}$ $s^{-1}$) | $k_{off}$ ($s^{-1}$) | $K_D$ (nM) | $k_{on}$ ($M^{-1}$ $s^{-1}$) | $k_{off}$ ($s^{-1}$) | $K_D$ (nM) |
| WT | $6.46\times10^{5}$ | $6.01\times10^{-4}$ | 0.931 | $6.24\times10^{5}$ | $7.35\times10^{-5}$ | 0.118 |
| L452R | $1.66\times10^{5}$ | $9.01\times10^{-3}$ | 54.210 | $4.59\times10^{5}$ | $1.07\times10^{-4}$ | 0.232 |
| Kappa | $1.29\times10^{5}$ | $1.19\times10^{-2}$ | 92.130 | $4.54\times10^{5}$ | $1.61\times10^{-4}$ | 0.353 |
| Delta | $1.58\times10^{5}$ | $1.15\times10^{-2}$ | 72.600 | $4.64\times10^{5}$ | $1.65\times10^{-4}$ | 0.355 |
| Lambda | $6.67\times10^{4}$ | $6.36\times10^{-2}$ | 953.000 | $8.65\times10^{4}$ | $1.03\times10^{-3}$ | 11.950 |
| BA.4/5 | $1.84\times10^{5}$ | $1.02\times10^{-2}$ | 55.520 | $4.80\times10^{5}$ | $7.98\times10^{-5}$ | 0.166 |
| BF.7 | $1.14\times10^{5}$ | $1.48\times10^{-2}$ | 129.300 | $1.53\times10^{5}$ | $7.63\times10^{-5}$ | 0.503 |
| BQ.1.1 | $1.02\times10^{5}$ | $2.15\times10^{-2}$ | 211.700 | $1.14\times10^{5}$ | $9.62\times10^{-5}$ | 0.843 |
| F490S | $2.82\times10^{5}$ | $3.34\times10^{-3}$ | 11.830 | $2.81\times10^{5}$ | $1.80\times10^{-4}$ | 0.643 |
| F490W | $2.30\times10^{5}$ | $2.78\times10^{-3}$ | 12.090 | $2.73\times10^{5}$ | $1.21\times10^{-5}$ | 0.044 |
| XBB | $2.29\times10^{5}$ | $1.13\times10^{-2}$ | 49.220 | $1.15\times10^{5}$ | $2.50\times10^{-4}$ | 2.178 |

**Table S5. Kinetic parameters of mutation-tolerant antibodies cross-binding to SARSr-CoV RBDs (related to Fig. S12).**

| | R1-32 | | | ZL525 | | |
|---|---|---|---|---|---|---|
| | $k_{on}$ ($M^{-1}$ $s^{-1}$) | $k_{off}$ ($s^{-1}$) | $K_D$ (nM) | $k_{on}$ ($M^{-1}$ $s^{-1}$) | $k_{off}$ ($s^{-1}$) | $K_D$ (nM) |
| WT | $6.46\times10^{5}$ | $6.01\times10^{-4}$ | 0.931 | $9.46\times10^{5}$ | $2.80\times10^{-4}$ | 0.295 |
| Pangolin-GD-2019 | $3.86\times10^{5}$ | $3.00\times10^{-4}$ | 0.777 | $2.61\times10^{5}$ | $1.04\times10^{-5}$ | 0.040 |
| Bat-RaTG13 | $2.07\times10^{5}$ | $7.92\times10^{-3}$ | 38.200 | $1.98\times10^{5}$ | $3.92\times10^{-6}$ | 0.020 |
| Pangolin-GX-2017 | $1.80\times10^{5}$ | $8.59\times10^{-4}$ | 4.760 | $3.55\times10^{5}$ | $3.70\times10^{-5}$ | 0.104 |
| Laos-20-52 | $7.48\times10^{5}$ | $7.21\times10^{-4}$ | 0.963 | $5.97\times10^{5}$ | $2.16\times10^{-4}$ | 0.362 |
| Laos-20-236 | $3.28\times10^{5}$ | $5.90\times10^{-4}$ | 1.800 | $3.31\times10^{5}$ | $2.68\times10^{-4}$ | 0.810 |
| Bat-RsSHC014 | $1.80\times10^{4}$ | $5.35\times10^{-3}$ | 297.000 | $9.04\times10^{4}$ | $4.04\times10^{-5}$ | 0.447 |
| Bat-WIV1 | — | — | no binding | $7.60\times10^{4}$ | $5.73\times10^{-4}$ | 7.540 |
| SARS-CoV-1 | — | — | no binding | $1.56\times10^{5}$ | $9.89\times10^{-3}$ | 63.100 |
| BtKY72 | — | — | no binding | $2.03\times10^{4}$ | $3.66\times10^{-4}$ | 18.000 |
| GX2013 | — | — | no binding | $2.61\times10^{5}$ | $1.02\times10^{-3}$ | 3.890 |
| HeB2013 | — | — | no binding | $1.07\times10^{5}$ | $1.51\times10^{-3}$ | 14.100 |
| RmYN02 | — | — | no binding | $2.96\times10^{5}$ | $7.79\times10^{-4}$ | 2.630 |

| | C092 | | | C807 | | |
|---|---|---|---|---|---|---|
| | $k_{on}$ ($M^{-1}$ $s^{-1}$) | $k_{off}$ ($s^{-1}$) | $K_D$ (nM) | $k_{on}$ ($M^{-1}$ $s^{-1}$) | $k_{off}$ ($s^{-1}$) | $K_D$ (nM) |
| WT | $7.62\times10^{5}$ | $3.53\times10^{-5}$ | 0.046 | $6.56\times10^{5}$ | $7.74\times10^{-5}$ | 0.118 |
| Pangolin-GD-2019 | $4.10\times10^{5}$ | $4.55\times10^{-5}$ | 0.111 | $2.84\times10^{5}$ | $1.74\times10^{-4}$ | 0.613 |
| Bat-RaTG13 | $1.63\times10^{5}$ | $2.88\times10^{-5}$ | 0.177 | $1.45\times10^{5}$ | $1.07\times10^{-4}$ | 0.738 |
| Pangolin-GX-2017 | $1.94\times10^{5}$ | $5.08\times10^{-5}$ | 0.262 | $2.02\times10^{5}$ | $9.47\times10^{-5}$ | 0.469 |
| Laos-20-52 | $5.25\times10^{5}$ | $3.10\times10^{-5}$ | 0.059 | $3.83\times10^{5}$ | $3.73\times10^{-4}$ | 0.972 |
| Laos-20-236 | $6.55\times10^{5}$ | $2.18\times10^{-5}$ | 0.034 | $4.87\times10^{5}$ | $3.35\times10^{-4}$ | 0.687 |
| Bat-RsSHC014 | $8.89\times10^{4}$ | $5.53\times10^{-5}$ | 0.622 | $5.97\times10^{4}$ | $2.09\times10^{-4}$ | 3.500 |
| Bat-WIV1 | $4.76\times10^{4}$ | $1.01\times10^{-3}$ | 21.300 | $4.86\times10^{4}$ | $3.65\times10^{-4}$ | 7.500 |
| SARS-CoV-1 | — | — | no binding | $1.34\times10^{5}$ | $4.57\times10^{-2}$ | 341.000 |
| BtKY72 | $1.41\times10^{5}$ | $4.13\times10^{-3}$ | 29.200 | $1.69\times10^{5}$ | $9.19\times10^{-3}$ | 54.500 |
| GX2013 | $1.88\times10^{5}$ | $4.94\times10^{-3}$ | 26.300 | $3.82\times10^{5}$ | $1.19\times10^{-2}$ | 31.100 |
| HeB2013 | $1.72\times10^{5}$ | $5.98\times10^{-3}$ | 34.900 | $1.91\times10^{5}$ | $1.98\times10^{-2}$ | 103.000 |
| RmYN02 | $1.04\times10^{5}$ | $3.28\times10^{-3}$ | 31.500 | $2.20\times10^{5}$ | $4.46\times10^{-3}$ | 20.300 |

| | BD56-104 | | | BD56-597 | | |
|---|---|---|---|---|---|---|
| | $k_{on}$ ($M^{-1}$ $s^{-1}$) | $k_{off}$ ($s^{-1}$) | $K_D$ (nM) | $k_{on}$ ($M^{-1}$ $s^{-1}$) | $k_{off}$ ($s^{-1}$) | $K_D$ (nM) |
| WT | $8.62\times10^{5}$ | $3.29\times10^{-5}$ | 0.038 | $9.29\times10^{5}$ | $4.46\times10^{-5}$ | 0.048 |
| Pangolin-GD-2019 | $4.26\times10^{5}$ | $7.53\times10^{-5}$ | 0.177 | $4.09\times10^{5}$ | $6.54\times10^{-5}$ | 0.160 |
| Bat-RaTG13 | $1.95\times10^{5}$ | $4.42\times10^{-5}$ | 0.227 | $2.25\times10^{5}$ | $6.39\times10^{-5}$ | 0.284 |
| Pangolin-GX-2017 | $3.77\times10^{5}$ | $4.40\times10^{-5}$ | 0.117 | $4.22\times10^{5}$ | $3.61\times10^{-5}$ | 0.086 |
| Laos-20-52 | $6.66\times10^{5}$ | $9.55\times10^{-5}$ | 0.144 | $6.12\times10^{5}$ | $6.39\times10^{-5}$ | 0.104 |
| Laos-20-236 | $8.64\times10^{5}$ | $7.10\times10^{-5}$ | 0.082 | $7.79\times10^{5}$ | $2.58\times10^{-5}$ | 0.033 |
| Bat-RsSHC014 | $2.03\times10^{4}$ | $9.75\times10^{-4}$ | 48.000 | $9.41\times10^{4}$ | $1.86\times10^{-5}$ | 0.198 |
| Bat-WIV1 | — | — | no binding | $5.26\times10^{4}$ | $3.79\times10^{-4}$ | 7.190 |
| SARS-CoV-1 | — | — | no binding | $6.03\times10^{5}$ | $3.55\times10^{-1}$ | 588.000 |
| BtKY72 | — | — | weak binding | $8.70\times10^{4}$ | $1.94\times10^{-2}$ | 223.000 |
| GX2013 | — | — | weak binding | $3.15\times10^{5}$ | $5.01\times10^{-2}$ | 159.000 |
| HeB2013 | — | — | weak binding | — | — | weak binding |
| RmYN02 | — | — | weak binding | $2.21\times10^{5}$ | $6.35\times10^{-2}$ | 287.000 |

### Table S6. Kinetic parameters of antibody binding to constructed RBD mutants (related to Fig. S13a).

| | C092 | | | C807 | | | BD56-104 | | | BD56-597 | | |
|---|---|---|---|---|---|---|---|---|---|---|---|---|
| | $k_{on}$ ($M^{-1}$ $s^{-1}$) | $k_{off}$ ($s^{-1}$) | $K_D$ (nM) | $k_{on}$ ($M^{-1}$ $s^{-1}$) | $k_{off}$ ($s^{-1}$) | $K_D$ (nM) | $k_{on}$ ($M^{-1}$ $s^{-1}$) | $k_{off}$ ($s^{-1}$) | $K_D$ (nM) | $k_{on}$ ($M^{-1}$ $s^{-1}$) | $k_{off}$ ($s^{-1}$) | $K_D$ (nM) |
| WT | $7.62\times10^{5}$ | $3.53\times10^{-5}$ | 0.046 | $6.56\times10^{5}$ | $7.74\times10^{-5}$ | 0.118 | $8.62\times10^{5}$ | $3.29\times10^{-5}$ | 0.038 | $9.29\times10^{5}$ | $4.46\times10^{-5}$ | 0.048 |
| Y351F | $6.58\times10^{5}$ | $1.53\times10^{-4}$ | 0.233 | $6.26\times10^{5}$ | $1.80\times10^{-4}$ | 0.287 | $6.74\times10^{5}$ | $8.06\times10^{-5}$ | 0.120 | $7.75\times10^{5}$ | $1.81\times10^{-4}$ | 0.234 |
| I468V | $6.52\times10^{5}$ | $1.13\times10^{-4}$ | 0.173 | $5.39\times10^{5}$ | $3.82\times10^{-4}$ | 0.709 | $7.61\times10^{5}$ | $2.19\times10^{-4}$ | 0.288 | $7.18\times10^{5}$ | $5.65\times10^{-5}$ | 0.079 |
| I468T | $6.30\times10^{5}$ | $1.41\times10^{-4}$ | 0.223 | $5.22\times10^{5}$ | $3.57\times10^{-4}$ | 0.683 | $6.82\times10^{5}$ | $1.23\times10^{-4}$ | 0.180 | $7.50\times10^{5}$ | $1.30\times10^{-4}$ | 0.173 |
| T470N | $4.31\times10^{5}$ | $1.32\times10^{-4}$ | 0.305 | $4.12\times10^{5}$ | $2.20\times10^{-4}$ | 0.534 | $6.09\times10^{5}$ | $2.41\times10^{-4}$ | 0.396 | $5.86\times10^{5}$ | $1.15\times10^{-4}$ | 0.195 |
| K462R | $7.22\times10^{5}$ | $1.37\times10^{-4}$ | 0.189 | $5.97\times10^{5}$ | $1.31\times10^{-4}$ | 0.219 | $7.50\times10^{5}$ | $3.79\times10^{-5}$ | 0.051 | $7.31\times10^{5}$ | $1.01\times10^{-4}$ | 0.138 |
| E471V | $6.13\times10^{5}$ | $1.44\times10^{-4}$ | 0.235 | $4.82\times10^{5}$ | $2.59\times10^{-4}$ | 0.537 | $7.53\times10^{5}$ | $8.45\times10^{-4}$ | 0.112 | $6.96\times10^{5}$ | $1.45\times10^{-4}$ | 0.208 |
| Loop-R | $7.21\times10^{5}$ | $8.65\times10^{-5}$ | 0.120 | $5.20\times10^{5}$ | $1.81\times10^{-4}$ | 0.348 | $7.90\times10^{5}$ | $1.23\times10^{-4}$ | 0.156 | $7.21\times10^{5}$ | $1.64\times10^{-4}$ | 0.227 |
| I472L | $6.35\times10^{5}$ | $1.16\times10^{-4}$ | 0.183 | $5.48\times10^{5}$ | $1.78\times10^{-4}$ | 0.324 | $7.72\times10^{5}$ | $9.88\times10^{-5}$ | 0.128 | $7.23\times10^{5}$ | $9.55\times10^{-5}$ | 0.132 |
| I472V | $5.76\times10^{5}$ | $9.51\times10^{-5}$ | 0.165 | $4.99\times10^{5}$ | $1.88\times10^{-4}$ | 0.376 | $7.20\times10^{5}$ | $1.00\times10^{-4}$ | 0.139 | $6.69\times10^{5}$ | $9.83\times10^{-5}$ | 0.147 |
| I472T | $5.34\times10^{5}$ | $9.34\times10^{-5}$ | 0.175 | $4.53\times10^{5}$ | $1.51\times10^{-4}$ | 0.334 | $6.16\times10^{5}$ | $1.03\times10^{-4}$ | 0.167 | $5.78\times10^{5}$ | $1.05\times10^{-4}$ | 0.182 |

### Table S7. Kinetic parameters of antibody binding to constructed Lambda RBD mutants (related to Fig. S13b).

| | C092 | | | C807 | | | BD56-104 | | | BD56-597 | | |
|---|---|---|---|---|---|---|---|---|---|---|---|---|
| | $k_{on}$ ($M^{-1}$ $s^{-1}$) | $k_{off}$ ($s^{-1}$) | $K_D$ (nM) | $k_{on}$ ($M^{-1}$ $s^{-1}$) | $k_{off}$ ($s^{-1}$) | $K_D$ (nM) | $k_{on}$ ($M^{-1}$ $s^{-1}$) | $k_{off}$ ($s^{-1}$) | $K_D$ (nM) | $k_{on}$ ($M^{-1}$ $s^{-1}$) | $k_{off}$ ($s^{-1}$) | $K_D$ (nM) |
| WT | $7.62\times10^{5}$ | $3.53\times10^{-5}$ | 0.046 | $6.56\times10^{5}$ | $7.74\times10^{-5}$ | 0.118 | $8.62\times10^{5}$ | $3.29\times10^{-5}$ | 0.038 | $9.29\times10^{5}$ | $4.46\times10^{-5}$ | 0.048 |
| Lambda | $2.22\times10^{5}$ | $9.67\times10^{-5}$ | 0.436 | $2.56\times10^{5}$ | $1.12\times10^{-3}$ | 4.360 | $2.77\times10^{5}$ | $1.87\times10^{-3}$ | 6.760 | $2.90\times10^{5}$ | $1.62\times10^{-4}$ | 0.560 |
| Lambda-R346T | $1.33\times10^{5}$ | $2.71\times10^{-4}$ | 2.040 | $1.85\times10^{5}$ | $4.83\times10^{-3}$ | 26.200 | $7.29\times10^{4}$ | $7.90\times10^{-3}$ | 108.000 | $1.71\times10^{5}$ | $5.27\times10^{-4}$ | 3.080 |
| Lambda-R357T | $2.05\times10^{5}$ | $8.92\times10^{-4}$ | 4.340 | $1.52\times10^{5}$ | $1.13\times10^{-2}$ | 74.500 | $2.12\times10^{5}$ | $3.03\times10^{-2}$ | 143.000 | $1.84\times10^{5}$ | $6.22\times10^{-3}$ | 33.800 |
| Lambda-T470N | $1.18\times10^{5}$ | $2.80\times10^{-4}$ | 2.370 | $1.98\times10^{5}$ | $5.75\times10^{-3}$ | 29.100 | $8.44\times10^{4}$ | $1.98\times10^{-2}$ | 235.000 | $2.48\times10^{5}$ | $1.43\times10^{-3}$ | 5.770 |
| Lambda-R346T-T470N | $9.08\times10^{4}$ | $2.29\times10^{-3}$ | 25.200 | $1.76\times10^{5}$ | $3.17\times10^{-2}$ | 180.000 | — | — | no binding | $1.60\times10^{5}$ | $6.14\times10^{-3}$ | 38.300 |
| Lambda-R346T-R357T-T470N | — | — | weak binding | — | — | no binding | — | — | no binding | — | — | no binding |

### Table S8. Kinetic parameters of antibody binding to RBDs of newer variants and RBD glycan mutants (T356K) (related to Fig. S16)

**ZL525**

| | $k_{on}$ ($M^{-1}$ $s^{-1}$) | $k_{off}$ ($s^{-1}$) | $K_D$ (nM) |
|---|---|---|---|
| WT | $9.46\times10^{5}$ | $2.80\times10^{-4}$ | 0.295 |
| L452R | $7.17\times10^{5}$ | $1.84\times10^{-4}$ | 0.257 |
| Delta | $6.36\times10^{5}$ | $1.99\times10^{-4}$ | 0.314 |
| Kappa | $6.96\times10^{5}$ | $2.32\times10^{-4}$ | 0.334 |
| Lambda | $4.18\times10^{5}$ | $2.53\times10^{-4}$ | 0.605 |
| BA.4 | $7.45\times10^{5}$ | $6.62\times10^{-5}$ | 0.089 |
| BQ.1.1 | $6.49\times10^{5}$ | $7.79\times10^{-5}$ | 0.120 |
| XBB.1.5 | $4.61\times10^{5}$ | $3.26\times10^{-4}$ | 0.708 |
| EG.5.1 | $4.06\times10^{5}$ | $1.83\times10^{-5}$ | 0.045 |
| HK.3 | $4.15\times10^{5}$ | $3.08\times10^{-6}$ | 0.007 |
| JD.1.1 | $4.67\times10^{5}$ | $2.10\times10^{-5}$ | 0.045 |
| BA.2.86 | $2.74\times10^{5}$ | $3.61\times10^{-5}$ | 0.131 |
| JN.1 | $3.11\times10^{5}$ | $7.30\times10^{-5}$ | 0.234 |
| KP.2 | $1.29\times10^{5}$ | $1.43\times10^{-4}$ | 1.110 |
| KP.3 | $2.53\times10^{5}$ | $1.21\times10^{-4}$ | 0.475 |
| BA.2.86-T356K | $6.95\times10^{5}$ | $2.41\times10^{-5}$ | 0.035 |
| JN.1-T356K | $6.79\times10^{5}$ | $2.44\times10^{-5}$ | 0.036 |
| KP.2-T356K | $6.87\times10^{5}$ | $4.98\times10^{-5}$ | 0.073 |
| KP.3-T356K | $7.15\times10^{5}$ | $9.85\times10^{-5}$ | 0.138 |

**C092**

| | $k_{on}$ ($M^{-1}$ $s^{-1}$) | $k_{off}$ ($s^{-1}$) | $K_D$ (nM) |
|---|---|---|---|
| WT | $7.62\times10^{5}$ | $3.53\times10^{-5}$ | 0.046 |
| HK.3 | $3.54\times10^{5}$ | $5.66\times10^{-6}$ | 0.016 |
| JD.1.1 | $4.04\times10^{5}$ | $2.06\times10^{-5}$ | 0.051 |
| BA.2.86 | $3.14\times10^{4}$ | $4.95\times10^{-5}$ | 1.579 |
| JN.1 | $3.35\times10^{4}$ | $4.74\times10^{-5}$ | 1.414 |
| KP.2 | $3.03\times10^{4}$ | $4.98\times10^{-6}$ | 0.164 |
| KP.3 | $3.98\times10^{4}$ | $1.42\times10^{-5}$ | 0.356 |
| BA.2.86-T356K | $5.64\times10^{5}$ | $4.80\times10^{-5}$ | 0.085 |
| JN.1-T356K | $5.05\times10^{5}$ | $7.34\times10^{-5}$ | 0.145 |
| KP.2-T356K | $5.98\times10^{5}$ | $9.26\times10^{-5}$ | 0.155 |
| KP.3-T356K | $6.19\times10^{5}$ | $2.82\times10^{-5}$ | 0.046 |

**BD56-104**

| | $k_{on}$ ($M^{-1}$ $s^{-1}$) | $k_{off}$ ($s^{-1}$) | $K_D$ (nM) |
|---|---|---|---|
| WT | $8.62\times10^{5}$ | $3.29\times10^{-5}$ | 0.038 |
| HK.3 | $5.22\times10^{5}$ | $2.65\times10^{-3}$ | 5.073 |
| JD.1.1 | $5.79\times10^{5}$ | $2.98\times10^{-3}$ | 5.145 |
| BA.2.86 | $3.70\times10^{4}$ | $2.14\times10^{-3}$ | 57.750 |
| JN.1 | $2.95\times10^{4}$ | $1.45\times10^{-3}$ | 49.070 |
| KP.2 | $4.53\times10^{4}$ | $1.05\times10^{-2}$ | 230.800 |
| KP.3 | $3.40\times10^{4}$ | $1.95\times10^{-3}$ | 57.270 |
| BA.2.86-T356K | $6.50\times10^{5}$ | $4.76\times10^{-3}$ | 7.323 |
| JN.1-T356K | $6.45\times10^{5}$ | $3.69\times10^{-3}$ | 5.714 |
| KP.2-T356K | $3.64\times10^{5}$ | $4.08\times10^{-2}$ | 112.100 |
| KP.3-T356K | $6.62\times10^{5}$ | $5.02\times10^{-3}$ | 7.589 |

**C807**

| | $k_{on}$ ($M^{-1}$ $s^{-1}$) | $k_{off}$ ($s^{-1}$) | $K_D$ (nM) |
|---|---|---|---|
| WT | $6.56\times10^{5}$ | $7.74\times10^{-5}$ | 0.118 |
| HK.3 | $4.05\times10^{5}$ | $2.89\times10^{-4}$ | 0.715 |
| JD.1.1 | $4.49\times10^{5}$ | $2.33\times10^{-4}$ | 0.518 |
| BA.2.86 | $2.59\times10^{4}$ | $1.07\times10^{-3}$ | 41.160 |
| JN.1 | $2.34\times10^{4}$ | $6.63\times10^{-4}$ | 28.360 |
| KP.2 | $4.07\times10^{4}$ | $3.98\times10^{-3}$ | 97.710 |
| KP.3 | $2.62\times10^{4}$ | $1.30\times10^{-3}$ | 49.840 |
| BA.2.86-T356K | $5.10\times10^{5}$ | $2.77\times10^{-3}$ | 5.440 |
| JN.1-T356K | $4.96\times10^{5}$ | $2.01\times10^{-3}$ | 4.044 |
| KP.2-T356K | $3.92\times10^{5}$ | $1.23\times10^{-2}$ | 31.240 |
| KP.3-T356K | $5.33\times10^{5}$ | $3.61\times10^{-3}$ | 6.772 |

**BD56-597**

| | $k_{on}$ ($M^{-1}$ $s^{-1}$) | $k_{off}$ ($s^{-1}$) | $K_D$ (nM) |
|---|---|---|---|
| WT | $9.29\times10^{5}$ | $4.46\times10^{-5}$ | 0.048 |
| HK.3 | $4.58\times10^{5}$ | $3.03\times10^{-5}$ | 0.066 |
| JD.1.1 | $5.03\times10^{5}$ | $1.60\times10^{-5}$ | 0.032 |
| BA.2.86 | $3.23\times10^{4}$ | $7.32\times10^{-5}$ | 2.267 |
| JN.1 | $2.20\times10^{4}$ | $2.30\times10^{-5}$ | 1.043 |
| KP.2 | $3.76\times10^{4}$ | $6.31\times10^{-4}$ | 16.760 |
| KP.3 | $3.12\times10^{4}$ | $1.16\times10^{-4}$ | 3.725 |
| BA.2.86-T356K | $5.41\times10^{5}$ | $6.06\times10^{-5}$ | 0.112 |
| JN.1-T356K | $5.34\times10^{5}$ | $1.58\times10^{-5}$ | 0.030 |
| KP.2-T356K | $5.53\times10^{5}$ | $7.21\times10^{-4}$ | 1.302 |
| KP.3-T356K | $5.73\times10^{5}$ | $1.31\times10^{-4}$ | 0.228 |

**Table S9. Cryo-EM data collection, refinement, and validation statistics.**

| | C092 Fab:ACE2: BA.4/5:S-6P 2 RBD "up" (EMD-66891, PDB 9XHZ) | C092 Fab:ACE2: BA.4/5:S-6P 3 RBD "up" (EMD-66892, PDB 9XI0) | C092 Fab:ACE2: BA.4/5 S1 (EMD-66890, PDB 9XHY) | C092 Fab:ACE2: BA.4/5 S1 (EMD-66893, PDB 9XI1) | C807 Fab:ACE2: BA.4/5:S-6P 3 RBD "up" (EMD-66895, PDB 9XI3) | C807 Fab:ACE2: BA.4/5 S1 (EMD-66894, PDB 9XI2) |
|---|---|---|---|---|---|---|
| **Data collection and processing** | | | | | | |
| Magnification | 45000 | 45000 | 45000 | 16500 | 16500 | 16500 |
| Voltage (kV) | 200 | 200 | 200 | 300 | 300 | 300 |
| Electron exposure ($e^{-}$/Å$^2$) | 60 | 60 | 60 | 50 | 50 | 50 |
| Defocus range (μm) | 0.8-2.5 | 0.8-2.5 | 0.8-2.5 | 0.6-2.0 | 0.6-2.0 | 0.6-2.0 |
| Pixel size (Å) | 0.88 | 0.88 | 0.88 | 0.73 | 0.73 | 0.73 |
| Movies (no.) | 5,405 | 5,405 | 5,405 | 10,614 | 5,878 | 5,878 |
| Initial particle images (no.) | 1,865,286 | 1,865,286 | 1,865,286 | 1,691,029 | 923,614 | 923,614 |
| Symmetry imposed | *C1* | *C1* | *C1* | *C1* | *C1* | *C1* |
| Final particle images (no.) | 56,953 | 148,642 | 229,072 | 548,261 | 15,224 | 336,871 |
| Map resolution (Å) | 5.28 | 3.98 | 3.84 | 2.58 | 4.74 | 2.83 |
| FSC threshold | 0.143 | 0.143 | 0.143 | 0.143 | 0.143 | 0.143 |
| Map resolution range (Å) | 4.15- 17.52 | 3.60-14.02 | 3.10-7.07 | 2.47-7.75 | 3.65-25.76 | 2.71-9.14 |
| **Refinement** | | | | | | |
| Initial model used | PDB 7YE5 | PDB 7YEG | PDB 7YDI | PDB 7YDI | PDB 7YEG | PDB 7YDI |
| Model resolution (Å) | 7.17 | 4.18 | 4.01 | 2.68 | 7.12 | 2.95 |
| FSC threshold | 0.5 | 0.5 | 0.5 | 0.5 | 0.5 | 0.5 |
| Map sharpening *B* factor (Å$^2$) | | | | 76.2 | | 81.3 |
| Model composition | | | | | | |
| Non-hydrogen atoms | 41525 | 49646 | 9678 | 9678 | 49626 | 9671 |
| Protein residues | 5215 | 6246 | 1224 | 1224 | 6246 | 1224 |
| Ligands | 46 | 51 | 5 | 5 | 51 | 5 |
| *B* factors (Å$^2$) | | | | | | |
| Protein | 537.86 | 209.85 | 76.65 | 41.76 | 524.16 | 11.84 |
| Ligand | 366.93 | 137.58 | 81.51 | 54.63 | 308.11 | 21.53 |
| R.m.s. deviations | | | | | | |
| Bond lengths (Å) | 0.002 | 0.004 | 0.004 | 0.003 | 0.002 | 0.004 |
| Bond angles (°) | 0.531 | 0.687 | 0.697 | 0.630 | 0.560 | 0.6619 |
| **Validation** | | | | | | |
| MolProbity score | 1.57 | 1.74 | 1.45 | 1.53 | 1.57 | 1.54 |
| Clashscore | 5.87 | 7.27 | 4.22 | 5.17 | 5.99 | 5.76 |
| Poor rotamers (%) | 0.00 | 0.11 | 0.00 | 1.14 | 0.09 | 0.96 |
| Ramachandran plot | | | | | | |
| Favored (%) | 96.34 | 95.06 | 96.30 | 96.71 | 96.37 | 96.55 |
| Allowed (%) | 3.66 | 4.94 | 3.70 | 3.29 | 3.63 | 3.45 |
| Disallowed (%) | 0.00 | 0.00 | 0.00 | 0.00 | 0.00 | 0.00 |

| | BD56-104 Fab: ACE2:BA.4/5 S-6P 3 RBD "up" (EMD-66897, PDB 9XI5) | BD56-104 Fab: ACE2:BA.4/5 S1 (EMD-66896, PDB 9XI4) | BD56-597 Fab: ACE2:BA.4/5 S-6P 3 RBD "up" (EMD-66899, PDB 9XI7) | BD56-597 Fab: ACE2:BA.4/5 S1 (EMD-66898, PDB 9XI6) | C092 Fab: D1H4 Fab: GX2013 S1 (EMD-66900, PDB 9XI8) | ZL525 Fab: ZL58 Fab: LP.8.1 RBD (EMD-66901, PDB 9XI9) |
|---|---|---|---|---|---|---|
| **Data collection and processing** | | | | | | |
| Magnification | 16500 | 16500 | 16500 | 16500 | 45000 | 165000 |
| Voltage (kV) | 300 | 300 | 300 | 300 | 200 | 300 |
| Electron exposure ($e^{-}/Å^2$) | 50 | 50 | 50 | 50 | 60 | 50 |
| Defocus range (μm) | 0.6-2.0 | 0.6-2.0 | 0.6-2.0 | 0.6-2.0 | 0.8-2.5 | 0.6-2.0 |
| Pixel size (Å) | 0.73 | 0.73 | 0.73 | 0.73 | 0.88 | 0.73 |
| Movies (no.) | 10,426 | 10,426 | 10,849 | 10,849 | 1,897 | 6,117 |
| Initial particle images (no.) | 1,416,220 | 1,416,220 | 1,416,145 | 1,416,145 | 1,619,759 | 696,375 |
| Symmetry imposed | *C1* | *C1* | *C1* | *C1* | *C1* | *C1* |
| Final particle images (no.) | 113,516 | 344,337 | 65,205 | 401,936 | 385,893 | 237,455 |
| Map resolution (Å) | 3.06 | 2.83 | 3.22 | 2.60 | 3.45 | 2.93 |
| FSC threshold | 0.143 | 0.143 | 0.143 | 0.143 | 0.143 | 0.143 |
| Map resolution range (Å) | 2.60-14.49 | 2.65-8.62 | 2.71-15.92 | 2.49-8.51 | 2.93-10.45 | 2.55-8.37 |
| **Refinement** | | | | | | |
| Initial model used | PDB 7YEG | PDB 7YDI | PDB 7YEG | PDB 7YDI | 8ZY6 | 8WXL |
| Model resolution (Å) | 3.34 | 2.82 | 3.74 | 2.72 | 3.81 | 3.01 |
| FSC threshold | 0.5 | 0.5 | 0.5 | 0.5 | 0.5 | 0.5 |
| Map sharpening *B* factor ($Å^2$) | | 79 | | 72.9 | 160.9 | 81.3 |
| Model composition | | | | | | |
| Non-hydrogen atoms | 49581 | 9656 | 49617 | 9668 | 4622 | 8088 |
| Protein residues | 6246 | 1224 | 6246 | 1224 | 612 | 1061 |
| Ligands | 51 | 5 | 51 | 5 | 0 | 3 |
| *B* factors ($Å^2$) | | | | | | |
| Protein | 172.93 | 47.47 | 148.55 | 35.36 | 35.31 | 72.88 |
| Ligand | 165.78 | 64.69 | 108.48 | 57.06 | | 27.00 |
| R.m.s. deviations | | | | | | |
| Bond lengths (Å) | 0.003 | 0.004 | 0.003 | 0.004 | 0.004 | 0.003 |
| Bond angles (°) | 0.617 | 0.666 | 0.650 | 0.743 | 0.668 | 0.517 |
| **Validation** | | | | | | |
| MolProbity score | 1.62 | 1.63 | 1.63 | 1.48 | 1.45 | 1.59 |
| Clashscore | 5.68 | 6.03 | 6.27 | 5.65 | 5.51 | 6.03 |
| Poor rotamers (%) | 0.33 | 0.76 | 0.04 | 0.76 | 0.19 | 0.00 |
| Ramachandran plot | | | | | | |
| Favored (%) | 95.59 | 95.64 | 95.88 | 97.04 | 97.19 | 96.19 |
| Allowed (%) | 4.41 | 4.36 | 4.12 | 2.96 | 2.81 | 3.81 |
| Disallowed (%) | 0.00 | 0.00 | 0.00 | 0.00 | 0.00 | 0.00 |
